# Mitochondrial Complex I and ROS control neuromuscular function through opposing pre- and postsynaptic mechanisms

**DOI:** 10.1101/2024.12.30.630694

**Authors:** Bhagaban Mallik, Sajad A. Bhat, Xinnan Wang, C. Andrew Frank

## Abstract

Neurons require high amounts energy, and mitochondria help to fulfill this requirement. Dysfunctional mitochondria trigger problems in various neuronal tasks. Using the *Drosophila* neuromuscular junction (NMJ) as a model synapse, we previously reported that Mitochondrial Complex I (MCI) subunits were required for maintaining NMJ function and growth. Here we report tissue-specific adaptations at the NMJ when MCI is depleted. In *Drosophila* motor neurons, MCI depletion causes profound cytological defects and increased mitochondrial reactive oxygen species (ROS). But instead of diminishing synapse function, neuronal ROS triggers a homeostatic signaling process that maintains normal NMJ excitation. We identify molecules mediating this compensatory response. MCI depletion in muscles also enhances local ROS. But high levels of muscle ROS cause destructive responses: synapse degeneration, mitochondrial fragmentation, and impaired neurotransmission. In humans, mutations affecting MCI subunits cause severe neurological and neuromuscular diseases. The tissue-level effects that we describe in the *Drosophila* system are potentially relevant to forms of mitochondrial pathogenesis.

## INTRODUCTION

Neurons have vast energy needs. These needs are primarily satisfied by healthy pools of mitochondria ^1,2^. Mitochondria generate energy through the action of the ATP synthase complex in the electron transport chain ^3,4^. They also perform complementary functions, including maintaining calcium homeostasis ^5,6^, promoting cell survival ^7^, triggering reactive oxygen species (ROS) signaling ^8^, stimulating lipid synthesis ^9^, and regulating innate immunity ^10^. For energy-driven neurons, it is thought that the primary role of mitochondria is to provide ATP. It is less understood how other mitochondrial functions contribute to the regulation of normal neurophysiology. It is also not well understood how neural tissues or synaptic sites cope when they are challenged with a loss of mitochondria. Genetic models can help to address these puzzles.

Mitochondrial Complex I (MCI) (NADH ubiquinone oxidoreductase) is an essential part of the electron transport chain and ATP production, and it comprises dozens of distinct subunits. Much of our understanding about MCI derives from systemic analyses of its assembly. Studies have been performed on Complex I components from diverse organisms, including *Neurospora crassa* and *Drosophila melanogaster*. Those studies demonstrate that discrete MCI subunits are ancient; indeed, there are few differences between these MCI models from simple organisms and the corresponding human and bovine orthologs ^11–14^. For *Drosophila melanogaster*, MCI consists of at least 42 distinct subunits; the 14 core MCI subunits are present, as are at least 28 accessory subunits ^12^.

In humans, MCI dysfunction has been linked to diseases such as Leigh syndrome, mitochondrial myopathy, and encephalomyopathy, as well as forms of stroke ^15–18^. On a cellular level, MCI dysfunction can cause the demise of neurons and muscles; these phenotypes are typically attributed to defects in ATP production ^19,20^. However, in addition to the ATP production defects, mutations affecting MCI subunit components are also associated with excess mitochondrial ROS. Normally, ROS accumulation can be neutralized by the cellular antioxidant system ^21^. But if that system becomes overwhelmed, there can be consequences for cells and organ systems – including progressive neurodegeneration and seizures for the nervous system ^22–24^. On the level of synapses, it is possible that MCI loss triggers severe molecular consequences, and it is also possible that excess ROS plays a role.

For a previous paper, we depleted MCI function at the *Drosophila* neuromuscular junction (NMJ). Our data suggested fundamental synaptic functions for MCI ^13^. Here we expand upon that work, mostly taking advantage of RNAi-mediated depletion of the *Drosophila* homologs of human NDUFS7. We also scrutinize loss-of-function mutants of other MCI subunits and pharmacological inhibition of MCI. Our collective data show that MCI depletion causes *Drosophila* phenotypes reminiscent of mitochondrial diseases, such as progressive degeneration of muscle and presynaptic cytoskeleton, excess ROS production, loss of mitochondria and alteration in mitochondrial morphology.

On single-tissue levels, we were surprised to find that there were opposite effects on synapse activity in the presynaptic motor neurons vs. the postsynaptic muscles. MCI dysfunction in *Drosophila* motor neurons causes profound cytological phenotypes, but there are no significant functional phenotypes. This appears to be because neuronal mitochondrial ROS triggers an adaptive response, demonstrated visually by active zone enhancement. This ROS-driven enhancement of active zones occurs through at least two processes, 1) regulation of calcium flux from intracellular stores (ER) and mitochondria; and 2) use of glycolysis as an alternative energy source. By contrast, postsynaptic depletion of MCI and the associated elevation of muscle ROS triggers a destructive response: disruption of NMJ morphology and the Dlg-Spectrin scaffold that is critical for normal active zone-receptor apposition. To our knowledge, these cellular and molecular mechanisms of MCI deficiency have not previously been elucidated at synapse-specific or tissue-specific levels.

## RESULTS

### One RNAi transgene targets two *Drosophila* homologs of a core MCI subunit, NDUFS7

We previously reported impairments in neuromuscular junction (NMJ) synapse development and function when Mitochondrial Complex I (MCI) was depleted ^13^. We observed robust synaptic phenotypes when driving a transgenic RNAi line against an MCI subunit, termed *UAS-ND-20L[RNAi]* (*ND-20L^HMJ237^*^77^) ^13^. That RNAi line encodes a short hairpin that matches 21 consecutive nucleotides of the *ND-20L* gene. Recent work indicates that *ND-20L* is only sparsely expressed in *Drosophila* tissues ^25–27^. However, the same short hairpin also matches 19 consecutive nucleotides of the *ND-20* gene, and *ND-20* is ubiquitously expressed ^25–27^. *ND-20* and *ND-20L* encode *Drosophila* homologs of the core NDUFS7 MCI subunit. Because both genes are effectively “on-target” for the RNAi transgene, we term the *UAS-ND-20L[RNAi]* transgene as *UAS-NDUFS7[RNAi]* for this study.

We used the GAL4/UAS system to verify that *UAS-NDUFS7[RNAi]* can diminish MCI function in relevant tissues. We drove its expression using neuronal or muscle GAL4 drivers. By quantitative RT-PCR, message levels of both NDUFS7-encoding genes were significantly diminished in those tissues in larvae (Fig. S1A-D). *ND-20L* message levels were down in both neurons and muscle, but variable in neurons, likely due to the low endogenous expression of *ND-20L* (Fig. S1A-B). *ND-20* message levels were significantly knocked down by the transgene in both tissues (Fig. S1C-D). We also tested if the transgene could diminish mitochondrial respiration by a Seahorse assay.

To acquire sufficient material for the assay, we collected mitochondria from adult muscles (see Methods). Driving the *UAS-NDUFS7[RNAi]* transgene in adult muscle starkly reduced oxygen consumption rate in muscle mitochondria across all timepoints tested (Fig. S1E-F). These assays confirmed that *UAS-NDUFS7[RNAi]* targets MCI.

### Depletion of MCI affects mitochondrial integrity in multiple *Drosophila* synaptic tissues

We wanted to understand what was happening to mitochondria to affect synapse function when MCI was depleted. We started by examining mitochondria by microscopy. To visualize them, we expressed a *UAS-Mito-GFP* transgene ^28^ in *Drosophila* tissues. Concurrently, we used tissuespecific GAL4 drivers alone (as controls) or GAL4 drivers + *UAS-NDUFS7[RNAi]* to deplete MCI function ^13^. With these tools, we made qualitative observations of mitochondrial morphology (Fig. 1), and we quantified those observations in subsequent analyses.

**Figure 1:**
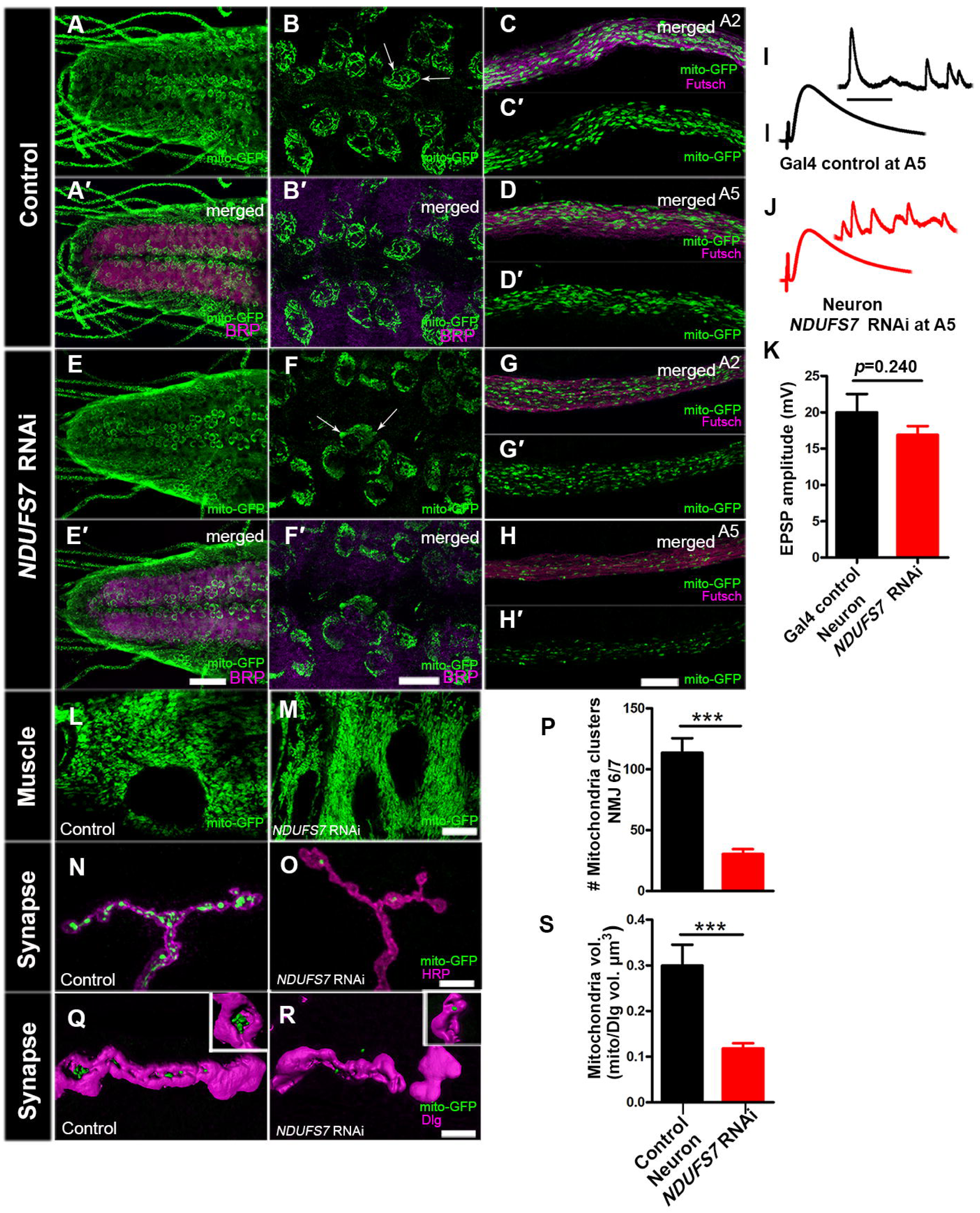
**MCI-depleted flies harbor fewer mitochondria at neuromuscular junctions** Mitochondrial morphology and trafficking defects in the ventral nerve cord, distal axons, and boutons. RNAi lines and controls were crossed to a motor neuron driver (*D42-GAL4*) and a mitochondrial marker (*UAS-mitoGFP*). (A-A′), (B-B′), (C-C′) and (D-D′) represent control ventral nerve cord (VNC), a magnified section of VNC, proximal (A2) and distal (A5) axons, respectively. These tissues exhibit regular mitochondrial clusters in the soma and axons. (E-E′), (F-F′), (G-G′) and (H-H′) represent *NDUFS7* knocked down ventral nerve cord (VNC), a magnified section of VNC, proximal (A2) and distal (A5) axons, respectively. Mitochondria are abnormally clustered in *NDUFS7[RNAi]* in the ventral nerve cord and distal segments of A5 axons. *NDUFS7[RNAi]* yields fewer mitochondria in the distal segments when compared to the proximal segments. (I-J) Representative electrophysiological traces showing evoked potentials of *mitoGFP, D42-Gal4 × UAS-NDUFS7*[RNAi] larvae at the A5 hemi segment of muscle 6/7 synapse. Scale bars for EPSPs (mEPSP) are x=50 ms (1000 ms) and y= 10 mV (1 mV). Fewer mitochondria at the presynaptic A5 hemi segment did not affect evoked NMJ excitation. K. Quantification showing EPSP amplitude at NMJ 6/7 in control (*D42-Gal4/+*; EPSP: 19.99 mV ± 2.53, n=6) and RNAi-depleted animals (*NDUFS7[RNAi] /+;D42-Gal4/+* ;EPSP: 16.88 mV ± 1.21, n=9). (L-M) Representative images showing mitochondria morphology in control and RNAi-depleted animals in muscle. Mitochondria in *NDUFS7[RNAi]*-depleted larvae are clustered compared to the control larvae. (N-Q) *NDUFS7[RNAi]* contains almost no mitochondria in boutons when co-stained with pre-(HRP) or post-synaptic markers (Discs Large, Dlg). (A-H′, L-M, N-O) Scale bar: 10 µm. (P) Quantification showing the number of mitochondrial clusters at NMJ 6/7 in control (*D42-Gal4/+*; # clusters: 113.6 ± 11.97, n=7) and RNAi-depleted animals (*NDUFS7[RNAi]/+; D42-Gal4/+*; # clusters: 30.33 ± 4.02, n=6). (Q-R) 3D rendered image showing the volume (µm^3^) of mitochondria at NMJ 6/7 in RNAi knockdown larvae (*NDUFS7[RNAi] /+; D42-Gal4/+ ;* 0.11 ± 0.01 µm^3^, n=16) compared to the driver control animals (*D42-Gal4/+*; 0.29 ± 0.04 µm^3^, n=14). (Q-R) Scale bar: 5 µm. (S) Quantification shows a significantly lower mitochondria volume in boutons at NMJ 6/7 in RNAi-depleted animals. \*\*\**p*<0.0001 and \*\*\**p*=0.0003 for mitochondrial clusters and volume, respectively. Statistical analyses based on Student’s t-test. Error bars represent mean ± s.e.m. Raw data for this figure are available in the S1 Data Excel file, tab Figure_1.

In motor neurons, the Mito-GFP signal localized to the neuropil of the ventral nerve cord (Fig.1: A-A′, B-B′). In control neurons, the neuropil mitochondria had a filamentous appearance. By contrast, NDUFS7-depleted neurons had punctate and clustered mitochondria. (Fig. 1E-E′, F-F′). We examined mitochondria in the motor axons that innervate proximal and distal NMJs (Fig. 1C-C′, D-D′, G-G’, H-H’). The proximal segment A2 axons had abundant mitochondria in all cases (Fig. 1C-C’, G-G’). However, for the distal segment A5 axons, NDUFS7 depletion elicited an obvious decrease in mitochondria number (Fig. 1D-D,′ H-H,′). This A2 vs. A5 discrepancy was consistent with prior work by others examining defects in mitochondrial trafficking dynamics: distal sites can show phenotypes more prominently ^29,30^.

We hypothesized that fewer mitochondria in the A5 axon might correlate with a neurotransmission defect at the NMJ. Yet by NMJ electrophysiology, we found no significant differences in the evoked amplitude compared to the control NMJs in the distal segment A5 (Fig. 1I-K). These data matched our prior examination of MCI at the A2 and A3 segments of the NMJ, where neuronal impairment of MCI was not sufficient on its own to reduce evoked NMJ neurotransmission ^13^.

In muscle, we observed an array of phenotypes. As with neurons, there were clustered mitochondria when *NDUFS7* gene function was depleted (Fig. 1L-M). Additionally, there was a tissue-level phenotype: *NDUFS7*-depleted muscles were developed, but they looked disorganized and fragmented, with oblong-shaped nuclei in the muscle syncytia (Fig. 1L-M). This phenotype could explain why we previously observed that muscle impairment of MCI was sufficient to reduce evoked NMJ neurotransmission ^13^.

To examine the mitochondria at presynaptic NMJ release sites, we used the motor neuron GAL4 driver to label NMJ boutons with Mito-GFP. For image analysis, we marked the presynaptic membrane boutons with anti-HRP immunostaining. Control NMJs contained abundant and large clusters of mitochondria in synaptic boutons, but by comparison, *NDUFS7[RNAi]* boutons contained small clusters and few mitochondria (Fig. 1N-P). We measured the mitochondrial volume in a 3-D stack and compared it to the synaptic volume (Fig. 1Q-S). The Mito-GFP signal occupied a sizeable proportion of the bouton volume in controls (∼30%), but this value was significantly diminished in *NDUFS7*-depleted animals (∼10%) (Fig. 1S). Collectively, our data suggest that the depletion of *NDUFS7* by RNAi leads to abnormal mitochondrial clustering in the neuronal cell body and muscle – as well as losses of distal axon and synaptic mitochondria.

### Loss of MCI phenocopies loss of Mitofusin

The cell-level NDUFS7-depletion phenotypes were reminiscent of *Drosophila* mutants impairing mitochondrial dynamics ^29,31^. Therefore, we re-examined MCI-depleted mitochondria, this time additionally impairing genes known to mediate mitochondrial fusion and fission. Mitofusin 1 (Mfn1) and Mitofusin 2 (Mfn2) are GTPases that regulate outer mitochondrial membrane fusion ^32,33^. The *Drosophila* gene encoding the Mitofusin homolog is called *marf*. Dynamin-related protein 1 is a GTPase that regulates mitochondrial fission. In *Drosophila*, this factor is encoded by the gene *drp1* ^34^. Previous work reported that defective fusion results in fragmented mitochondria, while defective fission can lead to enlarged mitochondria ^35^. We used RNAi-mediated knockdown constructs for each of these genes.

As before, we observed that wild-type motor neurons had filamentous and oval mitochondria, while NDUFS7-depleted neurons had fewer and smaller clustered mitochondria in the ventral nerve cord (VNC) (Fig. 2A) and axons (Fig. 2B). Knockdown of the fusion gene *marf* phenocopied *NDUFS7* loss, revealing small mitochondria in motor neurons, while knockdown of the fission gene *drp1* yielded filamentous mitochondria (Fig. 2A). Simultaneously depleting motor neurons of *marf* and *NDUFS7* by RNAi did not show any additive defect in mitochondrial appearance in the ventral nerve cord (VNC) and axons (Figs. 2A, B). This result could mean that the genes share a common process to regulate mitochondrial fusion. By contrast, depleting *drp1* and *NDUFS7* by RNAi simultaneously yielded punctate mitochondria. This result likely means that that the punctate *NDUFS7* mitochondrial phenotypes (potential fusion phenotypes) are epistatic to *drp1* loss (Fig. 2A, B).

**Figure 2:**
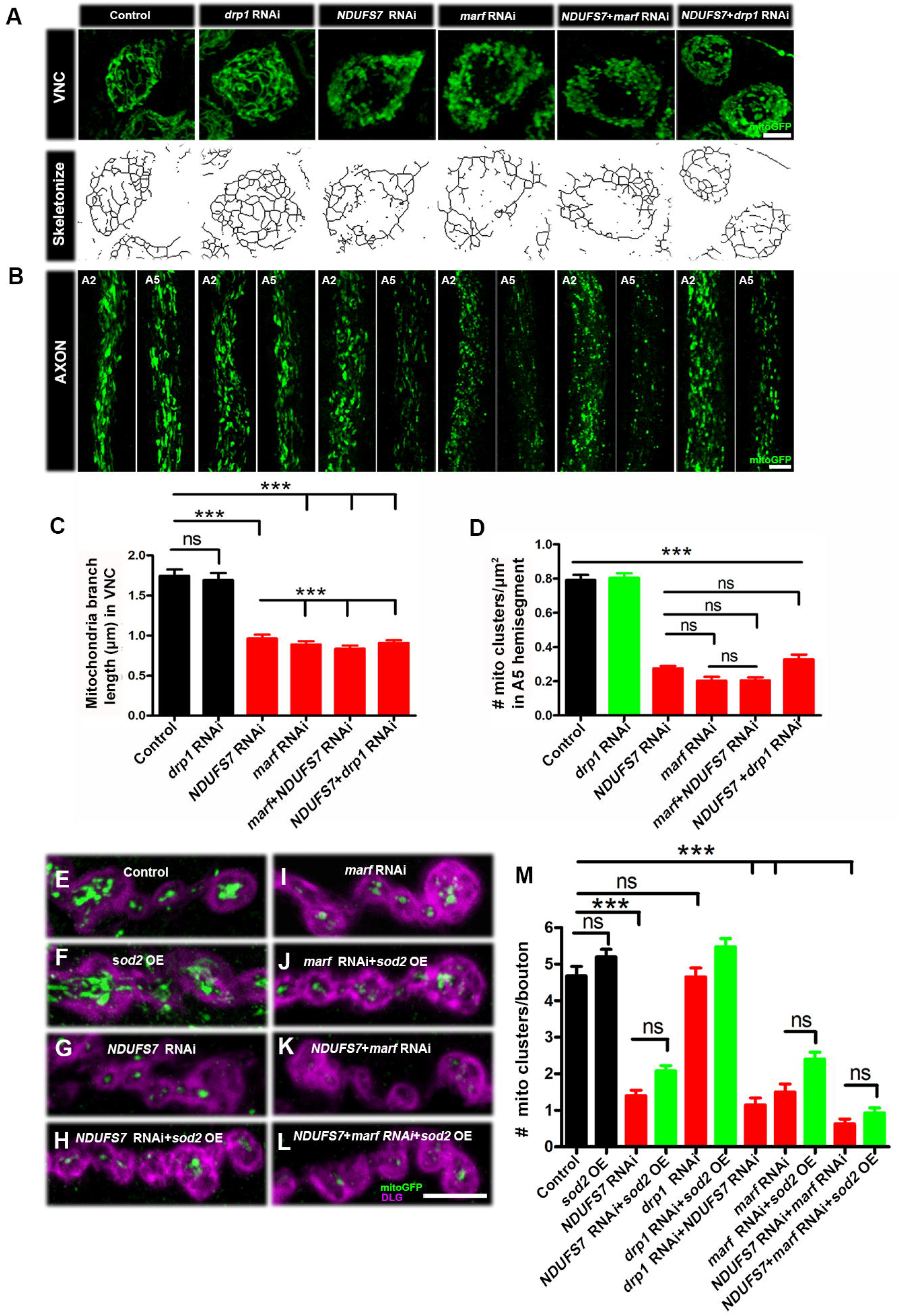
**Loss of *NDUFS7* in motor neurons phenocopies a *marf* depletion** Mitochondrial morphology and trafficking defects in the ventral nerve cord and distal axons. To label neuronal mitochondria, *UAS-[RNAi]* lines and controls were crossed to a motor neuron driver (*D42-Gal4*) and a mitochondrial marker (*UAS-mitoGFP*). (A) Ventral nerve cord (VNC): *UAS-mitoGFP* and *drp1[RNAi]* exhibit normal mitochondrial organization, *NDUFS7[RNAi]* and *marf*[RNAi] exhibit clustered mitochondria, *NDUFS7[RNAi] ; marf[RNAi]* and *ND-20[RNAi]; drp1[RNAi]* doubles exhibit clustered mitochondria in the soma. The fluorescent images were skeletonized to measure mitochondrial branch length. (B) Comparison of a proximal axonal segment in A2 and a distal segment in A5. Distal segments of A5 axons in *NDUFS7[RNAi]* and *marf[RNAi]* contain many fewer mitochondria than proximal segments. Knocking down *NDUFS7[RNAi]* and *marf[RNAi]* together does not show an additive effect. (A-B) Scale bar: 10 µm. (C-D) Histogram showing mitochondrial branch length (µm) and number (µm^2^ area of bouton) in VNC and axons of the third instar larvae in the indicated genotypes. (E-L) Representative images of the A2 hemi segment of muscle 6/7 NMJs in (E) *UAS-mito-GFP, D42-Gal4/+*, (F) *UAS-Sod2/+; UAS-mito-GFP, D42-Gal4/+,* (G) *NDUFS7[RNAi] /+; UAS-mito-GFP, D42-Gal4/+*, (H) *NDUFS7[RNAi] /UAS-Sod2;UAS-mito-GFP,D42-Gal4/+*, (I) *UAS-mito-GFP,D42-Gal4/marf[RNAi]*, (J) *UAS-Sod2;UAS-mito-GFP,D42-Gal4/marf[RNAi]*, (K) *NDUFS7[RNAi]/+; UAS-mito-GFP,D42-Gal4/marf[RNAi]* and (L) *NDUFS7[RNAi] /UAS-Sod2;UAS-mito-GFP,D42-Gal4/marf[RNAi]* larvae immunostained with antibodies against HRP (magenta) and GFP (mito-GFP:green) to label neurons and mitochondria. *NDUFS7[RNAi]-* and *marf[RNAi]*-depleted animals harbor fewer mitochondria at the terminals as compared to control animals. Transgenic *UAS-Sod2* rescued mitochondrial clustering defects. (E-L) Scale bar: 5 µm. (M) Histograms showing quantification of mitochondrial clusters at the NMJs in the indicated genotypes. \*\*\**p* <0.0001; ns, not significant. Statistical analysis based on one-way ANOVA followed by post-hoc Tukey’s multiple-comparison test. Error bars represent mean ± s.e.m. Raw data for this figure are available in the S1 Data Excel file, tab Figure_2.

We measured mitochondrial branch length from skeletonized images of the mitochondria (Skeletonize3D, ImageJ plugin, data in S1 Table, S2 Table and S3 Table). Control and *drp1* knock-down showed normal mitochondrial branch length, but knockdown using *NDUFS7[RNAi]* or *marf[RNAi]* – or knockdowns using combinations of each – exhibited short branch length (Fig. 2C, S1 Table).

To quantify mitochondria in axons, we counted Mito-GFP positive puncta in distal A5 motor axons labeled by anti-GFP. Control axons and *drp1-*depleted axons contained abundant mitochondria (Fig. 2D, S1 Table). By contrast, any gene manipulation or combination targeting *NDUFS7* or *marf* by RNAi resulted in diminished numbers of mitochondrial clusters (Fig. 2D, S1 Table).

We extended the analysis to NMJ terminals. We counted Mito-GFP clusters in presynaptic boutons apposed by postsynaptic densities, labeled by anti-Discs Large 1 (Dlg1) (Figs. 2E-M, S4 Table). The results matched our earlier observations (Figs. 1N-S). Control NMJs and *drp1*-depleted NMJs contained numerous mitochondrial clusters per bouton (Figs. 2E, M). However, NDUFS7*-* depleted boutons contained few mitochondria, and this was phenocopied by *marf[RNAi]* (Fig. 2E-M, S4 Table). Collectively, these results suggest that *Drosophila* NDUFS7 (and hence MCI) contributes to normal mitochondrial fusion, likely in conjunction with the Mitofusin homolog, Marf.

### Mitochondrial Reactive Oxygen Species Contribute to Synaptic Phenotypes

Several studies have demonstrated that Complex I loss results in high levels of mitochondrial reactive oxygen species (ROS) ^36–40^. This means that excess ROS could be contributing to the cytological and mitochondrial fusion phenotypes that we have described.

We checked if we could observe mitochondrial ROS (superoxide) in living *Drosophila* tissue and if ROS levels corresponded to Complex I function (Fig. S1). We used a commercially available fluorescent mitochondrial superoxide indicator, MitoSOX (ThermoFisher, Methods) ^41–43^. With MitoSOX, we observed mitochondrial superoxide in many tissues. There was a baseline level of ROS in controls (Fig. S2A, E, I, J, S5 Table), and the level was greatly increased in NDUFS7-deficient motor neuron cell bodies and muscle (Fig. S2B, F, I; J, S5 Table).

Next, we tested if ROS scavengers could reverse the high MitoSOX fluorescence levels in *NDUFS7*-deficient tissues. We fed a pharmacological scavenger, N-Acetyl Cysteine Amide (NACA) ^44–46^ to *Drosophila* larvae (Methods). We also used a transgene, *UAS-Sod2* ^47^, to express a superoxide dismutase enzyme. Both successfully diminished the high levels of mitochondrial ROS that resulted from *NDUFS7* depletion at the NMJ, and both worked in muscle and neurons (Fig. S2C, D, G-J, S5 Table). We also tested a related idea: if genetic *Sod2* knockdown by RNAi could phenocopy loss of MCI by enhancing Mito-SOX levels in these same tissues. It did. Transgenic *UAS-Sod2[RNAi]* enhanced Mito-SOX signal in all tissues and sub compartments, including the ventral nerve cord (Fig. S3A-C, S6 Table), the motor neuron axons (Fig. S3D-F, S6 Table), the synaptic boutons (Fig. S3G-I, S6 Table), and the muscle (Fig. S3J-L, S6 Table).

Next, we tested if these same ROS scavengers could reverse mitochondrial phenotypes caused by loss of MCI. Co-expressing *UAS-Sod2* or rearing larvae with NACA suppressed the mitochondrial morphology defects in the ventral nerve cord of *NDUFS7*-depleted animals; it also restored axonal loss of mitochondria (Fig S4A-D, S2 Table). To test an additional MCI manipulation, we knocked down *Drosophila ND-30* (homologous to human *NDUFS3*) in motor neurons. As with *NDUFS7*, depleting *ND-30* in motor neurons yielded punctate mitochondria in the ventral nerve cord, but the addition of *UAS-Sod2* restored a wild-type, filamentous mitochondrial morphology (Fig. S4A, C). Similarly, loss of *ND-30* gene function in neurons depleted A5 axonal mitochondria and this phenotype was also reversed by *UAS-Sod2* transgenic expression (Fig. S4B, D, S2 Table). But in contrast to *UAS-Sod2*, neither *UAS-Sod1* nor *UAS-Catalase* worked to reverse mitochondrial morphology defects (Fig. S5, S3 Table). SOD1 (cytosol) and Catalase (peroxisome) localize to different compartments than SOD2 (mitochondrial matrix). Therefore, these results could indicate that a scavenger needs to access the proper compartment for rescue.

Because of the links between MCI and mitochondrial fusion, we considered whether a *marf* loss of function could also yield high levels of neuronal ROS (Fig. S6). It did – both *marf* and *NDUFS7* loss-of-function conditions showed high levels of mitochondrial superoxide in motor neuron cell bodies (Fig. S6A-D, A’-D’, M, S5 Table) in motor axons (Fig. S6E-H, E’-H’, N, S5 Table); and at NMJ sites (Fig. S6I-L, I’-L’, O, S5 Table). These ROS phenotypes were not confined to genetic manipulations. We made similar observations when MCI was impaired pharmacologically by feeding rotenone to developing larvae (Fig. S6, S5 Table). In the case of rotenone, the amount of mitochondrial ROS in the tissues was high, but it was generally not increased as much as with the genetic manipulations (Fig. S6M-O, S5 Table).

Finally, because the distal A5 motor axons accumulated high levels of ROS when subjected to these insults, we examined them for synaptic vesicle trafficking defects. We immunostained for Cysteine String Protein (CSP), a DNAJ-like co-chaperone and synaptic vesicle-associated protein (Cysteine string protein: CSP). *NDUFS7* depletion caused aberrant accumulation of CSP in the A5 motor axons; and this defect was suppressed by motor neuron transgene overexpression of *UAS-Sod2* or by feeding animals with NACA (Fig. S7A-L, S7 Table). However, this phenotype was not suppressed by motor neuron overexpression of *UAS-Sod1* or *UAS-Catalase* (Fig. S8, S8 Table).

ROS scavengers did not reverse all mitochondrial abnormalities. Expressing *UAS-Sod2* in the *UAS-NDUFS7[RNAi]* or *UAS-marf[RNAi]*-depletion backgrounds did not restore mitochondrial clusters to motor neuron terminals (Fig. 2E-M, Table 2). For the remainder of the study, we used scavengers as complementary tools to test which MCI-loss phenotypes were likely due to mito-chondrial ROS.

### Loss of MCI subunits impairs synaptic cytoskeletal stability

ROS can modulate the cytoskeleton, either through redox modification of cytoskeletal proteins or by altering pathways that regulate cytoskeletal organization ^48^. To test whether the mitochondrial defects and abnormal accumulation of ROS were associated with the altered synaptic cytoskeleton, we labeled synaptic boutons with an anti-Futsch antibody (Fig. 3). Futsch is a *Drosophila* MAP1B homolog that associates with microtubules ^49^.

**Figure 3:**
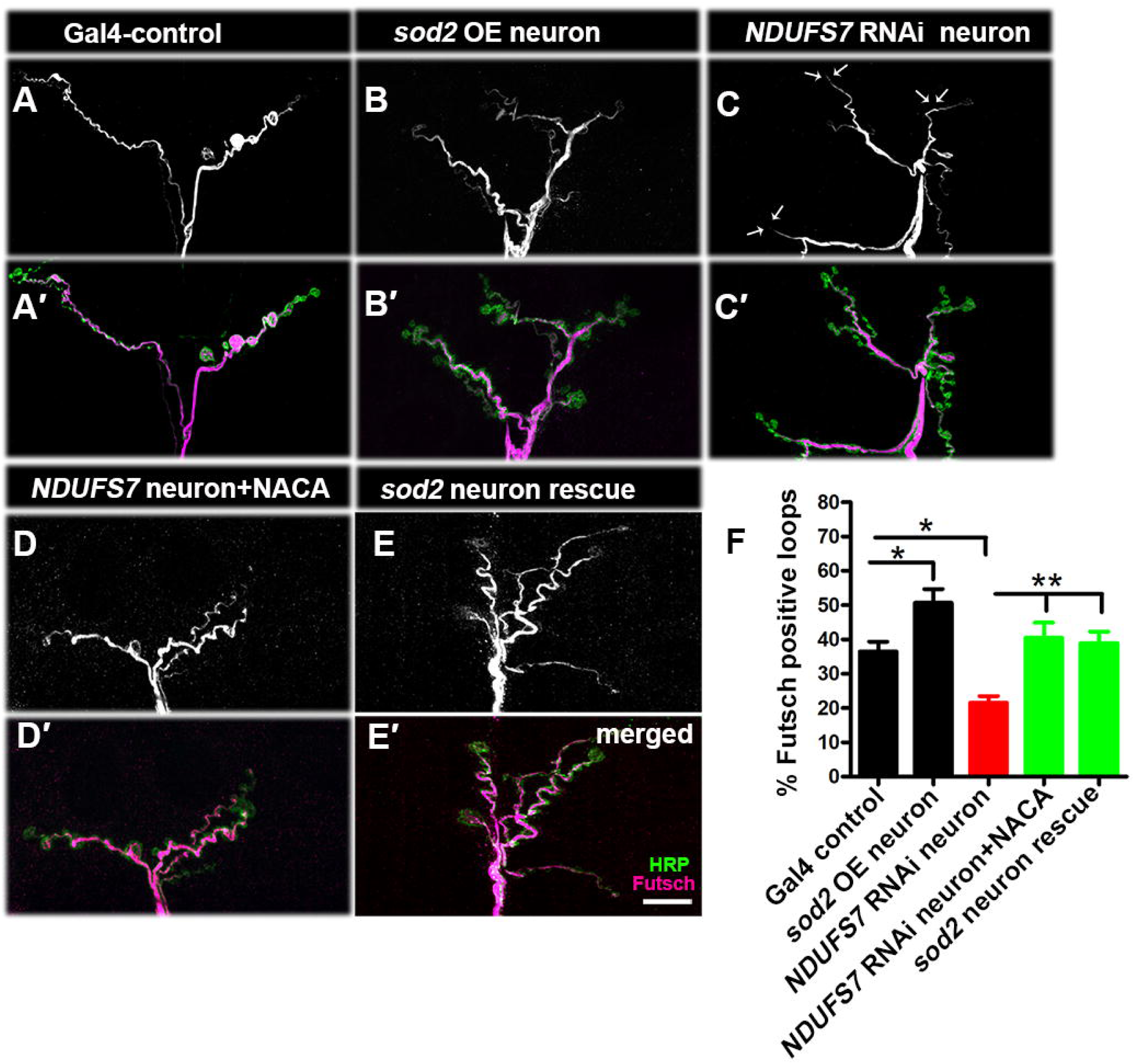
***NDUFS7* depletion in motor neurons affects synapse stability** Representative confocal images of NMJ synapses at muscle 6/7 of (A-A′) *D42-Gal4* control, (B-B′) *UAS-Sod2* overexpression (C-C′) *D42*-*Gal4*-driven *NDUFS7*[RNAi] (*NDUFS7[RNAi] /+; D42-Gal4/+*), (D-D′) *NDUFS7* knockdown with NACA rescue (*NDUFS7[RNAi] /+; D42-Gal4/+* +NACA), (E-E′) *NDUFS7* knockdown with *UAS-Sod2* (*UAS-NDUFS7[RNAi]/UAS-Sod2; D42-Gal4/+*). Each condition was double immunolabeled with 22C10 (anti-Futsch, magenta) and anti-HRP (green) antibodies. The motor neuron-depleted *NDUFS7[RNAi]* larvae showed a decrease in the number of Futsch-positive loops as compared to the Gal4 control. Futsch-positive loops were significantly restored to the control number when *NDUFS7*[RNAi] was raised in media containing NACA or genetically expressing *UAS-Sod2* in the *UAS-NDUFS7[RNAi]* background. Scale bar: 10 µm. (F) Histograms showing the percent-age of Futsch-positive loops in the indicated genotypes. \**p*=0.01 (Gal4 control vs *sod2* OE neuron), \**p*=0.0008 (Gal4 control vs *NDUFS7*[RNAi] neuron), \*\**p*=0.001 (*NDUFS7*[RNAi] neuron vs *NDUFS7*[RNAi] neuron + NACA) and \*\**p*=0.0005 (*NDUFS7*[RNAi] neuron vs *sod2* neuron rescue). Statistical analysis based on one-way ANOVA followed by post-hoc Tukey’s multiple-comparison test. Error bars represent mean ± s.e.m. Raw data for this figure are available in the S1 Data Excel file, tab Figure_3.

In motor neuron Gal4-control and *UAS-Sod2* overexpression larvae, Futsch organized in periodic loops, as expected from previous characterizations ^49^ (Fig. 3A-B, S7 Table). But in NDUFS7-depleted larvae, the anti-Futsch staining showed a significant reduction in microtubule loops (Figure 3C, S7 Table). These *NDUFS7* phenotypes were suppressed by motor neuron overexpression of *UAS-Sod2* or by raising animals on food containing 0.5 mM NACA (Fig. 3D-E, S7 Table). However, they were not significantly restored by neuronal overexpression of *UAS-Sod1* or *UAS-Catalase* (Fig. S9, S8 Table). These data indicate that loss of MCI regulates cytoskeletal architecture due to excessive accumulation of mitochondrial ROS in neurons.

### Modest neurotransmission phenotypes after motor neuron-specific loss of MCI or Marf

Given the cytological phenotypes after neuronal MCI loss, it was puzzling that there seemed to be little-to-no electrophysiological consequence at NMJs (^13^ and Fig. 1J). We probed this finding, this time depleting motor neurons of *marf* and/or *NDUFS7* gene function. We recorded spontaneous miniature postsynaptic potentials (mEPSP) and evoked excitatory postsynaptic potentials (EPSP).

Phenotypes were normal-to-mild (Fig. S10). For both *NDUFS7[RNAi]* and *marf[RNAi]*, there were small, but statistically significant decreases in mEPSP amplitude (Fig. S10A-C, G, S9 Table). But for *NDUFS7[RNAi]*, evoked events (EPSP) and calculated quantal content (QC) were at control levels (Fig. S10A-B, H-I, S9 Table). For neuronal *marf[RNAi]*, those measures were near-normal, with a slight decrease in EPSP amplitude (Fig. S10C, H, S9 Table) and a slight increase in QC (Fig. S10I, S9 Table). Finally, for a double *NDUFS7[RNAi]* + *marf[RNAi]* knockdown condition in neurons, we observed a small, but statistically significant decrease in EPSP amplitude (Fig. S10E, H, S9 Table).

Synaptic phenotypes caused by mitochondrial dysfunction might be masked until synapses are challenged with extreme conditions, like high frequency stimulation ^50,51^. Therefore, we challenged neuronally *NDUFS7*-depleted *Drosophila* NMJs in several ways (Fig. S11). First, we lowered recording saline [Ca^2+^] to 0.15 mM, which is roughly one order of magnitude lower than physiological calcium. In low calcium, the motor neuron-driven *NDUFS7[RNAi]* NMJs had slightly smaller evoked potentials compared to driver controls, but the numerical reduction was not statistically significant (Fig. S11A-C, S10 Table). Next, we lowered extracellular [Ca^2+^] even further, to 0.1 mM, which yielded a mix of successful EPSP firing events and failures. Failure analyses revealed an increase in failure rate at the neuronally depleted *NDUFS7[RNAi]* NMJs compared to control animals (Fig S11D, S10 Table), demonstrating a decreased probability of release. Failure rates in 0.1 mM calcium were restored to baseline levels when *NDUFS7*-depleted larvae were raised in a media containing NACA or genetically expressing *Sod2* in motor neurons (Fig S11D, S10 Table), indicating that a sensitivity to low calcium could be related to ROS levels.

Finally, we checked if forms of short-term neuroplasticity were affected by *NDUFS7* loss in motor neurons. For two different extracellular [Ca^2+^] conditions (0.4 to 1.5 mM), we did not observe any significant changes in paired-pulse ratios (Figs. S11E-J, S10 Table). Likewise, we did not note depreciation of evoked neurotransmission over the course of high frequency stimulus trains in high calcium (Figs. S11K-P). Collectively, these data suggest that there might be a small effect on NMJ physiology due to defective mitochondrial fusion, and the defect could be sensitive to low levels of extracellular calcium – but the aggregate data also indicate that neuronal mitochondrial defects alone do not drastically affect NMJ neurotransmission.

### Loss of MCI in neurons controls the level and distribution of the active zone to stabilize synaptic strength

We wondered how the NMJ synapse might evade severe dysfunction, despite loss of mitochondria in the motor neurons. One possibility is that *NDUFS7* loss/MCI impairment could trigger a form of functional homeostatic compensation of the NMJ. Another idea is that the mitochondrial ATP generated is superfluous at the NMJ – and that any energy-intensive functions that mitochondria support could be redundantly covered by glycolysis. These models are not mutually exclusive, and for any scenario, mitochondrial ROS downstream of defective MCI could be a candidate signal. Recent findings have demonstrated that ROS intermediates, mitochondrial distribution, and mitochondrial trafficking all affect development of the *Drosophila* NMJ ^48,52,53^.

We imaged the presynaptic active zone apparatus in neuronally depleted *NDUFS7[RNAi]* flies. Third-instar larval active zones showed a decrease in BRP puncta density per unit area in *NDUFS7*-depeleted NMJs compared to control NMJs (Fig.4A-C, G, H-I, S11 Table). But they also showed a robust enhancement phenotype: in *NDUFS7*-depleted animals, we found a 40% increase in active zone (BRP) immunofluorescence signal per unit area, compared to control (Fig. 4A-C, F, H-I, S11 Table), by laser scanning confocal microscopy. This result was intriguing because NMJ active zone enhancements (or changes in active zone sub-structure) have been proposed by other labs to be molecular correlates of forms of homeostatic plasticity and potentiation of neurotransmitter release ^54–58^. This raised the possibility that the NMJs evade severe dysfunction through a form of synaptic homeostasis.

**Figure 4:**
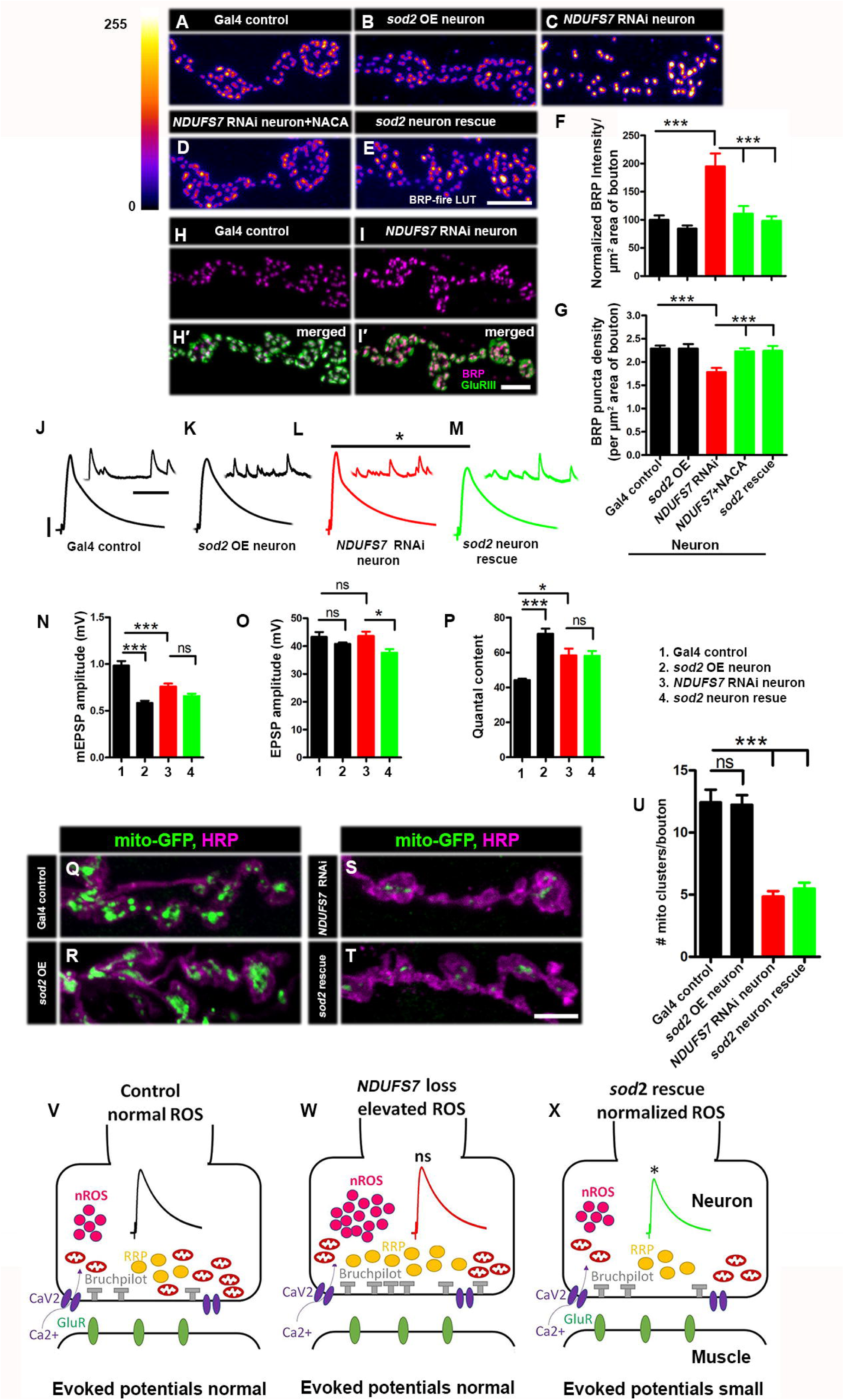
**Neuronal ROS (nROS) controls active zone material levels at NMJs** (A) Representative images of the A2 hemi segment of muscle 6/7 NMJs in *UAS-mito-GFP, D42-Gal4/+*, (B) *UAS-Sod2/+; UAS-mito-GFP, D42-Gal4/+* (C) *NDUFS7[RNAi]/+; UAS-mito-GFP, D42-Gal4/+*, (D) *NDUFS7[RNAi] /+; UAS-mito-GFP, D42-Gal4/+* with NACA and (E) *NDUFS7[RNAi]/UAS-Sod2; UAS-mito-GFP, D42-Gal4/+* larvae immunostained with antibodies against the active zone scaffold Bruchpilot (BRP:fire-LuT) to label the active zones. BRP levels are upregulated at the NMJs in *NDUFS7*[RNAi] depleted flies, while overexpression of ROS scavenger gene *Sod2* in the neuron or feeding the larvae with N-Acetyl L-cysteine amide (NACA) restores BRP to the control level. (A-E) Scale bar: 2.5 µm. (F-G) Histograms showing quantification of BRP intensity (F) and density (G) in µm^2^ area of bouton at muscle 6/7 in the genotypes mentioned above. At least 8 NMJs of each genotype were used for quantification. ***p <0.0001. Error bars denote mean ± s.e.m. Statistical analysis based on one-way ANOVA followed by post-hoc Tukey’s multiple-comparison test (H-H′-I-I′). Representative confocal images of muscle 6/7 NMJs in the (H-H′) control (*D42-Gal4/+*) and (I-I′) motor neuron Gal4 driven *NDUFS7[RNAi]* (*NDUFS7[RNAi]/+; D42-Gal4/+*) immunostained with antibodies against Bruchpilot (BRP: magenta) and GluRIII (green) to label a glutamate receptor subunit. (H-H′-I-I′) Scale bar: 2.5 µm. There are no significant changes of GluRIII-BRP apposed clusters. At least 8 NMJs of each genotype were used for quantification (J-P). Representative traces, quantification of mEPSPs, EPSPs and quantal content in the indicated genotypes. Scale bars for EPSPs (mEPSP) are x=50 ms (1000 ms) and y= 10 mV (1 mV). EPSPs amplitudes were maintained in *NDUFS7[RNAi]-*depleted flies due to increased levels of active zone material (e.g., BRP); however, NMJs with *Sod2-*rescued *NDUFS7[RNAi]* in neurons showed diminished evoked release when compared with *NDUFS7[RNAi]*. Minimum 8 NMJs recordings of each genotype were used for quantification. \**p* < 0.05, \*\*\**p* <0.0001; ns, not significant. Statistical analysis based on one-way ANOVA followed by post-hoc Tukey’s multiple-comparison test. Error bars denote the standard error of the mean. (Q) Representative images of the A2 hemi segment of muscle 6/7 NMJs in *UAS-mito-GFP, D42-Gal4/+*, (R) *UAS-Sod2/+; UAS-mito-GFP, D42-Gal4/+* (S) *NDUFS7[RNAi]/+; UAS-mito-GFP, D42-Gal4/+*, and (T) *NDUFS7[RNAi]/UAS-Sod2; UAS-mito-GFP,D42-Gal4/+* larvae immunostained with antibodies against HRP (magenta) and GFP (mito-GFP:green) to label neurons and mitochondria. *NDUFS7-*depleted and *sod2*-rescued *NDUFS7*[RNAi] animals harbor fewer mitochondria at the terminals than control animals. (Q-T) Scale bar: 5 µm. (U) Histograms showing quantification of mitochondrial clusters in the above-indicated genotypes. (V-X) Schematic illustration (drawn on bioRender) showing ROS (magenta) levels, BRP (grey) and mitochondria (red) number in the indicated genotypes. At least 8 NMJs of each genotype were used for quantification. \*\*\**p*<0.0001. Error bars represent mean ± s.e.m. Statistical analysis based on one-way ANOVA followed by post-hoc Tukey’s multiple-comparison test. Raw data for this figure are available in the S1 Data Excel file, tab Figure_4.

As an independent test, we impaired MCI pharmacologically. To do this, we raised larvae on 50 µM rotenone-spiked food; and we also incubated wild-type fillet preparations with 500 µM of rotenone for extended time. For both cases, we observed significant increases in BRP protein at the presynaptic active zones (Fig S12A-U, S12 Table). For the extended incubation, the fillet preparations required sufficient rotenone incubation time (six hours) and an intact motor nerve to show the active zone enhancement (Fig. S12U, S12 Table). This result suggested that downstream of MCI depletion, a compensatory delivery of active zone material required either substantial trafficking time and/or fully intact neuroanatomy.

Next, we checked if the enhanced active zone signal by was triggered by excess mitochondrial ROS in motor neurons. Indeed, we found that the *NDUFS7*-depletion active zone enhancements were fully reversed by ROS scavengers, either by raising larvae in food containing NACA or by neuronally expressing *UAS-Sod2* (Fig. 4D-G, S11 Table). However, they were not reversed by overexpressing *UAS-Sod1* or *UAS-Catalase* (Fig. S13A-G, S13 Table).

Finally, we assessed synapse function. As with our prior recordings, evoked postsynaptic potentials at the NMJ were not significantly changed by *NDUFS7* depletion in motor neurons. But interestingly, scavenging mitochondrial ROS in the *NDUFS7[RNAi]* neuronal depletion background with *UAS-Sod2* unmasked a small deficit in NMJ excitation, compared to controls (Fig. 4J-P, S11 Table). As before, neither *UAS-Sod1* overexpression nor *UAS-Catalase* overexpression unmasked this deficit in NMJ excitation (Fig. S13N-P, S13 Table).

These data could mean that mitochondrial ROS is helping to maintain synaptic activity. Neuronal expression of *UAS-Sod2* did not restore mitochondrial clusters to the NMJ after *NDUFS7* gene function depletion (Fig. 4Q-U, S11 Table; like Fig. 2H), meaning that the synaptic sites were still deficient in mitochondria. And as expected, neuronal expression of *UAS-Sod1* and *UAS-Catalase* also failed to mitochondrial clusters to the NMJ (Fig. S13Q-W, S13 Table). Collectively, our data support a model in which neuronal ROS (nROS) triggers active zone enhancement and functional compensation when MCI is limiting (Fig. 4V-X, S11 Table).

### Neuronal MCI subunits stabilize synaptic strength in conjunction with intracellular calcium signaling proteins

Recent work described a mechanism for local calcium uptake into mitochondria that drives ATP production to maintain synaptic function ^59^. The process is governed by the mitochondrial calcium uniporter (MCU) and its accessory EF-hand MICU proteins ^59^. Beyond this role for mitochondrial calcium, there are also known roles for core synaptic functions like vesicle cycling ^60,61^.

To test if mitochondrial or neuronal calcium could be involved in maintaining synapse function at the NMJ, we acquired genetic reagents to examine depressed MCU function in conjunction with depressed MCI. We also used pharmacological reagents to inhibit release of intracellular sources of calcium, like those from the Ryanodine Receptor (RyR) and the IP_3_ receptor (IP_3_R) of the endoplasmic reticulum (ER) ^62^, as well as a genetic reagent that we previously used at the NMJ to deplete IP_3_ signaling (*UAS-IP_3_-sponge*) (Fig. 5A-D). For this set of experiments, we used the *NDUFS7* neuronal knockdown condition as a sensitized genetic background (Fig. 5E-T, S14 Table). We also used a pan-neuronal driver (*elaV-Gal4*) for ease *Drosophila* genetic stock construction instead of a motor neuronal driver. We previously observed similar NMJ effects either for motor neuronal or pan-neuronal depletion of MCI ^13^.

**Figure 5:**
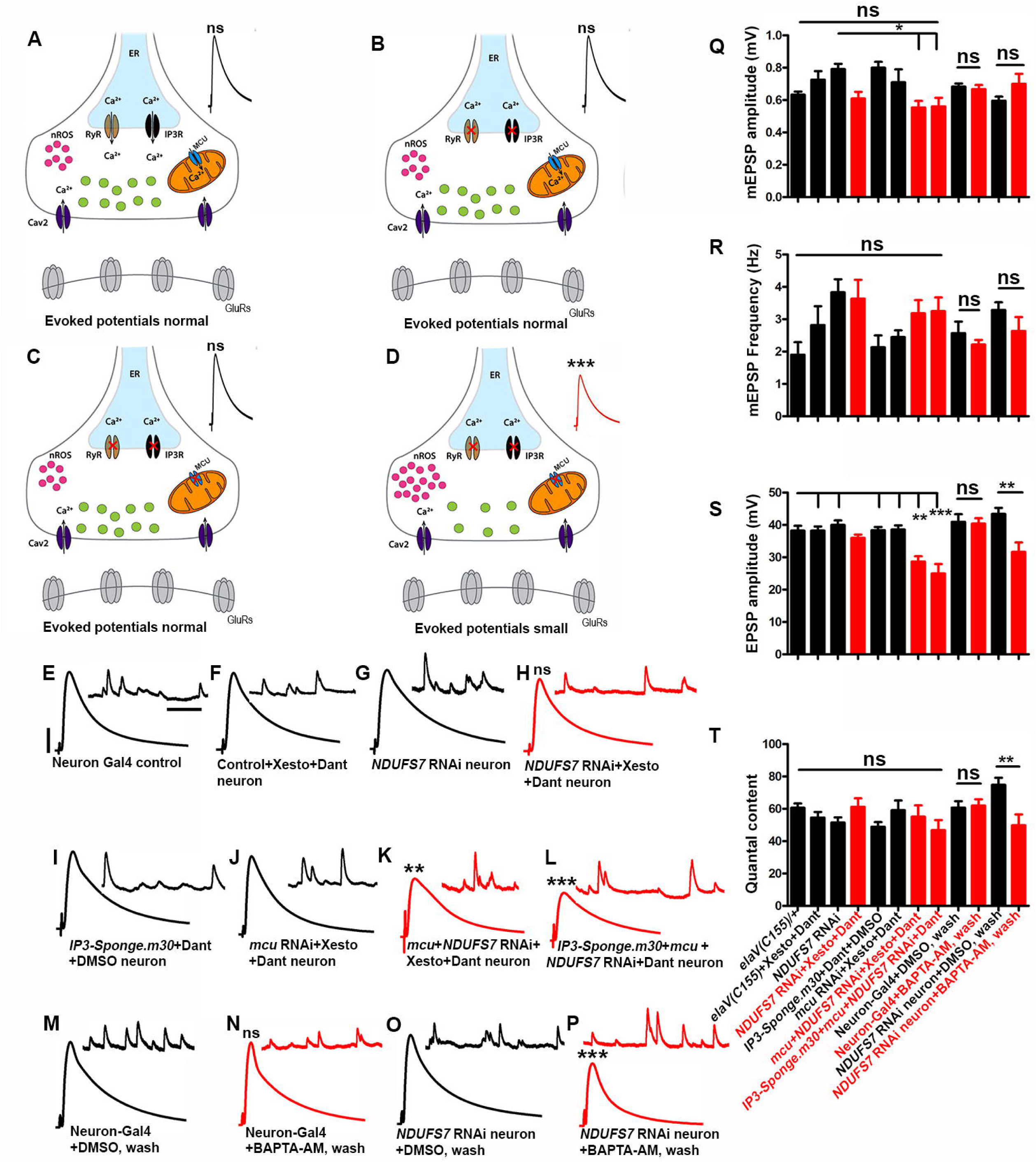
Loss of an MCI subunit necessitates ER-mediated calcium release to maintain evoked neurotransmission at the NMJs. (A-D) Schematics (drawn on Adobe Illustrator) illustrating the role of IP_3_ receptor (IP_3_R), Ryanodine receptor (RyR) in the endoplasmic reticulum, mitochondrial calcium uniporter Complex (MCU), Ca_v_2 calcium channel and synaptic vesicles at the presynaptic nerve terminal. The IP_3_ and Ryanodine receptors were blocked pharmacologically (red X symbols) by using the IP_3_R antagonist Xestospongin C or neuronally expressing *UAS-IP_3_-Sponge* and RyR antagonist Dantrolene, while *mcu[RNAi]* was used to block the Mitochondrial calcium uniporter complex (red X). (E-P) Representative traces of mEPSPs and EPSPs in (E) pan-neuronal Gal4 control (*elaV(C155)-Gal4/+*), (F) pan-neuronal Gal4 control (*elaV(C155)-Gal4/+*) with an acute application (10 minutes) of 20 µM Xestospongin C and 10 µM Dantrolene, (G) pan-neuronal Gal4 driven *NDUFS7*[RNAi] (*elaV(C155)-Gal4)/+;NDUFS7[RNAi] /+*), (H) pan-neuronal Gal4-driven *NDUFS7*[RNAi] (*elaV(C155)-Gal4/+;NDUFS7[RNAi] /+*) with 20 µM Xestospongin C and 10 µM Dantrolene, (I) pan-neuronal Gal4 driven UAS-*IP3-Sponge.m30* with an acute application of 10 µM Dantrolene (*elaV(C155)-Gal4*)*/+*; *NDUFS7[RNAi]/+ ;UAS-IP3-Sponge.m30/+*), (J) pan neuronal Gal4 driven *mcu*[RNAi] with 20 µM Xestospongin C and 10 µM Dantrolene (*elaV(C155)-Gal4)/+; mcu[RNAi]/+),* (K) pan-neuronal *mcu[RNAi] + NDUFS7*[RNAi] with 20 µM Xestospongin C and 10 µM Dantrolene and(*elaV(C155)-Gal4)/+; mcu[RNAi]/NDUFS7[RNAi]*) and (L) pan neuronal *UAS-IP3-Sponge.m30* + *mcu[RNAi] + NDUFS7*[RNAi] with 10 µM Dantrolene (*elaV(C155)-Gal4/+;NDUFS7[RNAi] /mcu[RNAi] ;UAS-IP_3_-Sponge.m30*/+), (M) Pan neuronal Gal4 (*elaV(C155)-Gal4/+*) with DMSO, (N) pan neuronal Gal4 (*elaV(C155)-Gal4/+*) with 20 µM BAPTA-AM, (O) pan-neuronal Gal4 driven *NDUFS7*[RNAi] (*elaV(C155)-Gal4)/+;NDUFS7[RNAi] /+*) with DMSO and (P) pan neuronal Gal4 driven *NDUFS7*[RNAi] (*elaV(C155)-Gal4)/+;NDUFS7[RNAi] /+*) with 20 µM BAPTA-AM. Scale bars for EPSPs (mEPSP) are x=50 ms (1000 ms) and y= 10 mV (1 mV). Note that EPSPs amplitudes were reduced in pan-neuronal Gal4 driven *mcu[RNAi] + NDUFS7*[RNAi] with an acute exposure of 20 µM Xestospongin C, 10 µM Dantrolene or *UAS-IP_3_-sponge.m30* + *mcu[RNAi] + NDUFS7*[RNAi] with 10 µM Dantrolene and pan-neuronal Gal4 driven *NDUFS7*[RNAi] with 20 µM BAPTA-AM. (Q-T) Histograms showing average mEPSPs, EPSPs amplitude, and quantal content in the indicated genotypes. A minimum of 8 NMJs recordings of each genotype were used for quantification. \*\**p* < 0.05 (EPSP and QC: *NDUFS7*[RNAi] neuron + DMSO, wash vs *NDUFS7*[RNAi] neuron + BAPTA-AM, wash), *p < 0.05, \*\**p*=0.001, \*\*\**p* <0.0001; ns, not significant. Statistical analysis based on one-way ANOVA followed by post-hoc Tukey’s multiple-comparison test. Error bars represent mean ± s.e.m. Raw data for this figure are available in the S1 Data Excel file, tab Figure_5.

We observed no significant differences in EPSP amplitudes when we impaired *mcu* function neuronally (S14 Table). Similarly, we did not observe deficits in baseline synaptic activity by blocking RyR and IP_3_R, alone or in conjunction with *NDUFS7[RNAi]* (Fig. 5E-H, Q-T, S14 Table). However, when we concurrently impaired a combination of *NDUFS7*, *mcu*, and ER calcium store channels, we observed marked decreases in evoked amplitude (Fig. 5K-L, S, S14 Table). These results are consistent with a model in which mitochondrial calcium uptake, MCU activation, and ER (store) calcium efflux combine to stabilize synaptic strength.

If this idea were correct, then it should also be possible to chelate cytoplasmic calcium in a neuronal *NDUFS7[RNAi]* background and reveal neurotransmission defects. Direct application of the membrane-permeable chelator BAPTA-AM, followed by a wash to remove chelator residing in the saline, had no significant effect on baseline neurotransmission parameters (DMSO carrier + wash). But in the *NDUFS7[RNAi]* background, BAPTA-AM + wash significantly diminished evoked potentials, compared to mock-treated (DMSO + wash) NMJs. (Fig. 5 M-T, S14 Table).

To check if these effects on neurotransmission correlated with effects on active zone protein accumulation, we also conducted anti-Brp immunostaining experiments (Fig. S14). As before, neuronal knockdown of *NDUFS7* gene function triggered a marked, compensatory increase in presynaptic active zone material that was readily apparent by confocal microscopy (Figs. S14A, C, I, S15 Table). But this increase was reversed when combined with *mcu* gene function knockdown and pharmacological blockade of store calcium release channels (Figs. S14G-I, S15 Table). Together, our data indicate that loss of MCI subunits in neurons sensitizes synapses to decreases in intracellular calcium.

### A combination of mitochondria, glycolysis, and the TCA Cycle stabilizes NMJ function

Neuronal calcium handling and MCU play roles in NMJ stability that are uncovered by loss of MCI at the NMJ. Downstream of calcium handling, a logical hypothesis is that synaptic energy would play a role ^59^. If this were the case, would mitochondria be the sole energy source? Alternatively, in the absence of full mitochondrial function, could glycolysis theoretically substitute, as a homeostatic (or redundant) means for staving off synapse dysfunction?

We tested these ideas by limiting glycolysis as an energy source in two ways: 1) swapping out sucrose and trehalose in our recording saline in favor of 2-deoxy-D-glucose (a non-glycolytic sugar; (Fig. 6A); 2) addition of lonidamine (LDA) to the saline to acutely inhibit hexokinase (Fig. 6A). Control recordings with these conditions showed little effect on baseline physiology (Fig. 6A, D, F, H-J, S16 Table). However, when Mitochondrial Complex I was impaired neuronally through *NDUFS7[RNAi]* in combination with inhibition of glycolysis, there was a drop in evoked neurotransmission (Fig. 6C, E, G, H-J, S16 Table). This correlated with a failure to increase active zone material after *NDUFS7* gene knockdown (Fig. 6K-Q, S16 Table). These results match the idea that a combination of mitochondrial function or glycolysis can work to maintain normal levels of NMJ output.

**Figure 6:**
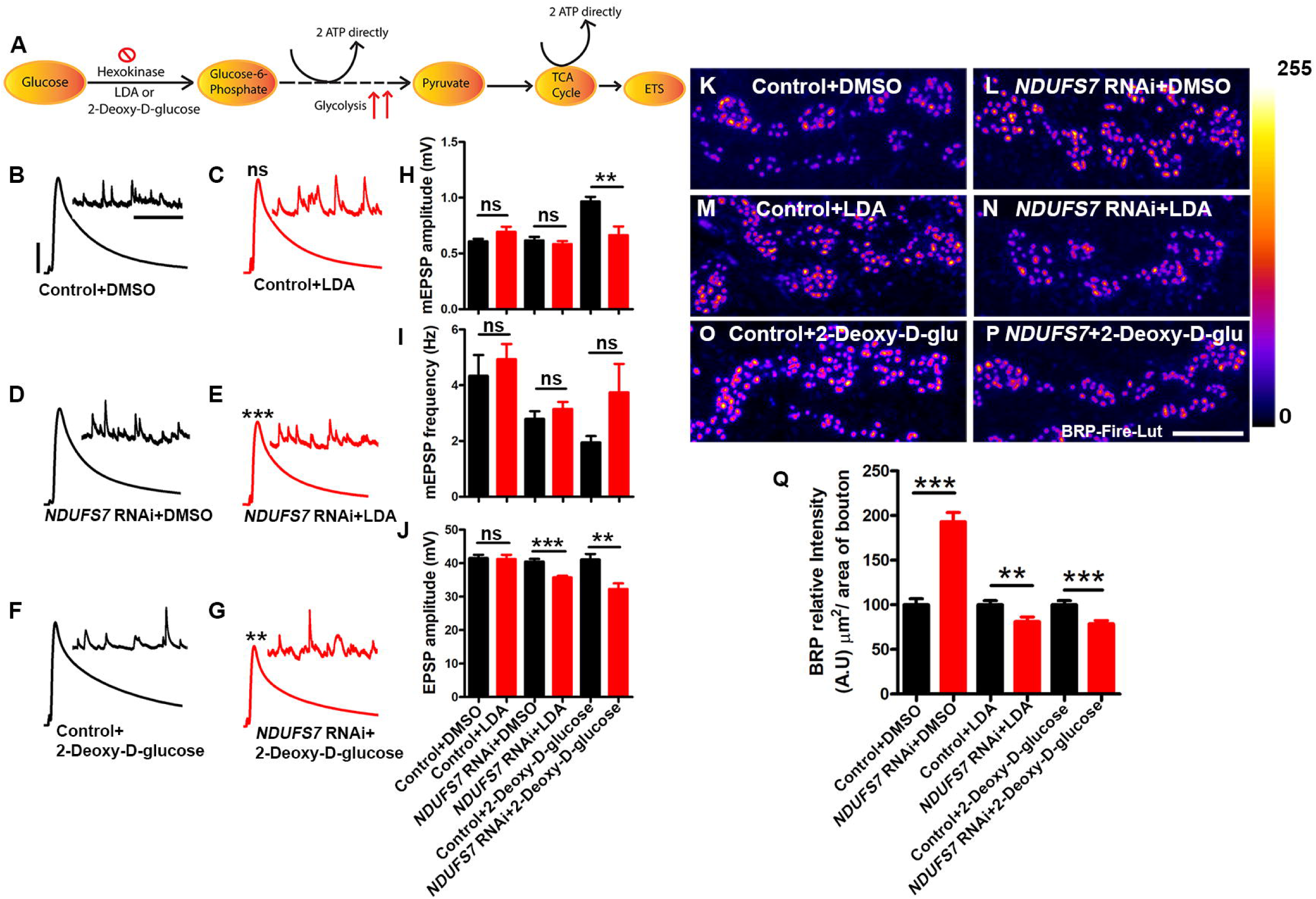
Loss of an MCI subunit necessitates glycolysis to regulate levels of active zone materials and stabilize synaptic strength. (A) Schematic illustrations showing steps of glucose metabolism and ATP production during glycolysis and TCA cycle in the cell. (B-G) Representative traces of mEPSPs and EPSPs in (B) pan-neuronal Gal4 control (*elaV(C155)-Gal4/+* with DMSO), (C) pan-neuronal Gal4 control (*elaV(C155)-Gal4/+*) with an acute application (30 minutes) of 150 µM Lonidamine (LDA) (D) pan-neuronal Gal4 driven *NDUFS7*[RNAi] (*elaV(C155)-Gal4)/+; NDUFS7[RNAi] /+* with DMSO), (E) pan-neuronal Gal4 driven *NDUFS7*[RNAi] (*elaV(C155)-Gal4)/+; NDUFS7[RNAi]/+*) with 150 µM Lonidamine, (F) pan-neuronal Gal4 control (*elaV(C155)-Gal4/+*) and HL3 containing 2-Deoxy-D-glucose and (G) pan-neuronal Gal4 driven *NDUFS7*[RNAi] (*elaV(C155)-Gal4)/+; NDUFS7[RNAi] /+*) with HL3 containing 2-Deoxy-D-glucose. Scale bars for EPSPs (mEPSP) are x=50 ms (1000 ms) and y= 10 mV (1 mV). The EPSPs amplitudes were reduced in pan-neuronal Gal4-driven *NDUFS7*[RNAi] with an acute exposure of 150 µM Lonidamine (LDA) or pan-neuronal Gal4-driven *NDUFS7*[RNAi] in HL3 containing 2-Deoxy-D-glucose for 30 minutes. However, we saw a reduction in mEPSP amplitudes when incubated in HL3 containing 2-Deoxy-D-glucose for 30 minutes in pan-neuronal Gal4-driven *NDUFS7[RNAi] -*depleted larvae compared to the control animals. (H-J) Histograms showing average mEPSPs, EPSPs amplitude, and frequencies in the indicated genotypes. A minimum of 8 NMJ recordings of each genotype were used for quantification.,\*\**p*=0.003 (mEPSP amplitude: *elaV(C155)-Gal4/*+2-Deoxy-D-glucose vs *elaV(C155)-Gal4)/+;NDUFS7[RNAi] /* +2-Deoxy-D-glucose), \*\**p*=0.002 (EPSP amplitude: *elaV(C155)-Gal4/*+2-Deoxy-D-glucose vs *elaV(C155)-Gal4)/+;NDUFS7[RNAi] /*+2-Deoxy-D-glucose),\*\*\**p*=0.0001 (EPSP amplitude: *elaV(C155)-Gal4/*+LDA vs *elaV(C155)-Gal4)/+;NDUFS7[RNAi] /* +LDA); ns, not significant. Statistical analysis is based on the Student’s t-test for pairwise sample comparison. Error bars represent mean ± s.e.m. (K-P) Representative images of the A2 hemi segment of muscle 6/7 NMJs in the above-indicated genotypes immunostained with antibodies against the active zone scaffold bruchpilot (BRP:fire-LuT) to label the active zones. The BRP levels downregulated at the NMJs in panneuronal Gal4 driven *NDUFS7*[RNAi] either incubated with 150 µM LDA or in HL3 containing 2-Deoxy-D-glucose for 30 minutes. (K-P) Scale bar: 5 µm. (Q) Histograms showing quantification of BRP intensity in µm^2^ area of bouton at muscle 6/7 in the genotypes mentioned above. At least 8 NMJs of each genotype were used for quantification.\*\*\**p*<0.0001 (BRP levels: *elaV(C155)-Gal4/*+DMSO vs *elaV(C155)-Gal4)/+;NDUFS7[RNAi]/*+ +DMSO),\*\**p*=0.008 (BRP levels: *elaV(C155)-Gal4/*+LDA vs *elaV(C155)-Gal4)/+;NDUFS7[RNAi] /*+LDA), \*\*\**p*=0.0005 (BRP levels: *elaV(C155)-Gal4/*+2-Deoxy-D-glucose vs *elaV(C155)-Gal4)/+;NDUFS7[RNAi] /*+2-Deoxy-D-glucose). Error bars denote mean ± s.e.m. Statistical analysis based on one-way ANOVA followed by post-hoc Tukey’s multiple-comparison test. Raw data for this figure are available in the S1 Data Excel file, tab Figure_6.

We continued this line of investigation genetically. We acquired RNA interference-based transgenes to target five genes involved in *Drosophila* glycolysis or subsequent ATP generation in the Citric Acid (TCA) Cycle: *hexokinase A (hex-A)*, *hexokinase C (hex-C)*, *Citrate (Si) Synthase I*, *Isocitrate dehydrogenase* (*Idh*), and *Succinyl-coenzyme A synthetase* α *subunit 1* (*Scsα1*). We knocked down these genes neuronally, either alone or in combination with *NDUFS7[RNAi]* (Figs. S15 and S16). Neuronal impairment of *hex-C* had no effect on baseline neurotransmission, but *hex-A* impairment reduced it (Figs. S15A-E, S17 Table). Impairment of the TCA Cycle enzymes on their own had little-to-no effect on baseline neurotransmission (Fig. S15A, F-H, S17 Table). However, concurrent impairment of *NDUFS7* and most of these genes significantly blunted neurotransmission, with the exception being *hex-C* (Figs. S15I-N, S17 Table). Collectively, the data suggest that MCI works in conjunction with – or redundantly to – alternative energy-generating pathways to support normal levels of neurotransmission (Fig. S15O-Q). In the case of the hexokinases, *Drosophila hex-A* seems more important for this process than *hex-C*.

We also quantified active zone material accumulation. The results mirrored the neurotransmission tests: as before, neuronal *NDUFS7[RNAi]* impairment elicited enhanced active zone material (Fig. S16A-B, M, S18 Table). On their own, impairment of glycolysis or TCA Cycle genes had variable effects active zone material (Fig. S16C-G, M, S18 Table). But concurrent neuronal impairment of *NDUFS7* and any of the glycolysis or TCA Cycle genes reversed active zone enhancement (Fig. S16H-M, S18 Table).

### MCI subunits in muscle are required for proper synapse development

We previously reported that impairments of MCI diminish *Drosophila* NMJ growth ^13^. The roles of MCI in specific tissues for this developmental process were unclear. For the present study, we tested for tissue-specific roles of MCI in NMJ development. To visualize NMJ boutons, we co-stained larval fillets with anti-Horseradish Peroxidase (HRP) a presynaptic membrane marker, and anti-Discs Large (Dlg), a postsynaptic density marker ^63,64^.

On a coarse level, MCI loss in muscle (*BG57-Gal4* > *UAS-NDUFS7[RNAi]*) caused a severe reduction in average bouton size, a decrease in bouton number, a notable decrease in Dlg expression, and a bouton “clustering” phenotype (Fig. 7 A-B), reminiscent of what we previously reported^13^. To quantify these observations, we measured bouton number, muscle area, and branch number per muscle in the third-instar larval NMJ synapses (S19 Table). We found that *NDUFS7* muscle knockdown resulted in a significant reduction in all these parameters compared to controls (Fig 7J-L, U-V, S19 Table).

**Figure 7:**
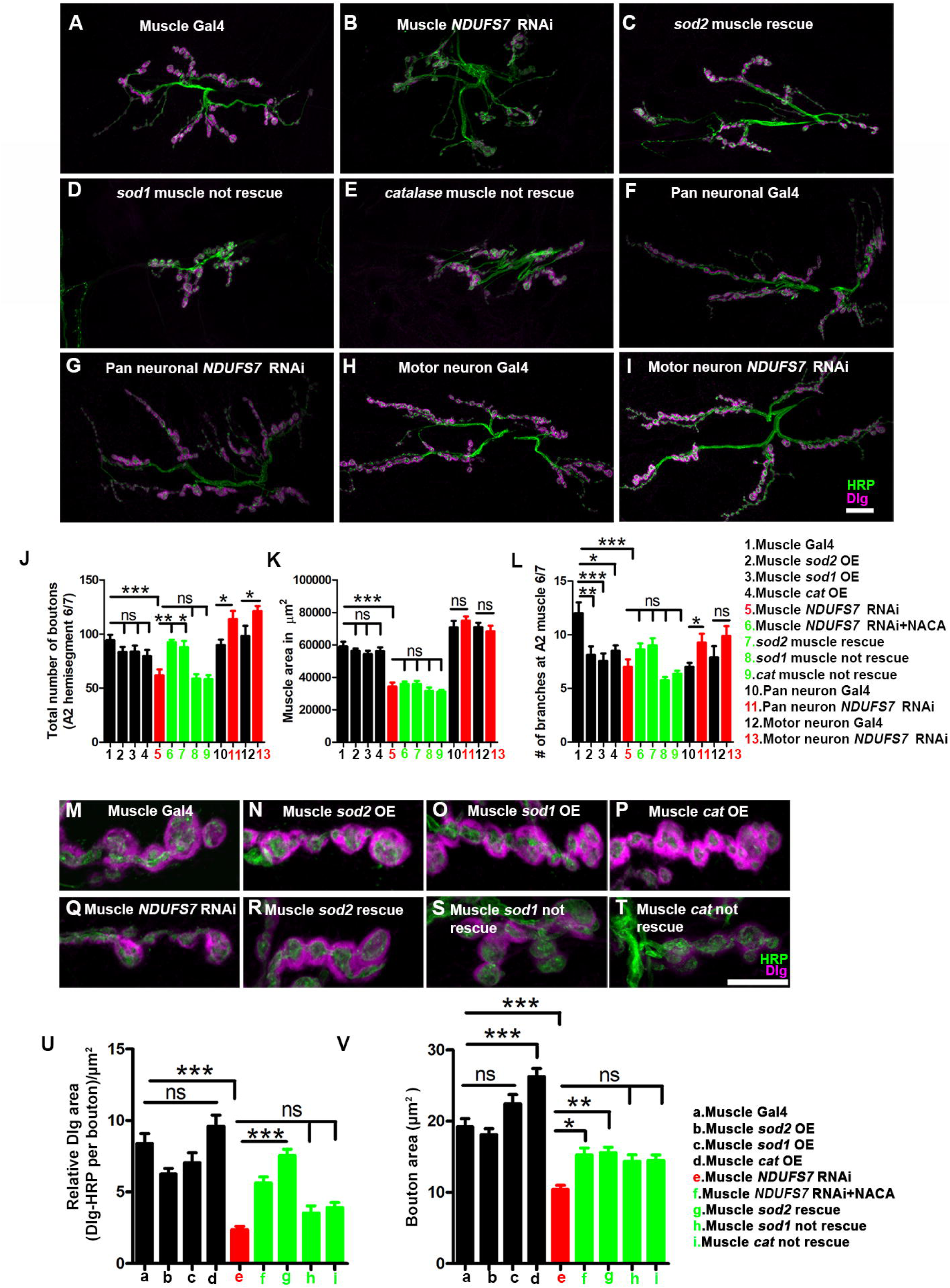
***NDUFS7* in muscles is required to promote normal synapse growth** (A-I) Representative confocal images of NMJ synapses at muscle 6/7 of (A) Muscle-Gal4 control (*BG57-Gal4/+*), (B) Muscle Gal4 driven *NDUFS7*[RNAi] (*NDUFS7[RNAi] /+; BG57-Gal4/+*), (C) *sod2* muscle rescue (*UAS-Sod2/NDUFS7[RNAi] ; BG57-Gal4/BG57-Gal4*), (D) *Sod1* muscle rescue (*UAS-Sod1/NDUFS7[RNAi] ; BG57-Gal4/BG57-Gal4*), (E) Catalase muscle rescue (*UAS-Cat/NDUFS7[RNAi]; BG57-Gal4/BG57-Gal4*), (F) Pan neuronal Gal4 control (*elaV(C155)-Gal4)/+*), (G) *elaV(C155)-Gal4)-Gal4* driven *NDUFS7*[RNAi] (*elaV(C155)-Gal4/+; NDUFS7[RNAi] /+*), (H) Motor neuron Gal4 control *(D42-Gal4/+)* and (I) *D42*-Gal4-driven *NDUFS7*[RNAi] (*NDUFS7[RNAi] /+; D42-Gal4/+*) double immunolabeled with Dlg (magenta) and HRP (green) antibodies. (A-I) Scale bar: 10 µm. The NMJ morphological defects in *NDUFS7*[RNAi] were restored upon co-expression of *UAS-Sod2 in* muscle; however, it did not rescue with *UAS-Sod1* or *UAS-Catalase* transgene. (J-L) Histograms show the number of boutons, muscle area, and average NMJ length at muscle 6/7 of A2 hemi segment in the indicated genotypes. \**p* <0.05, \**p*=0.006 (# of boutons: Muscle *NDUFS7*[RNAi] vs *sod2* muscle rescue), \*\**p*=0.0002 (# of boutons: Muscle *NDUFS7*[RNAi] vs Muscle *NDUFS7*[RNAi] +NACA), \**p*=0.009 (# of branches: Muscle Gal4 vs Muscle *sod2* OE), \*\*\**p*=0.002 (# of branches: Muscle Gal4 vs Muscle *sod1* OE), \**p*=0.008 (# of branches: Muscle Gal4 vs Muscle *cat* OE), \*\*\**p*=0.001 (Muscle Gal4 vs Muscle *NDUFS7*[RNAi]), \*\*\**p*<0.0001; ns, not significant. Statistical analysis based on one-way ANOVA with post-hoc Tukey’s test for multiple and Student’s t-tests for pairwise comparison. Error bars represent mean ± s.e.m. (M-T) Representative confocal images of boutons at the third instar larval NMJ synapse in (M) Muscle-Gal4 control (*BG57-Gal4/+*), (N) Muscle Gal4 driven *UAS-Sod2* (*UAS-Sod2*/*+; BG57-Gal4/+*), (O) *UAS-Sod1* (*UAS-Sod1*/*+; BG57-Gal4/+*), (P) *UAS-Catalase* (*UAS-Catalase*/*+; BG57-Gal4/+*) (Q) Muscle *NDUFS7*[RNAi] (*NDUFS7[RNAi] /+; BG57-Gal4* (R) *sod2* muscle rescue (*UAS-Sod2/NDUFS7[RNAi] ; BG57-Gal4/BG57-Gal4*), (S) *sod1* muscle rescue (*UAS-Sod1/NDUFS7[RNAi] ; BG57-Gal4/BG57-Gal4*) and (T) Catalase muscle rescue (*UAS-cat/NDUFS7[RNAi] ; BG57-Gal4/BG57-Gal4*) animals double immunolabeled with anti-HRP (green) and anti-Dlg (magenta) antibodies. (M-T) Scale bar: 5 µm. Note that the gross morphology of SSRs and immunoreactivity of Dlg were reduced in *NDUFS7[RNAi]* animals. As mentioned, phenotypes were restored to wild-type level when *NDUFS7*[RNAi] -depleted flies were reared in NACA or by genetically expressing *Sod2 transgene* in muscle. (U-V) Histograms showing normalized synaptic fluorescence of Dlg and bouton area in the indicated genotypes. \**p* <0.01, \*\**p* <0.001, \*\*\**p*<0.0001; ns, not significant. Error bars represent mean ± s.e.m. Statistical analysis based on one-way ANOVA with post-hoc Tukey’s test for multiple and Student’s t-tests for pairwise comparison. Raw data for this figure are available in the S1 Data Excel file, tab Figure_7.

In contrast to muscle knockdown, pan-neuronal or motor neuron-specific knockdown of *NDUFS7* showed slight NMJ overgrowth phenotypes (Fig. 7F-L, S19 Table). As was the case with the active zone enhancement associated with neuronal *NDUFS7* knockdown (Fig. 4), this NMJ overgrowth could represent a developmental mechanism to stave off dysfunction caused by missing neuronal mitochondria. Taken together, our NMJ immunostaining results indicated to us that blunted NMJ growth due to MCI loss was likely due to muscle MCI dysfunction.

We tested if the NMJ undergrowth phenotypes could be due to the increased levels of mitochondrial reactive oxygen species we had observed in muscle (mROS) (Fig. S2F). If that idea were correct, then the undergrowth phenotypes should be reversed if mitochondrial mROS were scavenged. Consistent with this idea, bouton number and synaptic undergrowth phenotypes were fully restored to wild-type levels when the *NDUFS7[RNAi]* muscle knockdown animals also had *UAS-Sod2* transgenically expressed in the muscles (Fig. 7C, J-L, R, S19 Table). They were also restored to wild-type levels when muscle *NDUFS7* knockdown animals were raised on food containing the antioxidant N-acetyl cysteine amide (NACA) (Fig. 7J-L, S19 Table). By contrast, none of these NMJ growth parameters were restored to wild-type levels if scavengers *UAS-Sod1* or *UAS-Catalase* were misexpressed in the muscle (Fig. 7D-E, J-L, S19 Table).

### Loss of MCI and mROS in muscle disorganize NMJ postsynaptic densities

The muscle Dlg-Spectrin network functions as an organizing scaffold for synaptic assembly ^64,65^. Dysregulation of this network can lead to an aberrant muscle subsynaptic reticulum (SSR) ^64^. Therefore, one possible target of excess mROS in the absence of MCI function is Dlg. Dlg is the fly homolog of PSD-95/SAP97/PSD-93, and it is a member of the membrane-associated guanylate kinase (MAGUK) family of NMJ scaffolding proteins ^64^. It is present both within presynaptic boutons and in the portion of the SSR closest to the bouton.

Using the same antibodies detailed above (anti-Dlg and anti-HRP), we used a high magnification to examine at the postsynaptic densities closely. By confocal immunofluorescence, Dlg area was significantly reduced when *NDUFS7* was depleted postsynaptically by[RNAi] (Fig. 7M-U, S19 Table). To quantify the relative Dlg area, we measured Dlg area with respect to HRP (Relative Dlg area = Dlg area minus HRP area) in type 1b boutons at muscle 6/7 of the A2 hemi segment (Fig. 7U). Compared with the control synapses, *NDUFS7* knockdown resulted in a significant reduction in the relative Dlg area (Fig 7M-U, S19 Table). Consistently, the relative α-Spectrin area was also reduced when *NDUFS7* was depleted in muscle (Fig. S17A-F, S7 Table).

Next, we scavenged mROS to check how that affected the postsynaptic densities. The relative Dlg and α-Spectrin areas were restored to wild-type levels when animals were grown in a media containing NACA (*NDUFS7[RNAi]/+; BG57-Gal4*/+ with NACA) or genetically expressing *UAS-Sod2* in the muscle (*UAS-Sod2/NDUFS7[RNAi]; BG57-Gal4/BG57-Gal4*) (Fig.7R, U, S19 Table; Fig. S16D-F, S7 Table). By contrast, Dlg levels were not restored while expressing other scavengers in the muscle, encoded by *UAS-Sod1* and *UAS-Catalase* transgenes (Fig. 7S-U, S19 Table). We conclude that depletion of MCI subunits in the muscle disables postsynaptic density formation via the formation of mitochondrial reactive oxygen species intermediates.

The postsynaptic density (PSD-95/Dlg) has been shown to cluster glutamate receptors at the SSR ^66^. Our results raised the possibility that glutamate receptor clusters could be disrupted when *NDUFS7* was depleted in the muscle (Fig. 8). To test this idea, we simultaneously immunostained NMJs with antibodies against Brp (neuron, presynaptic active zone) and glutamate receptor clusters (muscle). In controls, these pre- and postsynaptic structures were directly apposed to one another (Fig. 8A, G). But when *NDUFS7* gene function was depleted, we observed “missing” GluRIIA and GluRIII receptor clusters (Fig. 8 B, F, H, K, S7 Table), i.e., Brp puncta without apposed glutamate receptors. These lack of apposition phenotypes were fully reversed by raising the larvae with NACA or genetically expressing *UAS-Sod2* in the muscle (Fig. 8 D-F, I-K, S7 Table). However, as with prior tests, these phenotypes were not reversed by *UAS-Sod1* or *UAS-Catalase* overexpression (Fig. S18). Together, our data indicate that loss of MCI subunit in the muscle (*NDUFS7*) disrupts several aspects of the postsynaptic density organization, and these disruptions are likely due to the accumulation of mitochondrial ROS in the muscle.

**Figure 8:**
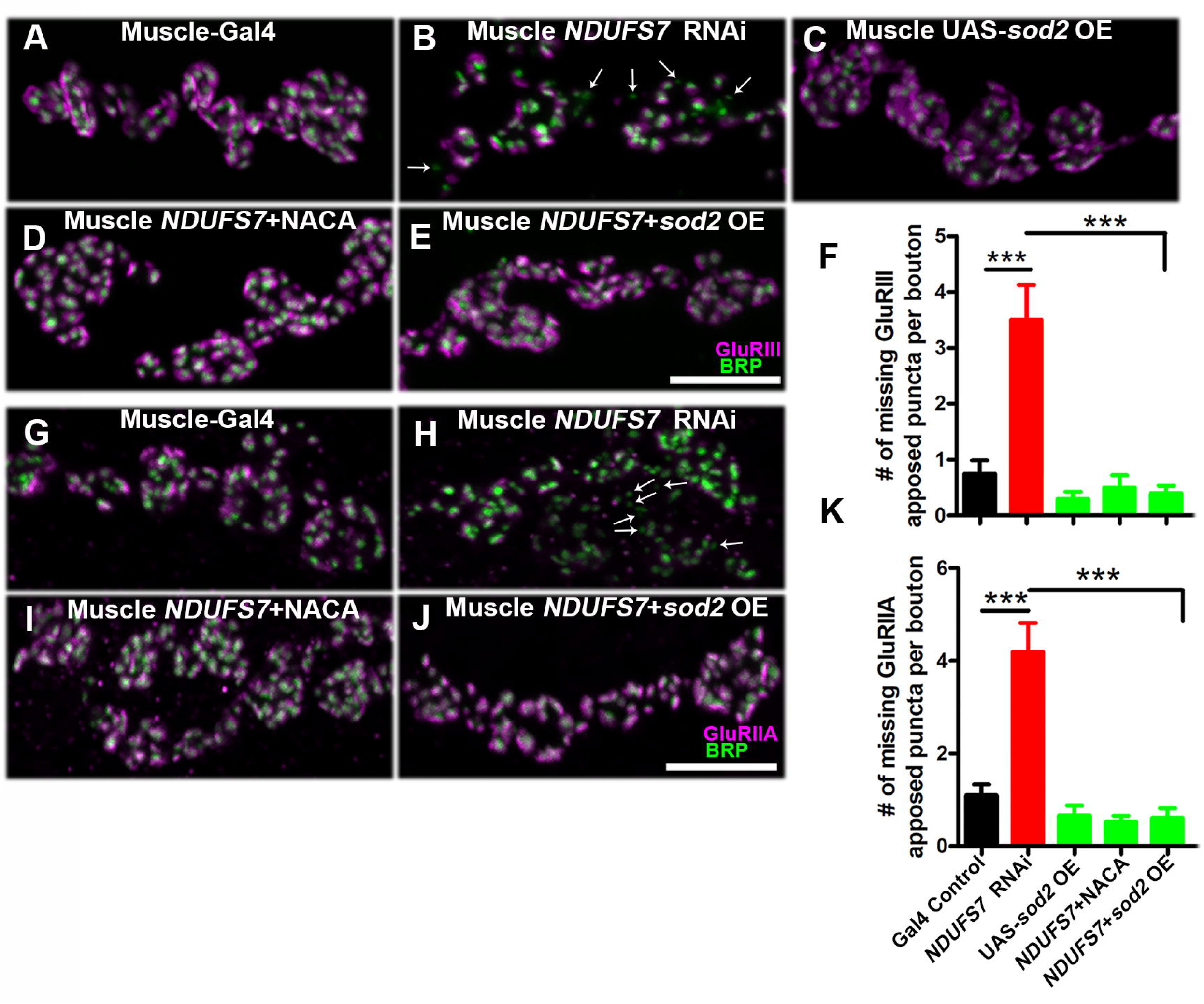
***NDUFS7* subunit in muscle affects the organization of GluRs cluster in *Drosophila*.** Representative confocal images of boutons at third instar larval NMJ synapse in (A) Muscle-Gal4 control (*BG57-Gal4/+*), (B) Muscle *NDUFS7*[RNAi] (*NDUFS7[RNAi] /+; BG57-Gal4/+*), (C) Muscle Gal4 driven *UAS-Sod2* (*UAS-Sod2*/*+; BG57-Gal4/+*), (D) *NDUFS7[RNAi] /+; BG57-Gal4/+*NACA) and (E) *sod2* muscle rescue (*UAS-Sod2/NDUFS7[RNAi] ; BG57-Gal4/BG57-Gal4*) animals immunolabeled with active zone marker BRP (green) and anti-GluRIII (magenta) antibodies. Scale bar: 5 µm. Note that GluRIII apposed clusters with BRP are missing in the *NDUFS7*[RNAi]-depleted animals (marked in arrow) compared to control. These phenotypes were restored to normal when *NDUFS7*[RNAi]-depleted flies were reared in media containing NACA or genetically expressing *Sod2 transgene* in muscle. (F) Histograms showing quantification of the number of missing BRP-GluRIII apposed puncta per bouton in the indicated genotypes. (G-J) Similar phenotypes were observed when analyzed for BRP-GluRIIA apposed clusters in boutons. (K) Histograms showing quantification of the number of missing BRP-GluRIIA apposed puncta per bouton in the indicated genotypes. \*\*\**p*<0.0001. Error bars represent mean ± s.e.m. Statistical analysis based on one-way ANOVA with post-hoc Tukey’s test for multiple comparisons. Raw data for this figure are available in the S1 Data Excel file, tab Figure_8.

### Loss of MCI subunits in muscle diminishes evoked NMJ neurotransmission through postsynaptic mROS

Mitochondria and reactive oxygen species influence presynaptic vesicle release and plasticity at synapses. This has been shown in diverse model systems like flies, mice, and worms ^52,53,67–69^. In our prior work, we identified an NMJ neurotransmission defect when MCI function is impaired ^13^, but based on our current study, that defect does not seem like it was dependent upon MCI’s neuronal functions (Figs. 1, 4, 5, 6). Therefore, we turned to analyzing postsynaptic muscle MCI and mROS to check if these parameters influenced neurotransmission.

We performed sharp electrode electrophysiological recordings of miniature and evoked excitatory postsynaptic potentials (mEPSP and EPSP) at NMJ muscle 6, hemi segment A2. We also used average mEPSP and EPSP values to estimate quantal content (QC) for each NMJ. For the most part, mEPSP values remained steady (Figs. 9A-E, S20 Table), with some exceptions. The starkest phenotypes came in terms of evoked amplitudes (Figs. 9A-D, F, S20 Table), sometimes due to changes in QC (Fig. 9G) or combinatorial changes in both mEPSP (Fig. 9E) and QC (Fig. 9G).

**Figure 9:**
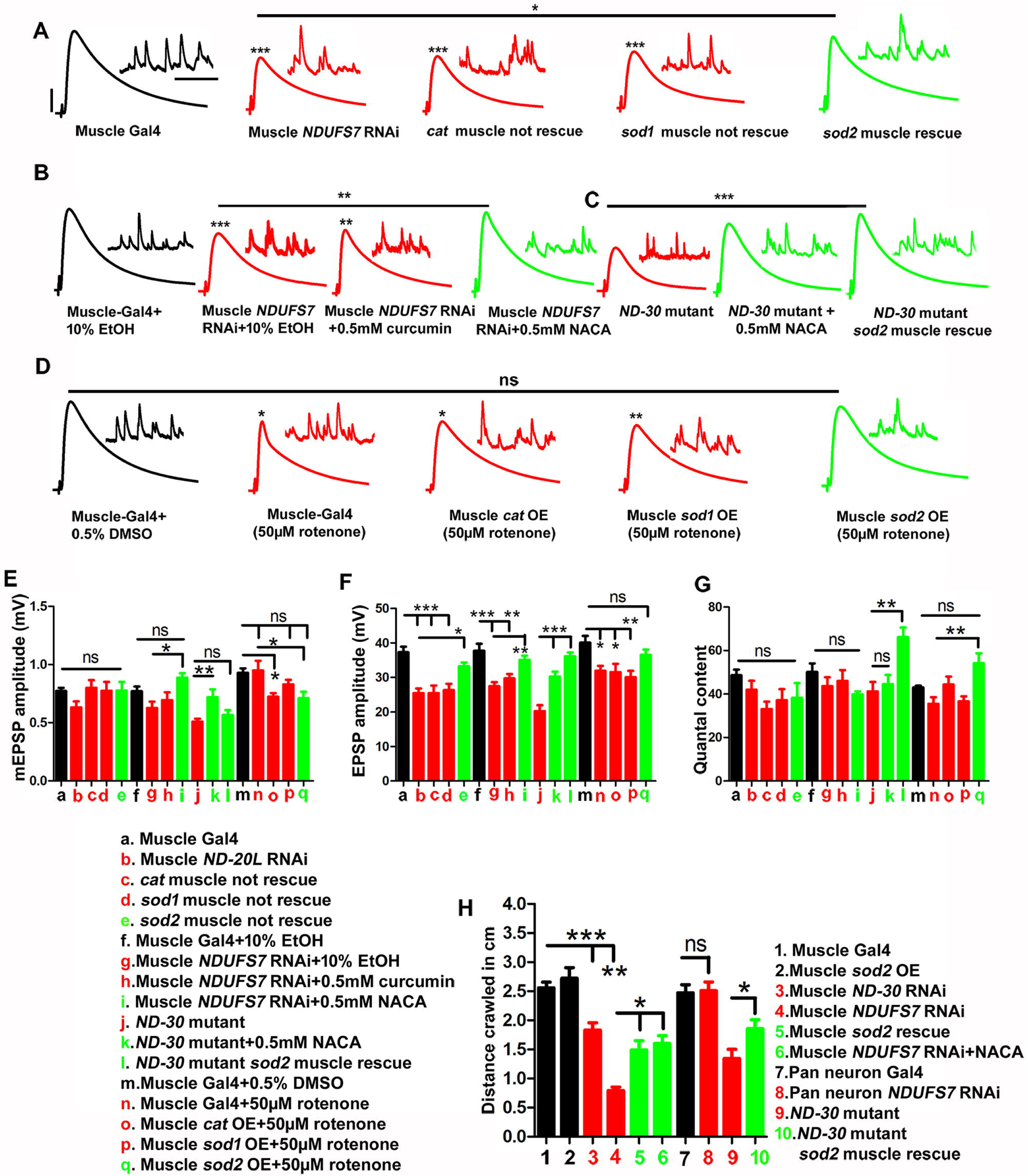
Loss of MCI subunits affects synaptic transmission via the formation of excess ROS in the muscle. (A) Representative traces of mEPSPs and EPSPs in muscle-Gal4 control (*BG57-Gal4/+*), muscle Gal4 driven *NDUFS7*[RNAi] (*NDUFS7[RNAi] /+; BG57-Gal4/+),* muscle *Catalase* rescue (2X muscle-Gal4>*UAS-Catalase/NDUFS7*[RNAi] : *UAS-Catalase/NDUFS7[RNAi] ; BG57-Gal4/BG57-Gal4*), muscle *Sod1* rescue (2X muscle-Gal4>*UAS-Sod1/NDUFS7*[RNAi] : *UAS-Sod1/NDUFS7[RNAi] ; BG57-Gal4/BG57-Gal4*), muscle *Sod2* rescue (2X muscle-Gal4>*UAS-Sod2/NDUFS7*[RNAi] : *UAS-Sod2/NDUFS7[RNAi] ; BG57-Gal4/BG57-Gal4*) animals. Note that EPSPs amplitudes were reduced in *NDUFS7[RNAi],* and the phenotype was restored to wild-type levels by expressing *Sod2 transgene* in the muscle but not with *Sod1* and *Catalase* transgenes. (B) Representative traces of mEPSPs and EPSPs in muscle Gal4 control (*BG57-Gal4/+*) larvae raised on 10% EtOH, *NDUFS7* muscle[RNAi] (*NDUFS7[RNAi] /+; BG57-Gal4/+*) raised on 10% EtOH, *NDUFS7* muscle[RNAi] (*NDUFS7[RNAi] /+; BG57-Gal4/+*) raised on 0.5mM curcumin, *NDUFS7* muscle[RNAi] (*NDUFS7[RNAi] /+; BG57-Gal4/+*) raised on 0.5mM NACA, (C) Representative traces of mEPSPs and EPSPs in *ND-30^epgy^/Df* mutants, *ND-30^epgy^/Df* mutants raised on 0.5 mM NACA and *UAS-Sod2* muscle rescued *ND-30^epgy^/Df* mutant (*UAS-Sod2/+; ND-30 (Df), BG57-Gal4/ND-30^epgy^*) animals. The EPSPs amplitudes were restored to wild type when muscle depleted *NDUFS7[RNAi]* larvae were raised in food containing NACA or by *UAS-Sod2* muscle overexpression in *NDUFS7[RNAi]* depleted animals. (D) Representative traces of mEPSPs and EPSPs in muscle-Gal4 control (*BG57-Gal4/+*) larvae raised on DMSO, muscle Gal4 (*BG57-Gal4/+*) larvae raised on 50µM rotenone (complex I inhibitor), *UAS-Catalase (UAS-Catalase/+; BG57-Gal4/)*, *UAS-Sod1(UAS-Sod1/+; BG57-Gal4/),* and *UAS-Sod2 (UAS-Sod2/+; BG57-Gal4/+)* muscle over-expression animals raised on 50µM rotenone. The EPSPs amplitudes were suppressed in rotenone raised larvae overexpressing *UAS-Sod2* in muscle *(UAS-Sod2/+; BG57-Gal4/)* due to its free radical scavenging activity. Scale bars for EPSPs (mEPSP) are x=50 ms (1000 ms) and y= 10 mV (1 mV). (E-G) Histograms showing average mEPSPs, EPSPs amplitude, and quantal content in the indicated genotypes. Minimum 8 NMJs recordings of each genotype were used for quantification. (H) Histogram representing crawling behavior (in cm) of the larvae in the indicated genotypes. Knocking down *NDUFS7*[RNAi] or *ND-30[RNAi]* in muscle and *ND-30^epgy^/Df* mutants showed a severe defect in crawling behavior. The abnormal crawling behavior was rescued by expressing a *Sod2 transgene* in the muscle or rearing the larvae in a media containing NACA. Moreover, neuronally depleting *NDUFS7* did not show any notable change in crawling defects. Minimum 10 animals were analyzed for crawling behavioral analysis. \**p* < 0.05 (mEPSP amplitude: Muscle *NDUFS7*[RNAi] +10% EtOH vs Muscle *NDUFS7*[RNAi] + 0.5 mM NACA),\*\**p*=0.006 (mEPSP amplitude: *ND-30* mutant vs *N-30* mutant + 0.5 mM NACA), \**p*=0.0004 (mEPSP amplitude: Muscle-Gal4+0.5% DMSO vs. Muscle *Cat* OE + 50 µM rotenone),\**p*=0.039 (mEPSP amplitude: Muscle Gal4+50mM rotenone vs Muscle *Sod2* OE+ 50 µM rotenone), \**p*=0.001 (EPSP amplitude: Muscle *NDUFS7[RNAi]* vs *Sod2* muscle rescue),\*\**p*=0.003 (EPSP amplitude: Muscle Gal4+10% EtOH vs Muscle *NDUFS7[RNAi]* +0.5mM curcumin), \*\**p*=0.0004 (EPSP amplitude: Muscle *NDUFS7*[RNAi] +10% EtOH vs Muscle *NDUFS7*[RNAi] +0.5mM NACA), \**p*=0.004 (EPSP amplitude: Muscle Gal4+0.5% DMSO vs Muscle Gal4+50 µM rotenone), *p=0.015 (EPSP amplitude: Muscle-Gal4+0.5% DMSO vs Muscle *cat* OE+50 µM rotenone),\*\**p*=0.015 (EPSP amplitude: Muscle-Gal4+0.5% DMSO vs Muscle *sod1* OE+50 µM rotenone), \*\**p*=0.001 (QC*: ND-30* mutant vs *sod2* muscle rescue),\*\**p*=0.003 (QC: Muscle Gal4+50 µM rotenone vs Muscle *Sod*2 OE+50 µM rotenone), \*\**p*=0.0002 (Distance crawled: Muscle Gal4 vs Muscle *ND-30*[RNAi]), \*\*\**p* <0.0001; ns, not significant. Statistical analysis based on one-way ANOVA with post-hoc Tukey’s test for multiple and Student’s t-test for pairwise comparison. Raw data for this figure are available in the S1 Data Excel file, tab Figure_9.

Evoked synaptic vesicle release (EPSP) was significantly reduced when *NDUFS7* was depleted in muscles by RNAi (Fig. 9A, F, S20 Table). That reduction was reversed after scavenging mitochondrial ROS in the muscle. Indeed, transgenic muscle-driven *Sod2* gene expression suppressed the *NDUFS7[RNAi]* phenotype, but expression of the *Catalase* and *Sod1* did not (Fig. 9A, F, S20 Table). We also tested if feeding a ROS scavenger to developing larvae would reverse the same neurotransmission defect. Carrier feeding alone (10% EtOH) did not affect evoked neurotransmission, nor did it influence the neurotransmission loss caused by *NDUFS7[RNAi]* (Fig. 9B, F, S20 Table). Feeding larvae 0.5 mM NACA successfully reversed the phenotype, but the nonspecific additive curcumin had no effect (Fig. 9B, F, S20 Table).

We checked different MCI manipulations. First, we examined hemizygous *ND-30^EY036^*^64^*^/Df^*genetic mutants. As with muscle-driven *NDUFS7[RNAi]*, the *ND-30* mutant NMJs had blunted evoked neurotransmission, but this defect was successfully reversed by ROS scavengers (Fig. 9C, S20 Table). We also impaired MCI pharmacologically, by feeding larvae 50 µM rotenone (or 0.5% DMSO carrier control), similar to conditions we previously published ^13^. As with the prior study, rotenone blunted neurotransmission (Fig. 9D, F, G, S20 Table), but this effect was ameliorated by a genetic background overexpressing *UAS-Sod2* in the muscle (Fig. 9D, F, G, S20 Table). By contrast, overexpressions of *UAS-Catalase* and *UAS-Sod1* were not effective at reversing the effect of rotenone (Fig. 9D, F, G, S20 Table).

Finally, we performed behavioral experiments on *NDUFS7* and *ND-30[RNAi]* and mutant animals. Consistent with the electrophysiological recordings, MCI muscle-depleted and mutant animals showed severe defects in crawling ability. The crawling behavior was rescued by expressing the *UAS-Sod2 transgene* in the muscle or raising the larvae in a media containing NACA (Fig. 9H, S21 Table). By contrast, neuronal depletion of *NDUFS7* in larvae did not show any significant crawling defects (Fig. 9H, S21 Table). Together, our data suggest that excess mitochondrial ROS accumulation in muscle (mROS) diminishes baseline synaptic physiology when MCI activity is lost, and it also triggers aberrant crawling behavior.

## DISCUSSION

We uncovered novel aspects of NMJ synapse biology controlled by Mitochondrial Complex I (MCI). Impairment of MCI causes profound cytological phenotypes in synaptic tissues (Figs. 1-3) ^13^. By examining mitochondria directly, we discovered shared phenotypes between MCI loss and loss of the *Drosophila* Mitofusin, Marf (Fig. 2). Additionally, with MCI loss, we noted an enhancement of mitochondrial reactive oxygen species (ROS, Fig. S1), consistent with prior work ^36–40^.

Unexpectedly, these perturbations spur functionally opposite responses in presynaptic neurons vs. postsynaptic muscles. In motor neurons, MCI loss and mitochondrial ROS appear to trigger a compensatory response, where the underlying cytological problems are offset by an increase in active zone material, resulting in normal levels of evoked excitation (Fig. 4). This process requires known intracellular calcium signaling components (Fig. 5). It also appears to require energy stores because loss of glycolysis – which may function as a supplemental energy source to mitochondria – abrogates the presynaptic compensation (Fig. 6). By contrast, in the muscle, MCI loss and mitochondrial ROS trigger a destructive response, where there is a disassembly of the postsynaptic density (Fig. 7). This disassembly correlates with mis-apposition of pre- and postsynaptic structures (Fig. 8) and defective neurotransmission (Fig. 9).

### Disruption of mitochondrial dynamics, ROS, and physiological responses in *Drosophila*

Energy is needed for normal levels of synaptic transmission ^2^. Intuitively, a loss of synaptic mitochondria should blunt transmission because transmission requires energy. Consistently, several labs have previously implicated mitochondrial dynamics in *Drosophila* synapse function, including mitochondrial fission (Dynamin related protein 1, Drp1 ^51^), fusion (Mitofusin/dMarf ^31^), trafficking (Miro and Milton ^50,70,71^), or quality control (Pink and Parkin ^72–75^). Additionally, it has been established from model organisms such as flies, worms, and mice that any misregulation in mitochondrial distribution could affect synaptic activity ^51,59,76,77^. Adding to that work, we uncovered synaptic transmission and developmental phenotypes after depletion of Mitochondrial Complex I (MCI) at the NMJ ^13^ potentially related to defective mitochondrial fusion (this study). Additionally, ROS has been studied in the context of mitochondrial dysfunction ^78^. Excess ROS can trigger mitochondrial calcium uptake and subsequently trigger apoptosis or degeneration of neurons or neural support cells ^79,80^. MCI deficiency elevates ROS levels, and this process can promote the fragmentation of mitochondria in cells like fibroblasts ^81^.

Numerous papers using *Drosophila* as a model have examined potential connections between MCI, ROS, aging, and physiology. The consequences of MCI loss and the induction of ROS are context dependent. They are not always in obvious agreement, either from tissue to tissue or from study to study. For example, one analysis demonstrated that reverse electron transfer (RET) at MCI increases ROS production, and this occurs with the normal aging process ^82^. Consistently, inhibition of RET decreases ROS and increases lifespan; this result would suggest that RET-induced ROS hurts survival ^82^. However, a different study reported the opposite finding under stress-inducing conditions. For that scenario, inhibition of RET decreases ROS production and decreases lifespan – while activation of RET enhances ROS and extends lifespan. Those findings would suggest that ROS is protective under some conditions ^83^. Effects are not limited to Complex I manipulation. *Drosophila* deficient for the Complex V subunit *bellweather* gene function demonstrate enhanced mitochondrial ROS; downstream this ROS, they have broadly enhanced levels of active zone material (BRP) in adult brains ^84^. Consistently, genetically boosting mitochondrial function in adults has the opposite effect – attenuated BRP accumulation in the fly brain ^84^. These findings closely match our larval motor neuron observations.

On the molecular level there are signaling puzzles uncovered by *Drosophila* work. For example, the unfolded protein response (UPR) is a key mediator when MCI function in lost. Neuronal impairment of the NDUFS1 (MCI subunit) homolog ND-75 elicits a spectrum of movement and seizure phenotypes, and it also activates the UPR on the cellular level ^85^. All those *Drosophila* neuronal phenotypes can be reversed when yeast ND1 is exogenously added to the system, which restores NADH dehydrogenase function (while not restoring ATP production) ^85^. Those findings suggest a damaging role in adult neurons for the MCI-deficient ROS-UPR axis. By contrast, another paper examined ND-75 disruption in adult muscle – and for this tissue, researchers found that elevated ROS levels trigger the UPR activation ultimately to preserve function ^86^. That suggests a protective in in muscles for the ROS-UPR axis ^86^.

Our study expands upon these results and supports the idea that MCI has roles at synaptic sites through disrupted mitochondrial dynamics. Lack of MCI in larval motor neurons causes loss of mitochondria at synaptic terminals (Fig. 1). This defect is linked to defective mitochondrial fusion (Fig. 2), which is known to maintain mitochondrial integrity ^87,88^. By contrast, lack of MCI in larval muscle appears to disassemble the postsynaptic density, which hurts neurotransmission (Figs. 7-9).

### A novel form of presynaptic homeostatic plasticity triggered by ROS?

ROS has been linked to short-term synaptic plasticity in *Drosophila* ^53^, as well as long-term potentiation (LTP) in mammals ^89–91^. Here, we uncovered a role for ROS in augmentation of active zone material when MCI is impaired in neurons. This finding could be considered a form of homeostatic plasticity: an increase in active zone components likely drives potentiated release to compensate for defective baseline synaptic transmission, which would be expected after the loss of an energy source like mitochondria.

Homeostatic augmentations of active zone material have been reported at the *Drosophila* NMJ. For example, *rab3* mutants have an increase in active zone material to offset decreased synapse growth ^92^. Additionally, ROS has been shown to be an obligate signal in *Drosophila* to maintain fundamental properties in both pre- and postsynaptic compartments, including at the NMJ ^53^. In our study, we observe an increase in active zone intensity (Fig. 4), which could overlap with mechanisms uncovered in those prior studies. Alternatively, it could be consistent with a form of homeostatic plasticity at the *Drosophila* NMJ called Presynaptic Homeostatic Potentiation (PHP). PHP is initiated when the activity of postsynaptic glutamate receptors is impaired. This decreases quantal size. The synapse detects the impairment, and muscle-to-nerve signaling drives an increase in presynaptic glutamate release ^93,94^. This happens in part through an increase in influx of calcium into the neuron through Ca_V_2 voltage-gated calcium channels ^93,95–98^. PHP coincides with an increase in the size of the readily releasable pool (RRP) of synaptic vesicles ^99–102^ and apparent increases in active zone protein content or organization ^54,56,103^. The general model is that these modifications drive the neuron to release more glutamate, offsetting the initial synaptic challenge.

Could the excess mitochondrial ROS that is caused by the loss of Complex I be triggering the same or overlapping homeostatic mechanisms? It is possible. Quenching presynaptic ROS by expression of *Sod2* or by feeding flies NACA led to a reversion of active zone material back down to the control levels (Fig. 4E). Under ideal conditions, control levels of active zone material would support normal neurotransmission (Fig. 4A, J). However, when combined with presynaptic *NDUFS7* loss (already depleting synaptic mitochondria; Figs. 1O, 2G), there is a loss in neurotransmission capacity (Fig. 4M).

The molecular underpinnings of this mitochondrial-loss-induced homeostatic plasticity are unknown, but prior work offers ideas. One possibility is that *Sod2* changes the redox status of the entire cell, which could potentially affect active zone components ^89^. Another possibility is that ROS could alter the release of calcium from intracellular stores such as ER; and in turn, this could induce calcium signaling to mitochondria, which would contribute to the formation of ATP and subsequent vesicle fusion ^59,60^. Consistent with this idea, we observed a reduced synaptic strength at the terminals after simultaneous blockade of calcium release and import from ER stores to mitochondria through genetic or pharmacological manipulations (Fig. 5). The link would not have to be a direct one. Indeed, a recent study indicated that activity-driven mitochondrial calcium uptake does not depend on the ER as a source of calcium to maintain normal synaptic strength ^59^. Future studies can refine conclusions about presynaptic calcium dynamics downstream of MCI disruption. This is possible through reagents in the *Drosophila* neurogenetic toolkit like compart-targeted genetic-encoded calcium indicators ^104^.

### Loss of MCI in muscle and subsequent ROS accumulation diminish synaptic excitation

Little is known about what role postsynaptic ROS plays in regulating synaptic plasticity at the NMJ. It is unlikely that ROS is an abstract signal that triggers wholesale destruction. It likely has specific targets. There are clues from previous work. For example, ROS signaling plays direct roles in the activity-dependent structural plasticity of motor neurons and postsynaptic dendrites ^53^. Additionally, postsynaptic ROS plays critical roles in dendritic development. In *Drosophila*, ROS appears to act as a developmental plasticity signal to regulate the size of the dendritic arbors. ^105^.

It is slightly surprising that the NMJ fails to compensate for a lack of MCI function in the muscle. This is because the NMJ employs multiple retrograde signals to stabilize function ^106^. One hypothesis is that MCI deficiency and ROS interfere with important muscle-to-nerve signals normally required for maintaining NMJ setpoint function. Some of these signals are required to correct acute challenges (minutes) to NMJ function. Some of the best characterized are: Bone Morphogenetic Protein (BMP) signaling ^107–109^; the Insomniac (adaptor)-Cul3 (ubiquitin ligase)-Peflin (substrate) signaling complex ^110^; and the endosomal recycling molecules class II PI3 kinase (PI3K) and Rab11 ^111^. Additionally, the field has uncovered instructive signaling molecules from the muscle that maintain robust neurotransmission, including: muscle-secreted Semaphorin-2b ^112^; and Target of Rapamycin (TOR) ^113,114^. Future studies can address whether these or similar targets are substrates of negative ROS regulation in the muscle.

The roles of ROS in muscle are less understood in terms of synapse regulation and function. In this study, we showed that excess ROS is sufficient to affect the organization of postsynaptic densities (Fig. 7), ionotropic glutamate receptor clusters (Fig. 8), and spectrin cytoskeleton (Fig. S11) in the muscle. The correlated uncoupling of pre- and postsynaptic structures seems to be responsible for neurotransmission defects (Fig. 9), and it likely coincides with broader structural instability ^65,115^. These phenotypes are reminiscent of other *Drosophila* mutant phenotypes showing a degenerative NMJ (e.g., ^65,115–117^). Consistent with a model of degeneration, we found behavioral defects in locomotion (Fig. 9).

Despite the profound defects that result from muscle depletion of MCI, we were able to reverse many of those structural and functional problems by culturing animals in an antioxidant-rich media (0.5mM NACA) or through muscle expression of *UAS-Sod2*. This is interesting finding for future work because it points to mitochondrial ROS as a signal of interest in the manifestation of synapse dysfunction. In subsequent work, we will take advantage of the *Drosophila* neurogenetic toolkit to identify specific targets.

### Limitations and Future Directions: Linking Mitochondrial ROS and Synapse Function

There are fundamental questions that remain to be answered in future work. Many of these deal with ROS. First, we have not identified the specific ROS molecule(s) responsible for the synaptic phenotypes we have observed. Our data point to mitochondrial matrix ROS because *UAS-Sod2* overexpression ameliorates MCI loss phenotypes, while *UAS-Catalase* and *UAS-Sod1* do not. Additionally, *UAS-Sod2[RNAi]* phenocopies the ROS accumulation associated with MCI loss.

If the relevant ROS molecule is signaling from the mitochondrial matrix, this could mean superoxide (O_2_•-) or hydrogen peroxide (H_2_O_2_) – or less likely, another species, like hydroxyl radicals (•OH). The function of superoxide dismutase enzymes is to convert superoxide into hydrogen peroxide ^118^. Because SOD2 is mitochondrial and it has a mollifying effect on MCI dysfunction in our system, we would hypothesize that mitochondrial matrix superoxide (O_2_•-) is the likely culprit for the synaptic phenotypes we observe. This idea would explain why other antioxidants like SOD1, and Catalase do not help. SOD1 accumulates mainly in the cytoplasm ^119,120^, while Catalase accumulates mainly in peroxisomes ^121,122^. NACA, which is the amide form of the antioxidant N-acetylcysteine, is membrane permeable; as such, it could accumulate in mitochondria ^123^. We have not affirmatively demonstrated sub-compartment mitochondrial localization of NACA in our system. However, recent work by other groups is consistent with a reductive function of NACA for mitochondria ^124,125^.

Even though our data point to mitochondrial ROS, the mechanistic connection between matrix O_2_•- and the synapse is not clear. Additionally, we do not know the relevant subcellular site(s) where ROS would need to accumulate to have its effects on the synapse. An intuitive idea is that the affected mitochondria and ROS would be near the synapse itself. For the NMJ, this is possible both in neurons and muscle. Yet our data do not neatly match that model. For example, six hours of rotenone exposure to the NMJ can enhance active zone accumulation, but only if the motor nerve is intact (Fig. S12). This finding suggests that the signaling downstream of mitochondrial ROS in neurons needs both time and space to work.

Concerning the muscle, mitochondrial ROS seems to have a damaging effect directly on the postsynaptic density. This effect hurts synaptic function. If the relevant site of mitochondrial ROS in muscle happens to be closer to the NMJ than the relevant site in neurons, then the damaging effects from the muscle could partly a function of proximity. Determining if it matters whether mitochondrial ROS is physically near the NMJ could inform the possible signaling modalities that govern mitochondrial regulation of synapse function.

## RESOURCE AVAILABILITY

### Lead Contact

The lead contact for this study is Dr. C. Andrew Frank.

### Materials Availability

Dr. C. Andrew Frank’s laboratory uses the fruit fly *Drosophila melanogaster* as a model organism. Consistent with ethical conduct of research, the lab maintains *Drosophila* stocks that it generates and publishes, as well as useful precursors and derivatives of those stocks. These stocks are available to researchers who make requests to Dr. Frank – or to the appropriate original source – and they are shipped promptly.

### Data and Code Availability

Consistent with the Data Sharing Policy for NIH-funded research, Dr. C. Andrew Frank’s laboratory will share data related to this study. The lab will share final data through publication of summary figures or tables. In addition, a large amount of final data derives from performing electrophysiological recordings or synapse imaging experiments and then performing subsequent analyses. Those experiments yield raw data in electronic form. Therefore, in addition to publishing final data, the lab will share raw data and related materials with researchers who make requests to Dr. Frank. No original coding was required to execute this study.

## Supporting information

Raw Data for Main Figures

Raw Data for Supplemental Figures

## Acknowledgments

We acknowledge the Developmental Studies Hybridoma Bank (University of Iowa, USA) for antibodies used in this study and the Bloomington Drosophila Stock Center (Indiana University, Bloomington, USA) for fly stocks. We thank members of the Frank lab for their helpful comments and discussions in this study. BM was supported in part by NIH/NINDS (https://www.ninds.nih.gov) grants to CAF (NS085164, NS130108, and NS136753). Collectively, the work was supported by those same NIH/NINDS grants, as well as funds from the University of Iowa’s Carver College of Medicine (https://medicine.uiowa.edu) to CAF. Funding supporting the work executed by SAB and XW was from an NIH/NINDS (https://www.ninds.nih.gov) R01 grant NS128040 to XW. The funders had no role in study design, data collection and analysis, decision to publish, or preparation of the manuscript.

## Author Contributions

BM, SAB, XW, and CAF designed the research; BM and SAB performed the research; BM, SAB, XW, and CAF analyzed the data; BM and CAF wrote the initial draft of the paper; BM, SAB, XW, and CAF wrote and edited the revision draft of the paper. For CRediT Taxonomy, the following contributions apply:

- BM - Conceptualization, Data Curation, Formal Analysis, Investigation, Methodology, Validation, Visualization, Writing – Original Draft Preparation, Writing – Review & Editing
- SAB - Conceptualization, Formal Analysis, Investigation, Methodology, Validation, Writing – Review & Editing
- XW - Conceptualization, Formal Analysis, Funding Acquisition, Methodology, Resources, Supervision, Validation, Writing – Review & Editing
- CAF - Conceptualization, Formal Analysis, Funding Acquisition, Methodology, Project Administration, Resources, Supervision, Validation, Writing – Original Draft Preparation, Writing – Review & Editing

## TABLES

All tables contain raw data supporting the graphs in the figures and are housed in the supplementary section.

## STAR METHODS

### Experimental Model Details

#### Drosophila husbandry

*Drosophila melanogaster* was cultured on a standard cornmeal media containing molasses and yeast prepared according to the Bloomington *Drosophila* Stock Center (BDSC, Bloomington, IN) recipe. Fruit fly husbandry was performed according to standard practices ^126^. Both male and female larvae were used for this study. Larvae were raised at 25°C in humidity-controlled and light-controlled Percival DR-36VL incubators (Geneva Scientific). For drug feeding experiments in larvae, the control, RNAi, and mutant animals were grown throughout life (egg laying to analysis) in media containing the relevant drug. Those drugs were 0.5 mM curcumin (Sigma Aldrich), 0.5 mM N-acetyl cysteine amide (NACA) (Sigma Aldrich), or 50 µM rotenone (Sigma Aldrich).

#### Drosophila stocks

*w*^1118^ was used as a non-transgenic wild-type stock ^127^. The GAL4 drivers used in this study were *elaV^C^*^155^–*Gal4* ^128^, *BG57-Gal4* ^64^, and *D42-Gal4* ^129^. For the neuronal experiments, a switch from *D42-Gal4* (motor neuron) to *elaV^C^*^155^–*Gal4* (pan-neuronal) was made between Figures 4 and 5 for practicality of genetic stock building with several markers in the latter experiments. The core neuronal results remained unchanged.

Several *UAS-RNAi* and genetic mutant lines were obtained from the Bloomington *Drosophila* stock center or directly from researchers who produced them (S18 Table includes the Bloomington Line (BL) numbers). We verified the effectiveness of *UAS-ND-20L[RNAi]* (*ND-20L^HMJ^*^23777^, now termed *UAS-NDUFS7[RNAi]*) in this study. Other mutant and transgenic lines have been characterized in prior studies. These include: *UAS-IP_3_-sponge* (*UAS-IP_3_-Sponge.m30* shared by Dr. Masayuki Koganezawa’s lab) ^126,130^; *UAS-Catalase* ^131,132^; *UAS-Sod1* ^133^; *UAS-Sod2* ^47^; *UAS-Sod2[RNAi]* ^134^; *Df(3L)ED4288* ^135^; *ND-30^epgy^* ^136^; *UAS-ND-30[RNAi]* ^12^; *UAS-mitoGFP* ^137^; *UAS-marf[RNAi]* ^31^; *UAS-drp1[RNAi]* ^138^; *UAS-CalX[RNAi]* ^139^; *UAS-mcu[RNAi]* ^140,141^; *UAS-hexokinase A[RNAi]* ^142^; *UAS-hexokinase C[RNAi]* ^142^; *UAS-idh[RNAi]* ^143^; *UAS-Cit.(Si)-Syn[RNAi]* ^144^; *UAS-Scsα[RNAi]* ^145^.

### Method Details Immunohistochemistry

Wondering third instar larvae were dissected and fixed on a sylgard Petri plate in ice-cold HL3 and fixed in 4% paraformaldehyde in PBS for 30 minutes or in Bouin’s fixative for 2 minutes as described earlier ^146^. Briefly, larvae were washed with PBS containing 0.2% Triton X-100 (PBST) for 30 min, blocked for an hour with 5% normal goat serum in PBST, and incubated overnight in primary antibodies at 4°C followed by washes and incubation in secondary antibodies. Monoclonal antibodies: anti-Dlg (4F3), anti-DGluRIIA (8B4D2), anti-CSP (ab49), anti-Synapsin (3C11), anti-Futsch (22C10), anti-Bruchpilot (nC82) and anti-α-Spectrin (3A9) were obtained from the Developmental Studies Hybridoma Bank (University of Iowa, USA) and were used at 1:30 dilution. Rabbit anti-GFP (Abcam) was used at 1:200 dilutions. Anti-GluRIII (1:100) (^147^; S18 Table) was gifted by Aaron DiAntonio (Washington University, St. Louis, U.S.A.). Fluorophore coupled secondary antibodies Alexa Fluor 488, Alexa Fluor 568 or Alexa Fluor 647 (Molecular Probes, ThermoFisher Scientific) were used at 1:400 dilution. Alexa 488 or 647 and Rhodamine conjugated anti-HRP were used at 1:800 and 1:600 dilutions, respectively. The larval preparations were mounted in VECTASHIELD (Vector Laboratories, USA) and imaged with a laser scanning confocal microscope (LSM 710; Carl Zeiss). All the images were processed with Adobe Photoshop 7.0 (Adobe Systems, San Jose, CA).

### Confocal imaging, quantification, and morphometric analysis

Samples were imaged using a Carl Zeiss scanning confocal microscope equipped with 63×/1.4 NA oil immersion objective using separate channels with four laser lines (405, 488, 561, and 637 nm) at room temperature. The stained NMJ boutons were counted using anti-Synapsin or anti-HRP costained with anti-Dlg on muscle 6/7 of A2 hemi segment, considering each Synapsin or HRP punctum to be a bouton. At least 8 NMJs were used for bouton number quantification. For fluorescence quantifications of GluRs, Dlg, Brp, α-Spectrin, HRP, and CSP, all genotypes were immunostained in the same tube with identical reagents, then mounted and imaged in the same session. Z-stacks were obtained using identical settings for all genotypes with z-axis spacing between 0.2-0.5 μm and optimized for detection without saturation of the signal.

### ROS and rotenone incubation assay

Larvae were dissected in ice-cold calcium-free HL3 to label ROS in neurons. ROS levels were detected in mitochondria by incubating live preparation in 1X Schneider’s media with MitoSOX Red (Molecular Probes, ThermoFisher Scientific) fluorogenic dye at 1:200 dilutions for 20-30 minutes. Briefly, larvae were washed with HL3, mounted in VECTASHIELD (Vector Laboratories, USA.), and immediately imaged in a laser scanning confocal microscope (LSM 710; Carl Zeiss). To study the effect of DMSO and rotenone (Sigma Aldrich) in BRP, larvae were dissected in HL3 and incubated in 1X Schneider’s media containing either 500 µM of rotenone or DMSO. After every 30 minutes, the old media was replaced with fresh media containing rotenone or DMSO. The above preparations were fixed with 4% paraformaldehyde, stained with anti-nc82 antibodies, mounted and imaged in a confocal microscope.

### Electrophysiology and pharmacology

All dissections and recordings were performed in modified HL3 saline ^148^ containing 70 mM NaCl, 5 mM KCl, 10 mM MgCl2, 10 mM NaHCO3, 115 mM sucrose, 4.2 mM trehalose, 5 mM HEPES, and 0.5 mM CaCl2 (unless otherwise noted), pH 7.2. Neuromuscular junction sharp electrode (electrode resistance between 20-30 MΩ) recordings were performed on muscles 6/7 of abdominal segments A2 and A3 in wandering third-instar larvae as described ^126^. Recordings were performed on a Leica microscope using a 10x objective and acquired using an Axoclamp 900A amplifier, Digidata 1440A acquisition system, and pClamp 10.7 software (Molecular Devices). Electrophysiological sweeps were digitized at 10 kHz and filtered at 1 kHz. Data were analyzed using Clampfit (Molecular Devices) and MiniAnalysis (Synaptosoft) software. Miniature excitatory postsynaptic potentials (mEPSPs) were recorded in the absence of any stimulation and motor axons were stimulated to elicit excitatory postsynaptic potentials (EPSPs).

### Larval crawling assay

Vials containing third instar larvae were analyzed for this assay. Vials were poured with 4 ml of 20% sucrose solution and left for 10 min to let the larvae float on top. Floating third instar animals were poured into a petri dish and washed gently twice with deionized water in a paintbrush. A minimum of 10 larvae of each genotype were analyzed on a 2% agarose gel in a petri dish with gridline markings 1 cm on graph paper. The larvae were acclimatized in the petri dish before videotaping. The average distance crawled (in centimeters) by larvae was calculated based on the average number of gridlines passed in 30 sec ^146^.

### Real-time PCR (RT PCR)

Total RNA was extracted from 10 larval muscle fillets or 20 larval brains for each experiment using TRIzol (GIBCO) according to the manufacturer’s instructions. Concentration and purity of total RNA were measured using a NanoDrop. 1 μg of total RNA was then subjected to DNA digestion using DNase I (Ambion). This was immediately followed by reverse transcription using the iScript Reverse Transcription Supermix (1708841, Bio-Rad). qPCR was performed using the StepOnePlus instrument (ThermoFisher Scientific) and SYBR Green Supermix (172-5270, Bio-Rad) by following the manufacturer’s instructions. The Step One Software analyzed the qPCR, and the relative expression level was presented as the ratio of the target gene to the internal standard gene, *rp49*. Each sample was analyzed in triplicate. At least three independent biological repeats were obtained for each experiment. The following primers were used for quantitative PCR (qPCR) analysis. For the *ND-20L* gene, forward: 5’ CATGCCGGTGTACGATTACC 3’ and reverse: 5’ CGTCCCCAGTTTAGCAGGTC 3’ primers are used. Similarly, for analyzing the *ND-20* transcript, forward: 5’ GAAGTGGCCCAAAATCTGCC 3’ and reverse: 5’ GAGCAGATCGTCCAGTCTGG 3’ primers were used respectively. All data for the RT PCR are presented as mean ± standard deviation.

### Seahorse analysis for oxygen consumption rate (OCR) measurement

The Seahorse analysis was carried out as previously described ^149^ with some modifications. Briefly, mitochondria were isolated from the thoracic muscles of 20 adult *Drosophila melanogaster* using a differential centrifugation method. Thoraces were homogenized in cold isolation buffer, followed by a 300 × g spin to remove debris and a subsequent 3,000 × g spin to pellet mitochondria. The final mitochondrial pellet was resuspended in mitochondrial isolation buffer, and protein concentration was quantified using a BCA assay. Isolated mitochondria from *Drosophila melanogaster* thoracic muscles were resuspended in Agilent Seahorse XF assay medium supplemented with glucose (10 mM), pyruvate (1 mM), and glutamine (2 mM), pH 7.4. For the Seahorse assay, ∼5–10 µg of isolated mitochondria were added to each well of a Seahorse XF Cell Culture Microplate. After centrifugation (3,000 × g for 20 min at 4°C), the plate was then loaded into the Seahorse XFeMini Analyzer, and the mitochondrial respiration profile was measured. The OCR data were normalized to protein concentration. All data for the Seahorse assay are presented as mean ± standard deviation.

### Rendering Schematic Cartoons, Figures, and Tables

For this study, the schematic cartoons for Figure 4 were drawn for this study using bioRender software. The schematic cartoons for Figure 5 were drawn Adobe Illustrator. All final Figures or Supplemental Figures were saved as TIF files and sized in Adobe Photoshop. Tables were generated in Microsoft Word.

### Quantification And Statistical Analysis Quantification of mitochondrial branch length

Controls and RNAi-depleted animals were analyzed using mitochondrial marker Mito-GFP, *D42* driver in cell bodies from larvae ventral nerve cord. All images were processed using image J software. Z-stacks of individual neurons were merged. The Mito-GFP signal was enhanced by adjusting brightness and contrast. The binary masks were created using Image>Adjust>Threshold., Method:Otsu, and Background:Dark. The branches were generated using Process>Binary>Skeletonised^150^. These skeletonized images were analyzed, and branch length of the individual cluster was manually calculated using image J tools.

### Imaging quantifications

Maximum intensity projections were used for quantitative image analysis with the Image J software (National Institutes of Health) analysis toolkit. Boutons from muscle 4 or type Ib terminal boutons on the muscle 6/7 of A2 hemi segment from at least six NMJ synapses were used for quantification using Image J software. Student’s t-test for pairwise and one-way ANOVA with post-hoc Tukey’s test for multiple comparisons was used for statistical analysis, using GraphPad Prism Software. Specific p-value and tests are noted in the figures and figure legends and supplementary files and shown in graphs as follows: * *p*<0.05, ** *p*<0.001, and *** *p*<0.0001. The data are presented as mean ± s.e.m. For quantification of Futsch loops, third instar larval preparations were double immunostained with HRP, 22C10 and images were captured in Zeiss LSM710 confocal microscope. Only NMJs of muscles 6/7 of A2 hemi segment were used for quantification. The images were digitally magnified using image J software and the total number of HRP-positive boutons was manually counted in each image. Futsch positive loops, which are co-localized with HRP, were included in this analysis. Images with incomplete loops and diffused staining were not included in the count ^151^.

### Electrophysiological analysis

Average mEPSP, EPSP, and quantal content were calculated for each genotype by dividing EPSP amplitude by mEPSP amplitude. Muscle input resistance (R_in_) and resting membrane potential (V_rest_) were monitored during each experiment. Recordings were rejected if the V_rest_ was above -60 mV, and R_in_ was less than 5 MΩ. Pharmacological agents were bath applied in recording saline at the final concentrations indicated in the text, figures, and tables. The agents included Xestospongin C (Abcam), Dantrolene (Tocris) and BAPTA-AM (Sigma Aldrich). Failure analysis was performed in HL3 solution containing 0.1 mM CaCl2, which resulted in failures in about half of the stimulated responses in wild-type larvae. A total of 30 trials (stimulations) were performed at each NMJ in all genotypes. The failure rate was obtained by dividing the total number of failures by the total number of trials (100). High-frequency (10 Hz) recordings were performed at a calcium concentration of 2 mM and paired-pulse recordings (10 Hz) were performed at calcium concentrations of 0.4 mM and 1.5 mM, respectively. Paired-pulse ratios were calculated as the EPSP amplitude of the second response divided by the first response (EPSP2/EPSP1).

## SUPPLEMENTAL INFORMATION

Document S1:

- Supplemental Figures S1-S18.
- Figure Legends for Supplemental Figures S1-S18. A list of the Supplemental Figure Legend headers is replicated here:
- **Figure S1, related to Figure 1**: Tissue-specific expression of the Mitochondrial Complex I subunit NDUFS7 and functional assessment of Mitochondrial respiration via Seahorse-based oxygen consumption rate analysis
- **Figure S2, related to Figure 2**: Loss of an MCI subunit triggers excess mitochondrial ROS accumulation in neurons and muscle
- **Figure S3, related to Figure 2**: Loss of *Sod2* induces excessive reactive oxygen species formation in neuronal and muscle tissues
- **Figure S4, related to Figure 2**: Misregulation of ROS in neurons affects mitochondria morphology and distribution in the ventral nerve cord and distal axons in third-instar larvae
- **Figure S5, related to Figure 2**: Both *UAS-Catalase* and *UAS-Sod1* overexpression constructs fail to rescue synaptic and mitochondrial phenotypes in *NDUFS7* RNAi-depleted animals
- **Figure S6, related to Figure 2**: Depleting a mitochondrial fusion gene induces excess mitochondrial ROS formation in neurons
- **Figure S7, related to Figure 2**: Loss of an MCI subunit in motor neurons causes abnormal Cysteine-String Protein (CSP) accumulation in distal axons
- **Figure S8, related to Figure 2**: Both *Catalase* and *Sod1* overexpression fail to reverse Cysteine-String Protein (CSP) accumulation in distal axons of *NDUFS7* RNAi-expressing animals
- **Figure S9, related to Figure 3**: Both *Catalase* and *Sod1* overexpression fail to restore cytoskeletal loops in *NDUFS7*-deficient animals
- **Figure S10, related to Figure 2**: *marf* is required presynaptically to maintain synaptic transmission at NMJs
- **Figure S11, related to Figure 4**: *NDUFS7* depletion in neurons differentially affects the failure rate in firing under very low calcium concentrations
- **Figure S12, related to Figure 4**: Rotenone incubation increases levels of active zone material at the NMJs
- **Figure S13, related to Figure 4**: Both *Catalase* and *Sod1* overexpression fail to reverse synaptic and mitochondrial phenotypes in *NDUFS7* RNAi-depleted animals
- **Figure S14, related to Figure 5**: Loss of MCI modulates activities of intracellular calcium channels and regulates levels of active zone materials to stabilize synaptic strength
- **Figure S15, related to Figure 6**: Combined genetic loss in neurons of an MCI subunit and glycolysis or TCA cycle genes impairs evoked neurotransmission
- **Figure S16, related to Figure 6**: Loss in neurons of glycolysis or TCA cycle genes reverses the active zone enhancements normally induced by *NDUFS7* loss
- **Figure S17, related to Figure 7**: *NDUFS7* subunit in muscle affects the organization of α-Spectrin scaffold at the NMJs
- **Figure S18, related to Figure 8**: Both *Catalase* and *Sod1* overexpression fail to restore defective glutamate receptor-active zone apposition in *NDUFS7*-deficient animals

Document S2:

- S1-S21 Tables: Summary data for graphs in all Figures and Supplemental Figures.
- S22 Table: Detailed information about reagents and materials for items described in the STAR METHODS section.

