## Supplemental Figures and Legends for "Mitochondrial Complex I and ROS control neuromuscular function through opposing pre- and postsynaptic mechanisms"

This is Supplemental Document S1 for Mallik *et al*. It contains the following:

- Figures S1-S18 and accompanying legends.

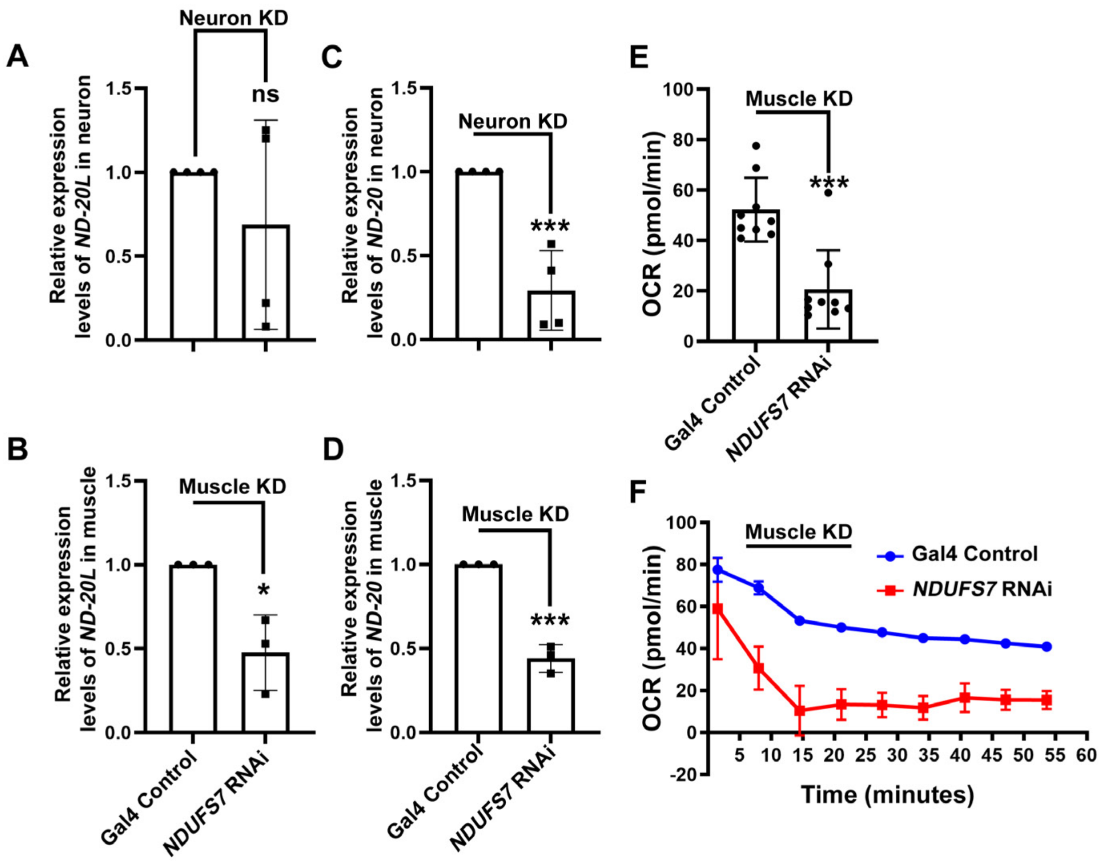


**Figure S1, related to Figure 1: Tissue-specific expression of the Mitochondrial Complex I subunit NDUFS7 and functional assessment of Mitochondrial respiration via Seahorse-based oxygen consumption rate analysis**

(A-B) Quantitative RT-PCR showing transcript levels of *ND-20L* in Gal4 controls and pan-neuron and muscle-Gal4 driven *UAS-NDUFS7*[RNAi]. Compared to pan-neuron Gal4 control (*elaV(C155)*-Gal4/+), *elaV(C155)*-Gal4 driven *UAS-NDUFS7*[RNAi] (*elaV(C155)-Gal4/+; NDUFS7[RNAi]/+*) led to ~25% reduction in *ND-20L* transcript level in neurons. The muscle Gal4-driven *UAS-NDUFS7[RNAi]* showed ~50% reduction in *ND-20L* transcript levels (*UAS-NDUFS7[RNAi]/+; BG57-Gal4/+*) compared to Gal4 control (*BG57-Gal4/+*) in the muscle. Error bars represent mean ± standard deviation *p*=0.354, **p*=0.015. Statistical analysis based on Student's t-test for pairwise comparisons. (C-D) Quantitative RT-PCR showing transcript levels of *ND-20* in Gal4 controls and pan-neuron and muscle-Gal4 driven *UAS-NDUFS7[RNAi]*. Compared to pan-neuron Gal4 control (*elaV(C155)*-Gal4/+), *elaV(C155)*-Gal4 driven *UAS-NDUFS7[RNAi]* (*elaV(C155)-Gal4/+; UAS-NDUFS7[RNAi]/+*) led to ~70% reduction in *ND-20* transcript level in neurons. The muscle Gal4-driven *UAS-NDUFS7[RNAi]* showed ~60% reduction in *ND-20* transcript levels (*UAS-NDUFS7[RNAi]/+; BG57-Gal4/+*) compared to Gal4 control (*BG57-Gal4/+*) in the muscle. Error bars represent mean ± standard deviation ****p*=0.001, ****p*=0.0003. Statistical analysis based on Student's t-test for pairwise comparisons. (E) Histogram showing oxygen consumption rate in the mitochondria isolated from the thoracic region in muscle Gal4 control and muscle-depleted *UAS-NDUFS7[RNAi]* animals. The muscle depletion of *NDUFS7* message resulted in a ~80% reduction in oxygen consumption compared to the Gal4 control. (F) Oxygen consumption rate in muscle Gal4 control and muscle-depleted *UAS-NDUFS7[RNAi]* are plotted on a 60-minute time scale. Raw data for this figure are available in the S2 Data Excel file, tab Figure_S1.

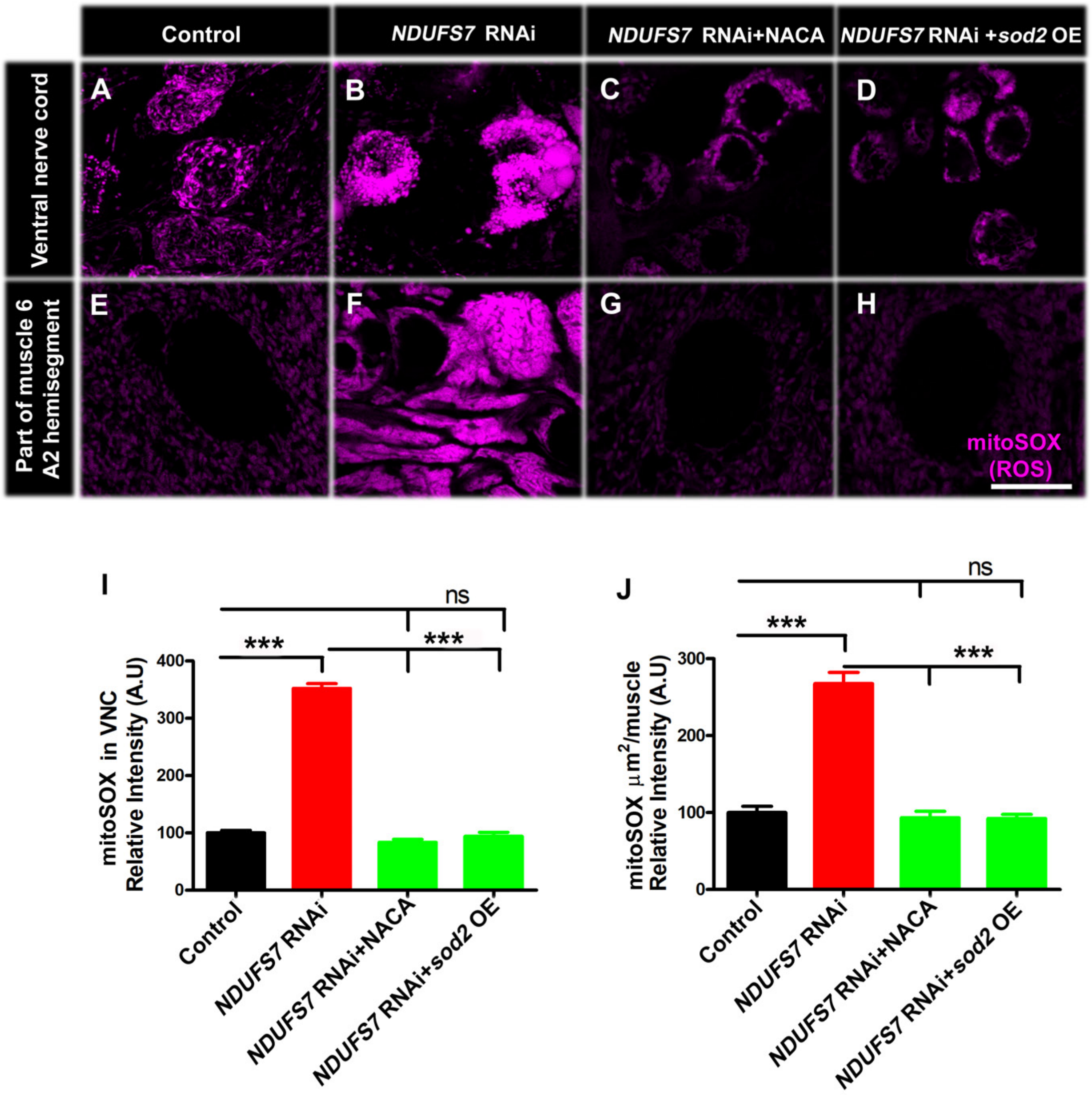


**Figure S2, related to Figure 2: Loss of an MCI subunit triggers excess mitochondrial ROS accumulation in neurons and muscle**

(A-D) Representative confocal images of the ventral nerve cord (VNC) at the third instar larval brain in (A) *mito-GFP, D42-Gal4/+* (B) *UAS-NDUFS7[RNAi]/+; mito-GFP, D42-Gal4/+*, (C) *UAS-NDUFS7[RNAi]/+; mito-GFP, D42-Gal4/+* NACA and (D) *UAS-NDUFS7[RNAi]/UAS-Sod2; mito-GFP, D42-Gal4/+* labeled with superoxide indicator mitoSOX (magenta) in live animals. *UAS-NDUFS7[RNAi]*-depleted larvae supplemented with NACA or co-expressing *Sod2* in neurons showed a significant reduction in mitochondrial ROS levels compared to the Gal4 control. (E-H) Confocal images of third instar larval body wall muscle 6 at A2 hemi segment in the indicated genotypes. Muscle-depleted *UAS-NDUFS7[RNAi]* restored ROS levels when the animals were supplemented with NACA or co-expression of *Sod2* transgene. Scale bar: 5 µm. (I-J) Histogram showing the relative intensity of mitoSOX in VNC and body wall muscle in (E) *UAS-mitoGFP/+; BG-57-Gal4/+*, (F) *UAS-NDUFS7[RNAi]/UAS-mitoGFP; BG-57-Gal4/+*, (G) *UAS-NDUFS7[RNAi]/UAS-mitoGFP; BG-57-Gal4/+* and (H) *UAS-NDUFS7[RNAi]/UAS-Sod2; BG-57-Gal4/+* labeled with superoxide indicator mitoSOX (magenta) in live animals. ****p*<0.0001; ns, not significant. Statistical analysis based on one-way ANOVA followed by post-hoc Tukey's multiple-comparison test. Error bars represent mean ± s.e.m. Raw data for this figure are available in the S2 Data Excel file, tab Figure_S2.

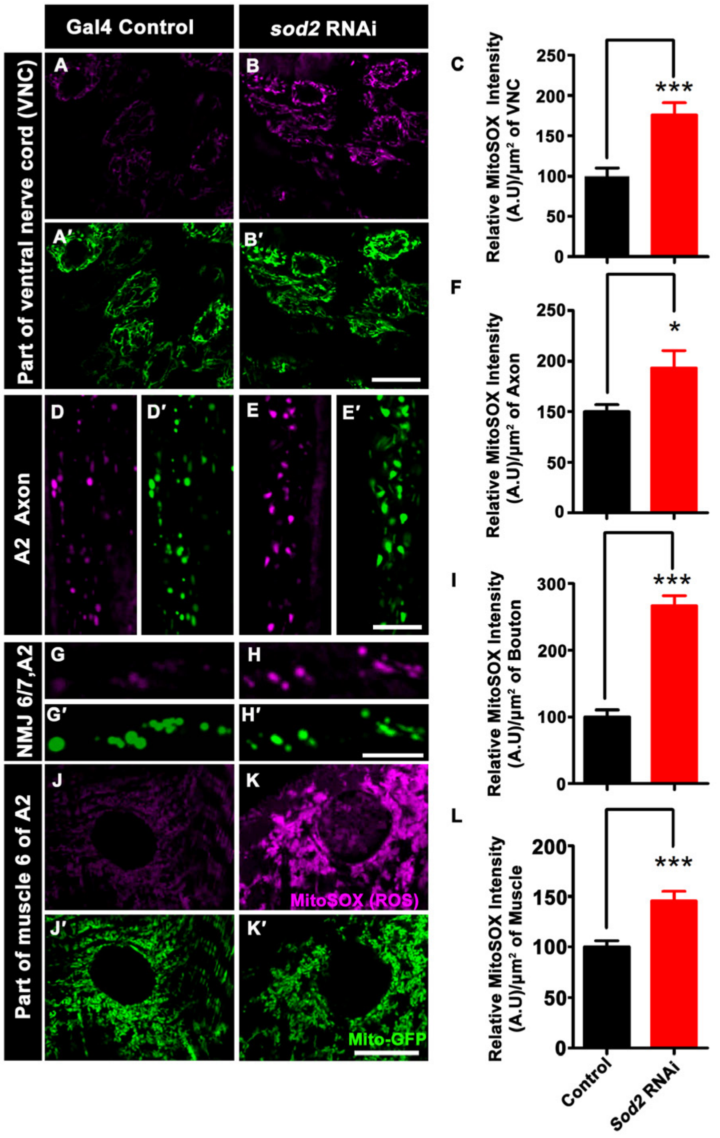


**Figure S3, related to Figure 2: Loss of *Sod2* induces excessive reactive oxygen species formation in neuronal and muscle tissues**

(A-A'- B-B ') Representative confocal images of the ventral nerve cord (VNC) at the third instar larval brain in (A-A') *UAS-mito-GFP, D42-Gal4/+* and (B-B') *UAS-mito-GFP, D42-Gal4/Sod2[RNAi]* labeled with superoxide indicator mitoSOX (magenta) and mito-GFP (green) in live animals. (C) Histogram showing the relative intensity of mitoSOX in VNC in the indicated genotypes. ***$p$=0.0003 (VNC: control vs *Sod2 [*RNAi]). (D-D'- E-E ') Representative confocal images of the axon at the third instar larval fillet in (D-D') *UAS-mito-GFP, D42-Gal4/+* and (E-E') *UAS-mito-GFP, D42-Gal4/Sod2 [RNAi]* labeled with superoxide indicator mitoSOX (magenta) and mitoGFP (green) in live animals. (F) Histogram showing the relative intensity of mitoSOX in axons in the indicated genotypes. *$p$=0.020 (Axon: control vs *Sod2* RNAi). (G-G'- H-H') Representative confocal images of the third instar bouton of A2 hemi segment in (G-G') *UAS-mito-GFP, D42-Gal4/+*, and (H-H') *UAS-mito-GFP, D42-Gal4/Sod2 [RNAi]* labeled with superoxide indicator mitoSOX (magenta) and mitoGFP (green) in live animals. (I) Histogram showing the relative intensity of mitoSOX at boutons in the indicated genotypes. ***$p$<0.0001 (boutons: control vs *Sod2* RNAi). (J-J'- K-K') Representative confocal images of the third instar 6/7 muscle of A2 hemi segment in (J-J') *UAS-mito-GFP/+; BG57-Gal4/+*, and (K-K') *UAS-mito-GFP/+; BG57-Gal4/Sod2 [RNAi]* labeled with superoxide indicator mitoSOX (magenta) and mitoGFP (green) in live animals. (L) Histogram showing the relative intensity of mitoSOX in 6/7 muscle in the indicated genotypes. ***$p$=0.0001 (6/7 muscles: control vs *Sod2 [*RNAi]). The depletion of *Sod2 [*RNAi] results in the abnormal accumulation of reactive oxygen species (ROS) in neurons and muscles. Scale bar: 10 µm (A-A'- E-E ') and 5 µm (G-G'- K-K '). Statistical analysis based on Student's t-test for pairwise comparison. Error bars represent mean ± s.e.m. Raw data for this figure are available in the S2 Data Excel file, tab Figure_S3.

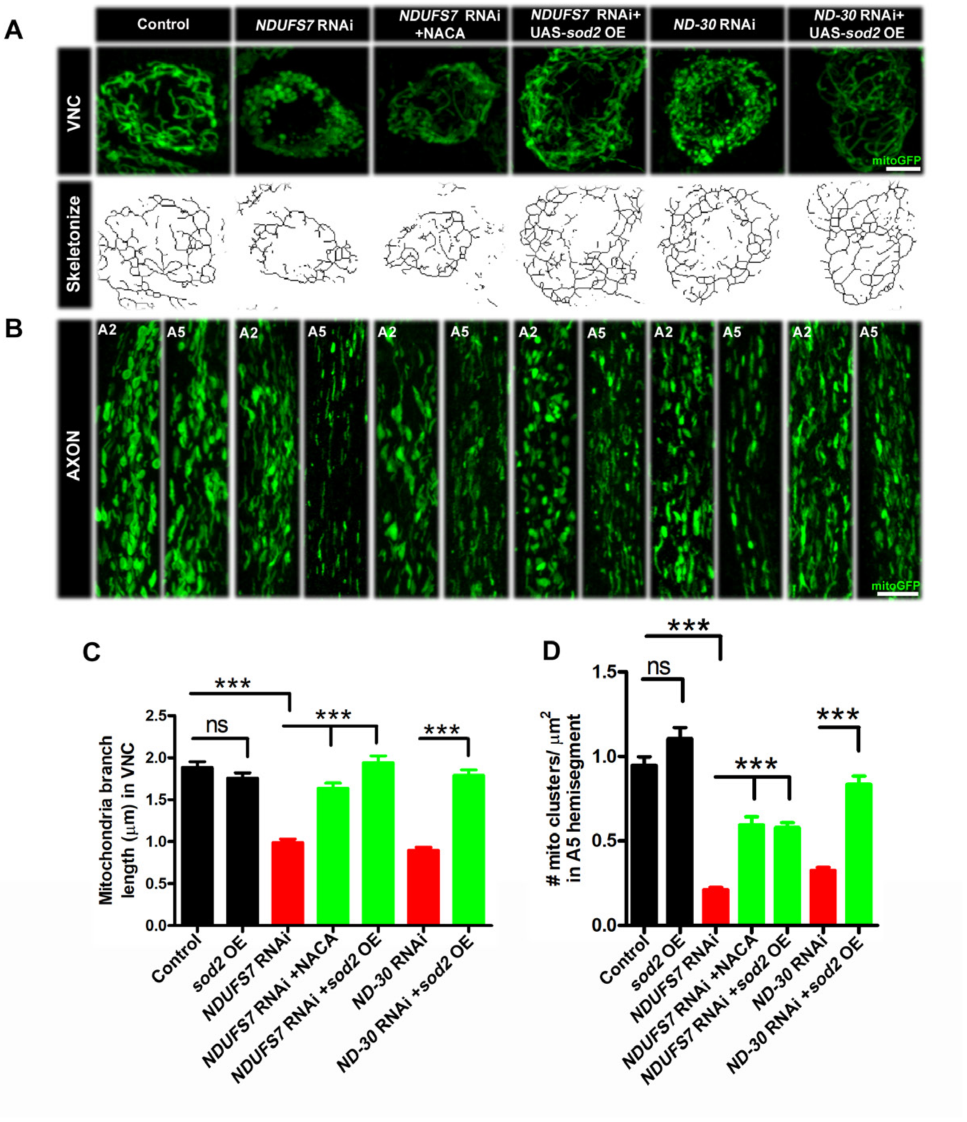


**Figure S4, related to Figure 2: Misregulation of ROS in neurons affects mitochondria morphology and distribution in the ventral nerve cord and distal axons in third-instar larvae.**

Mitochondrial morphology and distribution are affected in the ventral nerve cord and distal axons. *UAS-NDUFS7[RNAi], UAS-NDUFS7[RNAi]* co-expressing *UAS-Sod2*, *ND-30 RNAi* and *ND-30* RNAi co-expressing *UAS-Sod2* and controls were crossed to a motor neuron driver line (*D42-Gal4, UAS-mitoGFP*) to label neuronal mitochondria. (A) Ventral nerve cord (VNC): *UAS-mitoGFP* exhibits normal mitochondria organization, *UAS-NDUFS7[RNAi]* and *ND-30* RNAi exhibit clustered mitochondria; however, *UAS-NDUFS7[RNAi]* animals were raised in a media containing NACA or co-expression of *UAS-Sod2* in *UAS-NDUFS7[RNAi]* and *ND-30* RNAi depleted animals restore normal mitochondrial organization. The respective fluorescent images were skeletonized to measure mitochondrial branch length (organization). (B) Comparison of a proximal axonal segment in A2 and a distal segment in A5. Distal segments of A5 axons in *UAS-NDUFS7[RNAi]* contain fewer mitochondria than proximal segments. Scale bar: 5 µm. Mitochondrial distribution was significantly suppressed to control values when *UAS-NDUFS7[RNAi]* was raised in media containing NACA or genetically over-expressing *UAS-Sod2* in *UAS-NDUFS7[RNAi]* and *ND-30* RNAi in neurons. (C-D) Histogram showing mitochondrial branch length (µm) and number in the indicated genotypes. ****p*<0.0001, ns, not significant. Statistical analysis based on one-way ANOVA followed by post-hoc Tukey's multiple-comparison test. Error bars represent mean ± s.e.m. Raw data for this figure are available in the S2 Data Excel file, tab Figure_S4.

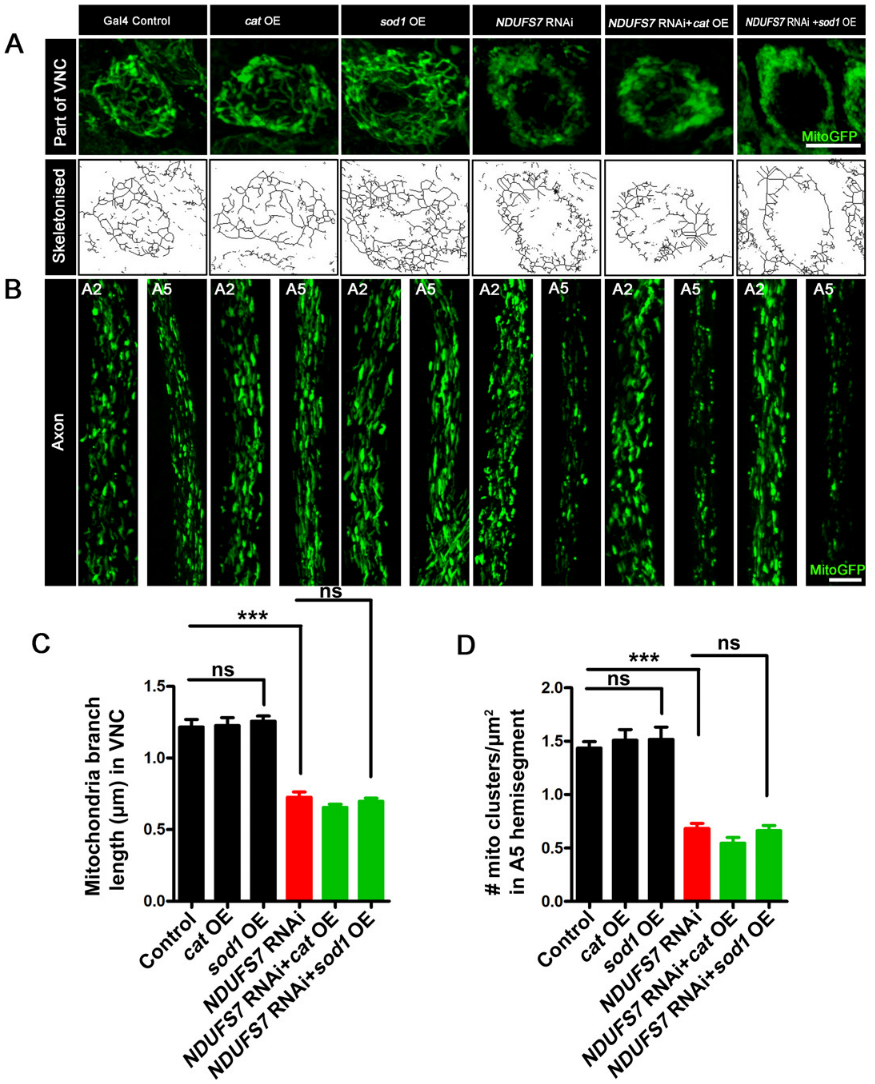


**Figure S5, related to Figure 2: Both *UAS-Catalase* and *UAS-Sod1* overexpression constructs fail to rescue synaptic and mitochondrial phenotypes in *NDUFS7* RNAi-depleted animals.** (A) Representative images of the A2 hemi segment of muscle 6/7 NMJs in *UAS-mitoGFP, D42-Gal4/*+, (B) *UAS-cat/+; UAS-mitoGFP, D42-Gal4/*+, (C) *UAS-sod1/+; UAS-mitoGFP, D42-Gal4/*+, (D) *NDUFS7[RNAi]/+; UAS-mitoGFP, D42-Gal4/*+, (E) *NDUFS7[RNAi]/UAS-cat; UAS-mitoGFP, D42-Gal4/*+ and (F) *NDUFS7[RNAi]/UAS-sod1; UAS-mitoGFP, D42-Gal4/*+ larvae immunostained with antibodies against the active zone scaffold Bruchpilot (BRP:fire-LuT) to label the active zones. BRP levels are upregulated at the NMJs in *NDUFS7[RNAi]*-depleted flies, while overexpression of ROS scavenger *cat* or *sod1* in the neuron fails to restore BRP to the control level. (A-F) Scale bar: 2.5 µm. (F-G) Histograms showing quantification of BRP intensity (F) and density (G) in µm$^2$ area of bouton at muscle 6/7 in the genotypes mentioned above. At least 8 NMJs of each genotype were used for quantification. ***$p$ <0.0001. Error bars denote mean ± s.e.m. Statistical analysis based on one-way ANOVA followed by post-hoc Tukey's multiple-comparison test. (H-P) Representative traces, quantification of mEPSPs, EPSPs and quantal content in the indicated genotypes. Scale bars for EPSPs (mEPSP) are x=50 ms (1000 ms) and y= 10 mV (1 mV). EPSP amplitudes were maintained in *NDUFS7[RNAi]*-depleted flies due to the induction of BRP; however, NMJs with *sod1* rescued *NDUFS7[RNAi]* in neurons showed diminished evoked release when compared with *NDUFS7[RNAi]*. Minimum 7 NMJs recordings of each genotype were used for quantification. mEPSP amplitude: **$p$=0.002 (Control vs UAS-*cat*), *$p$=0.009 (Control vs UAS-*sod1*), *$p$=0.018 (Control vs *NDUFS7* [RNAi]); Quantal content: **$p$=0.0009 (Control vs UAS-*cat*), *$p$=0.004 (Control vs UAS-*sod1*), *$p$=0.010 (Control vs *NDUFS7[RNAi]*); ns, not significant. Statistical analysis based on one-way ANOVA followed by posthoc Tukey's multiple-comparison test. Error bars denote the standard error of the mean. (Q) Representative images of the A2 hemi segment of muscle 6/7 NMJs in *UAS-mitoGFP, D42-Gal4/*+, (R) *UAS-cat/+; UAS-mitoGFP, D42-Gal4/*+, (S) *UAS-sod1/+; UAS-mitoGFP, D42-Gal4/*+, (T) *NDUFS7[RNAi]/+; UAS-mitoGFP, D42-Gal4/*+, (U) *NDUFS7[RNAi]/UAS-cat; UAS-mitoGFP, D42-Gal4/*+ and (V) *NDUFS7[RNAi]/UAS-sod1; UAS-mitoGFP, D42-Gal4/*+ larvae immunostained with antibodies against HRP(magenta) and GFP (mito-GFP:green) to label neurons and mitochondria. *NDUFS7*-depleted and *cat* or *sod1* rescued *NDUFS7[RNAi]* animals harbor fewer mitochondria at the terminals than control animals. (Q-V) Scale bar: 5 µm. (W) Histograms showing quantification of mitochondrial clusters in the above-indicated genotypes. ≥ 8 NMJs of each genotype were used for quantification. ***$p$<0.0001. Error bars represent mean ± s.e.m. Statistical analysis based on one-way ANOVA followed by post-hoc Tukey's multiple-comparison test. Raw data for this figure are available in the S2 Data Excel file, tab Figure_S5.

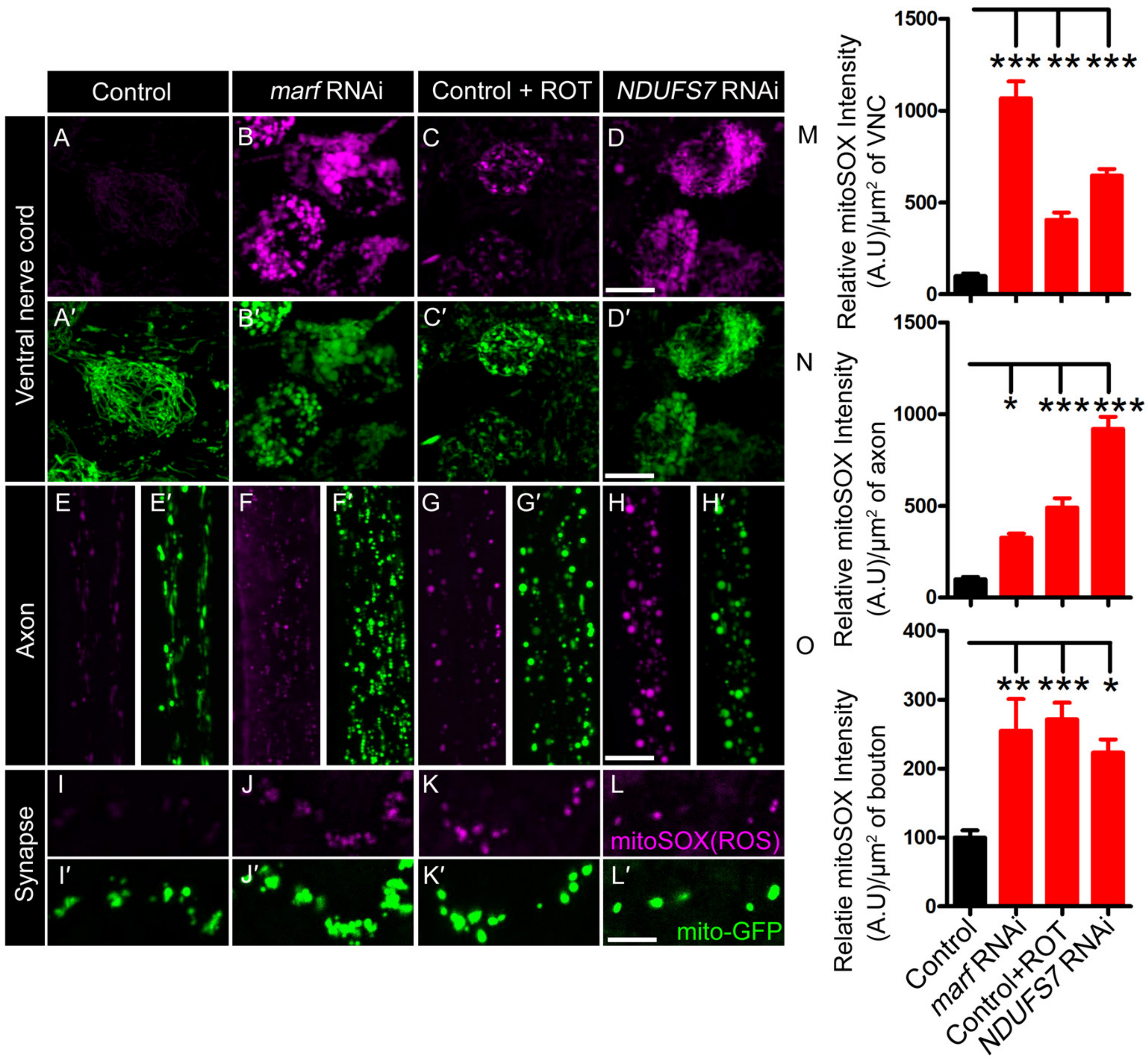


**Figure S6, related to Figure 2: Depleting a mitochondrial fusion gene induces excess mitochondrial ROS formation in neurons**

(A-A'- D-D ') Representative confocal images of the ventral nerve cord (VNC) at the third instar larval brain in (A-A') *mito-GFP, D42-Gal4/+,* (B-B') *mito-GFP, D42-Gal4/marf RNAi,* (C-C') *mito-GFP, D42-Gal4/+* with 25μM ROT and (D-D') *UAS-NDUFS7[RNAi]/+;mito-GFP, D42-Gal4/+* labeled with superoxide indicator mitoSOX (magenta) and mitoGFP (green) in live animals. (E-E'- H-H ') Representative confocal images of the axon at the third instar larval fillet in (E-E') *mito-GFP, D42-Gal4/+,* (F-F') *mito-GFP, D42-Gal4/marf RNAi,* (G-G') *mito-GFP, D42-Gal4/+* with 25μM ROT and (H-H') *UAS-NDUFS7[RNAi]/+;mito-GFP, D42-Gal4/+* labeled with superoxide indicator mitoSOX (magenta) and mitoGFP (green) in live animals. (I-I'- L-L'). Representative confocal images of the third instar bouton of A2 hemi segment in (I-I') *mito-GFP, D42-Gal4/+,* (J-J') *mito- GFP, D42-Gal4/marf RNAi,* (K-K') *mito-GFP, D42-Gal4/+* with 25μM ROT and (L-L') *UAS-NDUFS7[RNAi]/+;mito- GFP, D42-Gal4/+* labeled with superoxide indicator mitoSOX (magenta) and mitoGFP (green) in live animals. The depletion of *marf* RNAi or blocking MCI activity induces abnormal accumulation of ROS in neurons. Scale bar: 10 μm. (M-O) Histogram showing the relative intensity of mitoSOX in VNC, axon and boutons in the indicated genotypes ***$p$ <0.0001; **$p$ <0.001; **$p$=0.002 (Bouton: control vs marf RNAi), *$p$ <0.05. Statistical analysis based on one-way ANOVA followed by post-hoc Tukey's multiple-comparison test. Error bars represent mean ± s.e.m. Raw data for this figure are available in the S2 Data Excel file, tab Figure_S6.

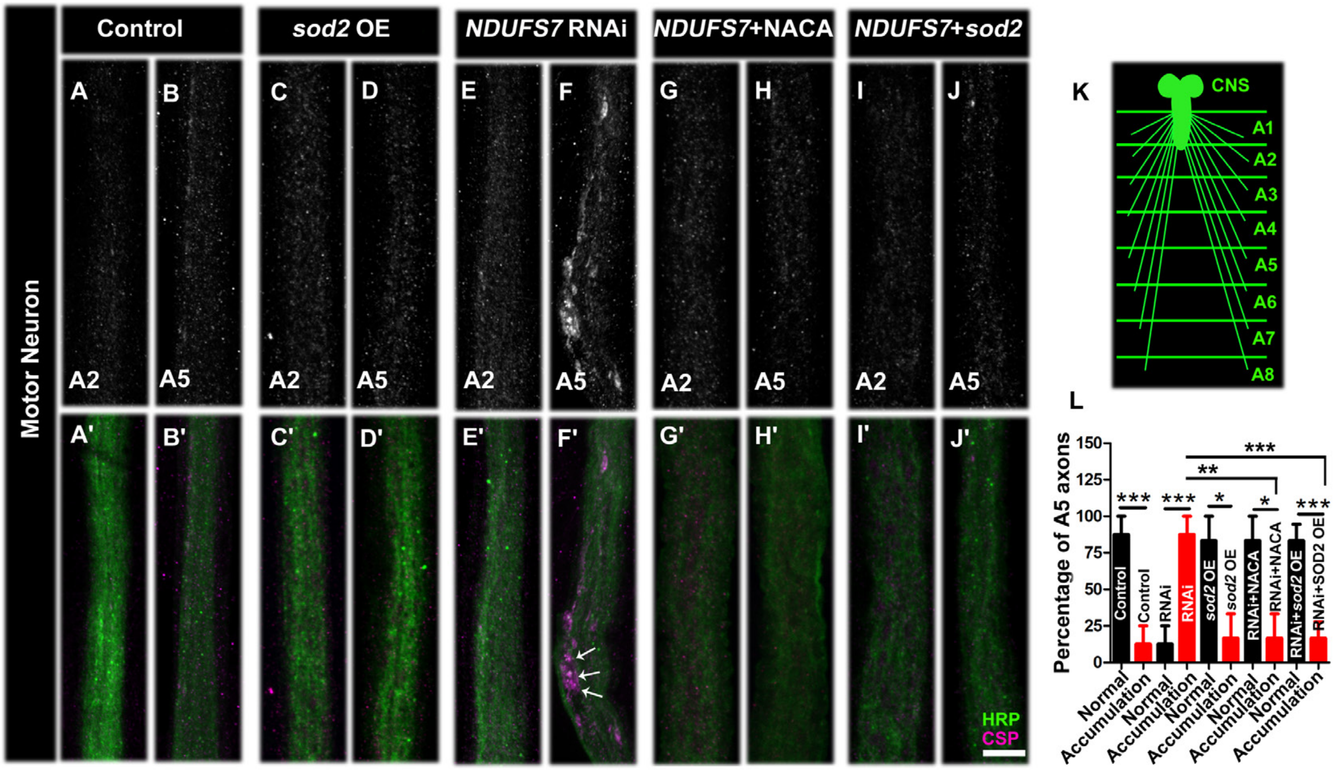


**Figure S7, related to Figure 2: Loss of an MCI subunit in motor neurons causes abnormal Cysteine-String Protein (CSP) accumulation in distal axons.**

Representative confocal images of the proximal (A2) and distal (A5) axons of larvae in (A-A′, B-B′) (Gal4 control), (C-C′, D-D′) (*UAS-Sod2/+; D42-Gal4/+*), (E-E′, F-F′) *D42*-Gal4 driven *UAS-NDUFS7[RNAi]* (*UAS-NDUFS7[RNAi]/+; D42-Gal4/+*), (G-G′, H-H′) *UAS-NDUFS7[RNAi]/+; D42-Gal4/+*NACA), and (I-I′ J-J′*)* (*UAS-NDUFS7[RNAi]/UAS-Sod2; D42-Gal4/+*) double immunolabeled with CSP (magenta) and HRP (green) antibodies. Motor neuron-depleted *UAS-NDUFS7[RNAi]* larvae showed abnormal accumulation of CSP in axons at lower hemi segment (A5) compared to Gal4 controls. Scale bar: 10 µm. CSP aggregates were cleared when *UAS-NDUFS7[RNAi]* animals were raised in media containing NACA or genetically expressing *UAS-Sod2* in neurons. (K) Schematic illustration showing VNC, axons and body wall muscle in a third instar larvae (L) Histogram showing the percentage of axons with abnormal accumulation in the indicated genotypes. ****p*=0.0008 (control: normal vs accumulation), **p*=0.0008 (*UAS-NDUFS7[RNAi]*: normal vs accumulation), **p*=0.0179 (*Sod2* OE: normal vs accumulation), **p*= 0.0179 (*UAS-NDUFS7[RNAi]* + NACA: normal vs accumulation), **p*=0.0004 (*UAS-NDUFS7[RNAi]*+Sod2 OE: normal vs accumulation), ***p*=0.0046 (*UAS-NDUFS7[RNAi]* vs *UAS-NDUFS7[RNAi]* + NACA: accumulation) and ****p*=0.0006 (*UAS-NDUFS7[RNAi]* vs *UAS-NDUFS7[RNAi]*+*Sod2* OE: accumulation). Statistical analysis was based on Fisher's exact test to differentiate two distinct phenotypes in the same sample. Error bars represent mean ± s.e.m. Raw data for this figure are available in the S2 Data Excel file, tab Figure_S7.

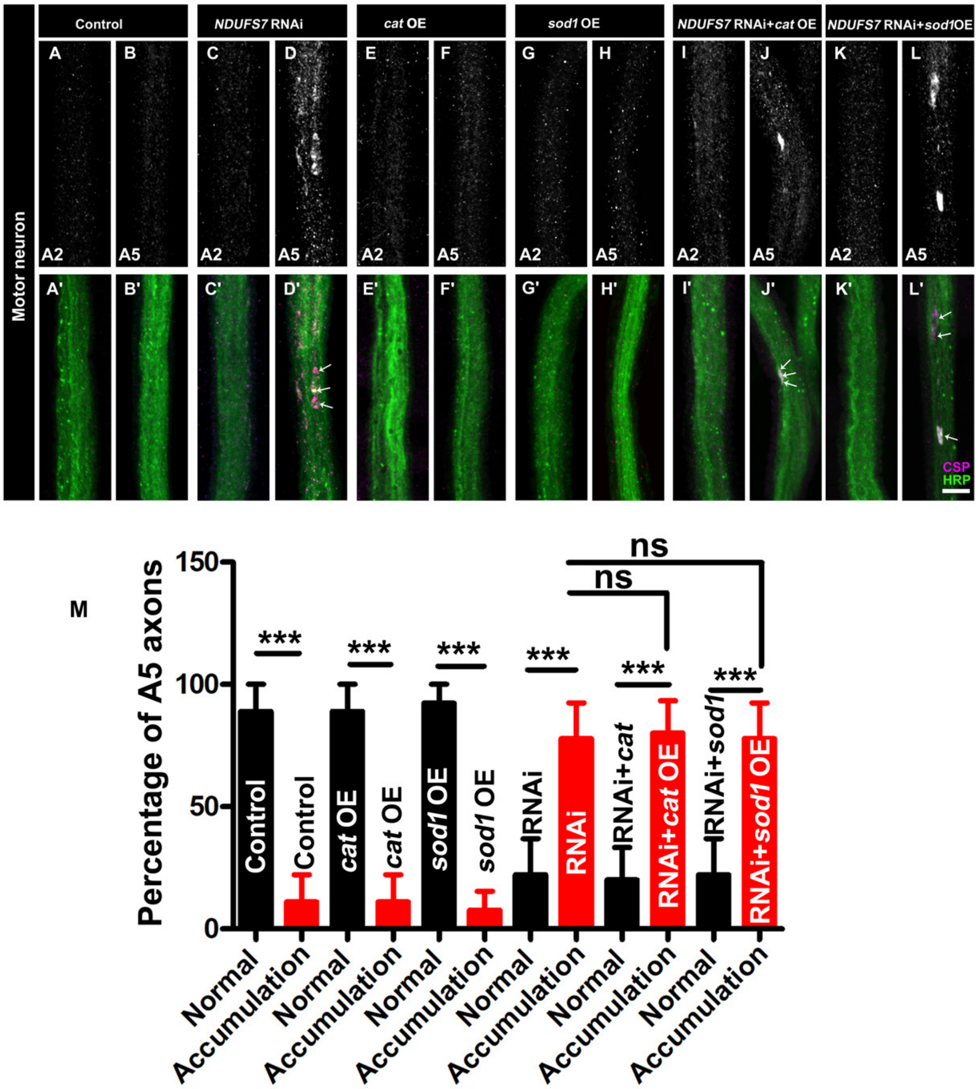


**Figure S8, related to Figure 2: Both *Catalase* and *Sod1* overexpression fail to reverse Cysteine-String Protein (CSP) accumulation in distal axons of *NDUFS7* RNAi-expressing animals**

Representative confocal images of the proximal (A2) and distal (A5) axons of larvae in (A-A′, B-B′) (Gal4 control: *UAS-mitoGFP, D42-Gal4*), (C-C′, D-D′) *UAS-mitoGFP, D42-Gal4* driven *NDUFS7* [RNAi] (*NDUFS7 [RNAi]/+; UAS-mitoGFP, D42-Gal4/+*), (E-E′, F-F′) (*UAS-cat/+; UAS-mitoGFP, D42-Gal4/+*), (G-G′, H-H′) (*UAS-sod1/+; UAS-mitoGFP, D42-Gal4/+*), (I-I′, J-J′) (*NDUFS7 [RNAi]/UAS-cat; UAS-mitoGFP, D42- Gal4/+*) and (J-J′ K-K′) (*NDUFS7 [RNAi]/UAS-sod1; UAS-mitoGFP, D42- Gal4/+*) double immunolabeled with CSP (magenta) and HRP (green) antibodies. Motor neuron-depleted *NDUFS7* [RNAi] larvae showed abnormal accumulation of CSP in axons at the lower hemi segment (A5) compared to Gal4 controls—scale bar: 10 µm. CSP aggregates were not cleared when *NDUFS7* RNAi animals were genetically expressing *UAS-cat* or *UAS-sod2* in neurons. (M) Histogram showing the percentage of axons with abnormal accumulation in the indicated genotypes. ***$p$<0.0001 (control: normal vs accumulation), ***$p$<0.0001 (*NDUFS7 [*RNAi]: normal vs accumulation), ***$p$<0.0001 (*cat* OE: normal vs accumulation), ***$p$<0.0001 (*sod1* OE: normal vs accumulation), ***$p$<0.0001 (*NDUFS7*[RNAi]+*cat* OE: normal vs accumulation), ***$p$<0.0001 (*NDUFS7* RNAi+*sod1* OE: normal vs accumulation), $p$=0.911 (*NDUFS7 [*RNAi] vs *NDUFS7 [*RNAi]+*cat* OE: accumulation) and $p$=1.00 (*NDUFS7* [RNAi] vs *NDUFS7* [RNAi]+*sod2* OE: accumulation). Statistical analysis was based on Fisher's exact test to differentiate two distinct phenotypes in the same sample. Error bars represent mean ± s.e.m. Raw data for this figure are available in the S2 Data Excel file, tab Figure_S8.

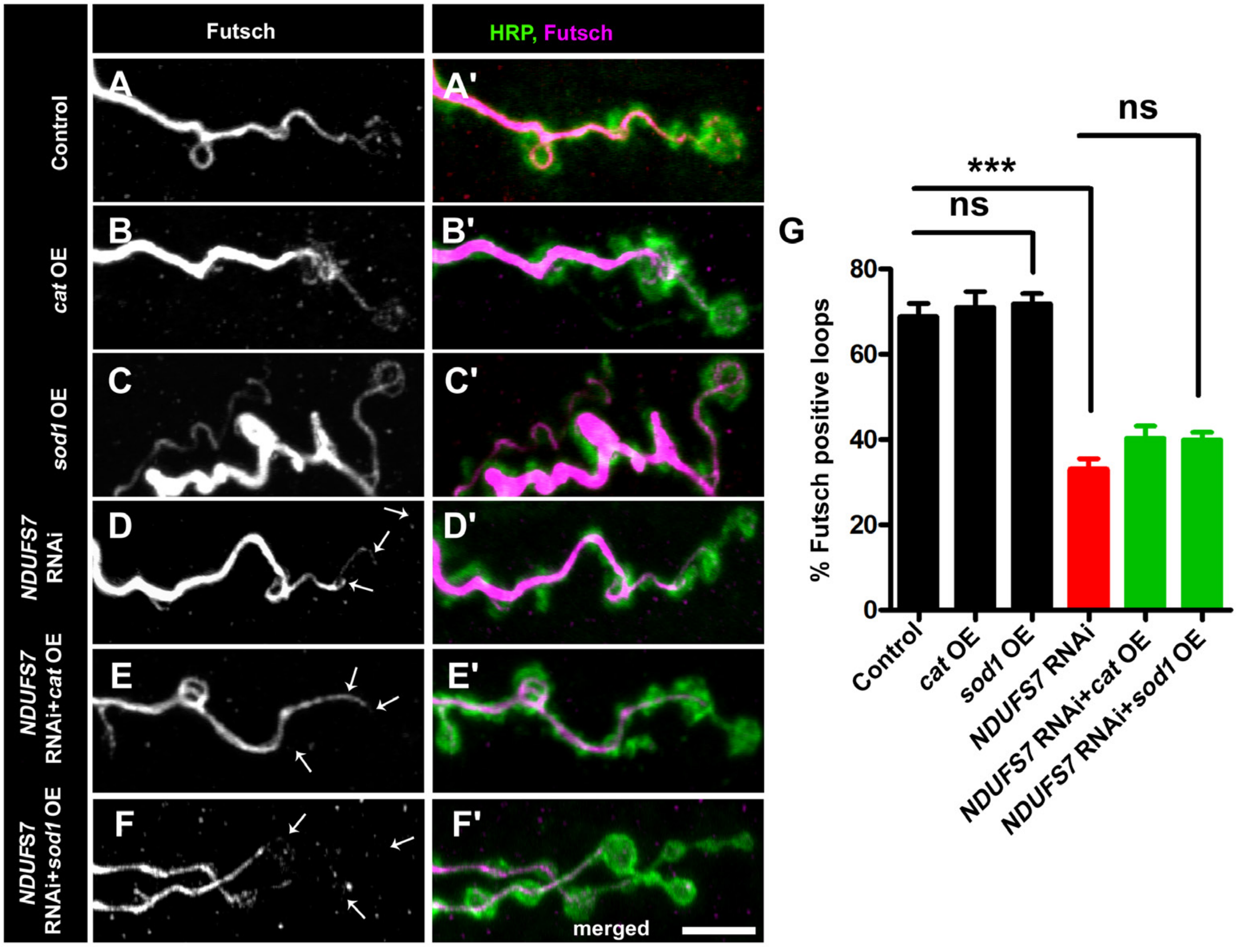


**Figure S9, related to Figure 3: Both *Catalase* and *Sod1* overexpression fail to restore cytoskeletal loops in *NDUFS7*-deficient animals**

Representative confocal images of NMJ synapses at muscle 6/7 of (A-A′) *UAS-mitoGFP, D42-Gal4* control, (B-B′) *UAS-cat* overexpression (*UAS-cat/+; UAS-mitoGFP, D42-Gal4/+*), (C-C′) UAS-*sod1* overexpression (*UAS-cat/+; UAS-mitoGFP, D42-Gal4/+*), (D-D′) *UAS-mitoGFP, D42-Gal4* driven *NDUFS7* [RNAi] (*NDUFS7 [RNAi] /+; UAS-mitoGFP, D42-Gal4/+*), (E- E′) *NDUFS7* knockdown with UAS-*cat* (*UAS-NDUFS7 [RNAi]/UAS-cat; UAS-mitoGFP, D42-Gal4/+*) and (F- F′) *NDUFS7* knockdown with UAS-*sod1* (*UAS-NDUFS7 [RNAi]/UAS-sod1; UAS-mitoGFP, D42-Gal4/+*). Each condition was double immunolabeled with 22C10 (anti-Futsch, magenta) and anti-HRP (green) antibodies. The motor neuron-depleted *NDUFS7* [RNAi] larvae showed a decrease in the number of Futsch-positive loops as compared to the Gal4 control. However, overexpression of either UAS-*cat* or UAS-*sod1* did not suppress Futsch-positive loops in the UAS-*NDUFS7* [RNAi] background. Scale bar: 5 µm. (G) Histograms showing the percentage of Futsch-positive loops in the indicated genotypes. $p$=0.659 (Control vs UAS-*cat*), $p$=0.505 (Control vs UAS-*sod1*), $p$<0.0001 (Control vs UAS-*NDUFS7 [RNAi],* $p$=0.071 (UAS-*NDUFS7 [RNAi]* vs UAS-*NDUFS7 [RNAi]*+UAS-*cat* and $p$=0.039 (UAS-*NDUFS7 [RNAi]* vs UAS-*NDUFS7 [RNAi]*+UAS-*sod1*. Statistical analysis based on one-way ANOVA followed by post-hoc Tukey's multiple-comparison test. Error bars represent mean ± s.e.m. Raw data for this figure are available in the S2 Data Excel file, tab Figure_S9.

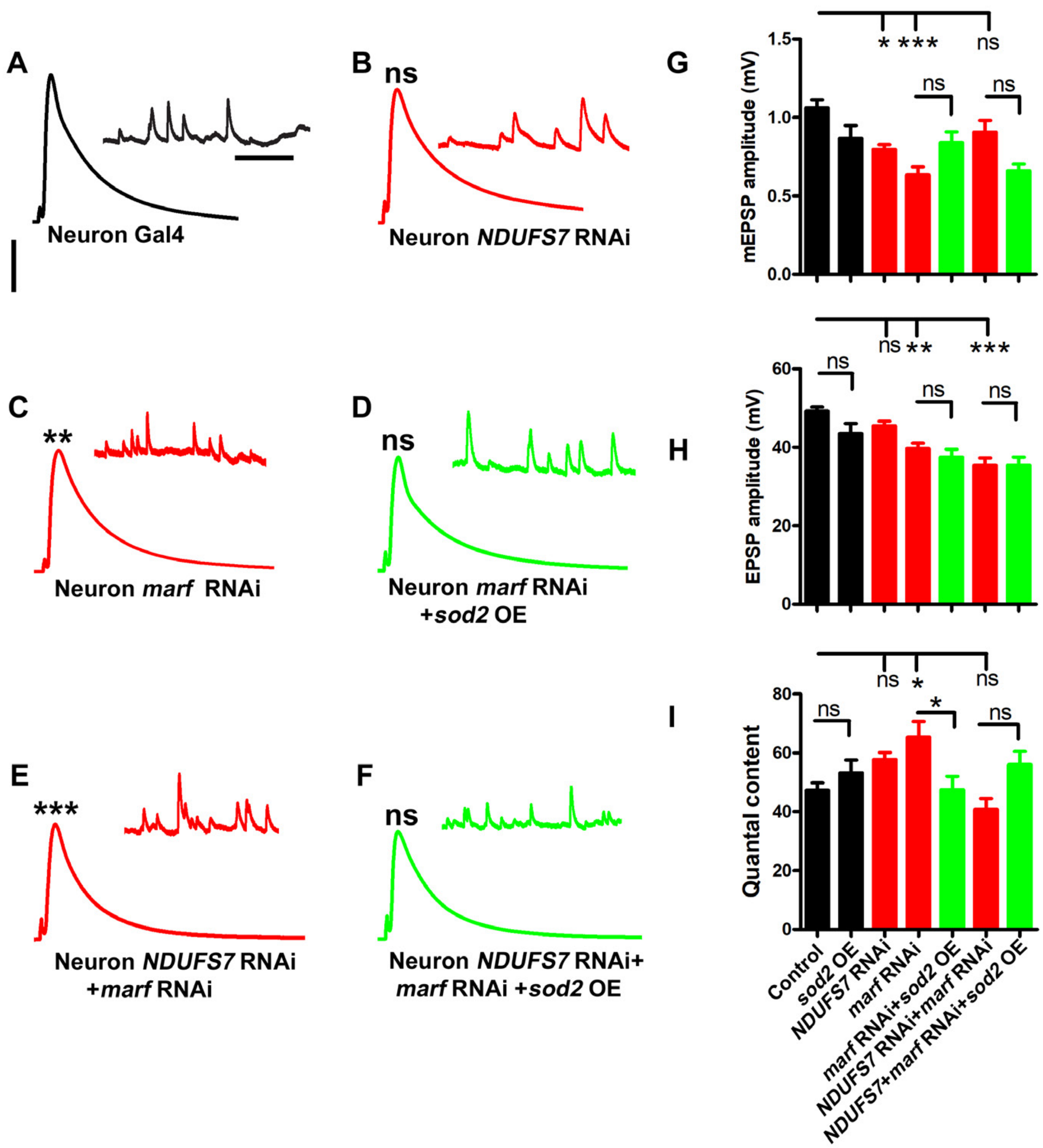


**Figure S10, related to Figure 2: *marf* is required presynaptically to maintain synaptic transmission at NMJs**

(A-F) The above figure shows representative traces of mEPSPs and EPSPs in (A) motor-neuron Gal4 control (*D42-Gal4/*+), (B) motor neuron Gal4 driven *UAS-NDUFS7[RNAi]* (*UAS-NDUFS7[RNAi]/+; D42-Gal4/*+), (C) motor neuron Gal4 driven *marf* RNAi (*marf RNAi/D42-Gal4*),(D) motor neuron Gal4 driven *marf* RNAi + *UAS-Sod2 OE* (*UAS-Sod2 OE/*+*; marf RNAi/*+), (E) motor neuron Gal4 driven *UAS-NDUFS7[RNAi]* + *marf* RNAi (*UAS-NDUFS7[RNAi]/*+*; marf RNAi/*+) and (F) motor neuron Gal4 driven *UAS-NDUFS7[RNAi]* + *marf* RNAi + *UAS-Sod2 OE* (*UAS-NDUFS7[RNAi]/*+*;marf RNAi/UAS-Sod2 OE*). Scale bars for EPSPs (mEPSP) are x=50 ms (1000 ms) and y= 10 mV (1 mV). Note that mEPSPs and EPSPs amplitudes were reduced in motor neuron Gal4-driven *marf* RNAi and *marf* RNAi+ *UAS-NDUFS7[RNAi]* animals. (G-I) Histograms showing average mEPSPs, EPSPs amplitude, and quantal content in the indicated genotypes. A minimum 8 NMJ recordings of each genotype were used for quantification. *$p$ <0.05 (mEPSP amplitude: control vs *UAS-NDUFS7[RNAi]*),*$p$=0.007 (QC: control vs marf RNAi),*$p$=0.026 (QC: *marf* RNAi vs *marf* RNAi+*Sod2* OE), **$p$<0.001, ***$p$<0.0001, ns, not significant. Statistical analysis based on one-way ANOVA followed by post-hoc Tukey's multiple-comparison test. Error bars represent mean ± s.e.m. Raw data for this figure are available in the S2 Data Excel file, tab Figure_S10.

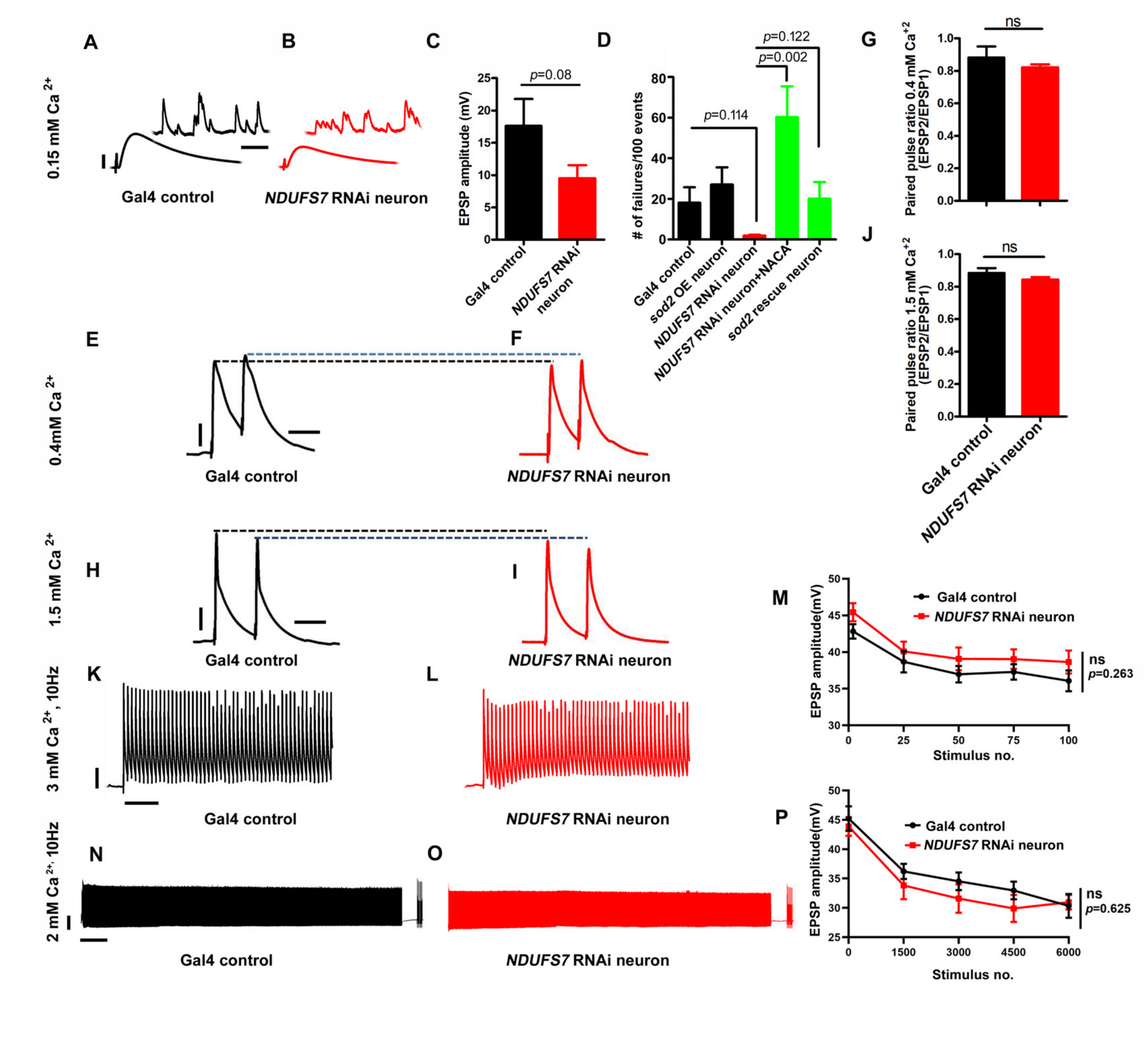


**Figure S11, related to Figure 4: *NDUFS7* depletion in neurons differentially affects the failure rate in firing under very low calcium concentrations.**

(A-B) Representative traces of EPSPs and mEPSPs in (A) motor neuron-Gal4 control (*D42-Gal4/*+) and (B) motor neuron-Gal4 driven *UAS-NDUFS7[RNAi]* (*UAS-NDUFS7[RNAi]/+; D42-Gal4/*+) at 0.15 mM extracellular $Ca^{2+}$ concentration. Scale bars for EPSPs (mEPSP) are x=50 ms (1000 ms) and y= 10 mv (1 mV). (C) Quantification of EPSPs in the indicated genotypes. Note that a very low extracellular calcium concentration mildly affects EPSPs in *NDUFS7[RNAi]* compared to control larvae. (D) Quantification of failure analysis at 0.1 mM $Ca^{2+}$ in Gal4 control, *UAS-Sod2/+; D42-Gal4/+, D42*-Gal4 driven *UAS-NDUFS7[RNAi]* (*UAS-NDUFS7[RNAi]/+; D42-Gal4/*+), *UAS-NDUFS7[RNAi]/+; D42-Gal4/*+ + NACA and *UAS-NDUFS7[RNAi]/UAS-Sod2; D42-Gal4/*+ animals. The number of failures were counted per hundred trials in each genotype. Failure analysis reveals a significant change in failure rate in motor neurons depleted *UAS-NDUFS7[RNAi]* as compared to control animals. The synaptic failure rates were restored to baseline level when *UAS-NDUFS7[RNAi]* depleted flies were reared in the media containing NACA or genetically expressing *Sod2* in motor neurons. (E-F) Representative paired-pulse EPSP traces at 0.4 mM extracellular $Ca^{2+}$ in the indicated genotypes. Scale bars for EPSPs are x=50 ms and y= 10 mV. No change in paired-pulse ratio was observed in motor neuron-depleted *UAS-NDUFS7[RNAi]* animals at 0.4 mM $Ca^{+2}$.

(G) Quantification of paired-pulse ratio (EPSP2/EPSP1) in motor neuron-Gal4 control (*D42-Gal4/+*) and (B) motor neuron-Gal4 driven *UAS-NDUFS7[RNAi]* (*UAS-NDUFS7[RNAi]/+; D42-Gal4/+*) larvae. (H-I) Representative paired-pulse EPSP traces at 1.5 mM extracellular $Ca^{2+}$ in the indicated genotypes. Scale bars for EPSPs are x=50 ms and y= 10 mV. No change in paired-pulse ratio was observed in motor neuron-depleted *UAS-NDUFS7[RNAi]* animals, even at higher calcium concentrations. (J) Quantification of paired ratio (EPSP2/EPSP1) in motor neuron-Gal4 control (*D42-Gal4/+*) and (B) motor neuron-Gal4 driven *UAS-NDUFS7[RNAi]* (*UAS-NDUFS7[RNAi]/+; D42-Gal4/+*) larvae at higher extracellular calcium. (K-L) Representative EPSP recordings of 100 stimuli at 3 mM extracellular $Ca^{2+}$ during a 10 Hz stimulus train in the indicated genotypes. Scale bars for EPSPs are x=1000 ms and y= 10 mV. (M) Quantifying EPSP amplitudes in the indicated stimulus number in the genotypes mentioned above. (N-O) Representative high-frequency recordings of 6000 stimuli at 2 mM extracellular $Ca^{2+}$ during a 10 Hz stimulus train in the indicated genotypes. Scale bars for EPSPs are x=1000 ms and y= 10 mV. (P) Quantifying EPSP amplitudes in the indicated stimulus number in the genotypes mentioned above. No significant depletion in vesicle pools was observed when *UAS-NDUFS7[RNAi]*-depleted animals were subjected to high-frequency nerve stimulation for 10 minutes. *p*-values are indicated in the figure, ns; they are not significant. Statistical analysis based on Student's t-test. Error bars signify mean ± s.e.m. Raw data for this figure are available in the S2 Data Excel file, tab Figure_S11.

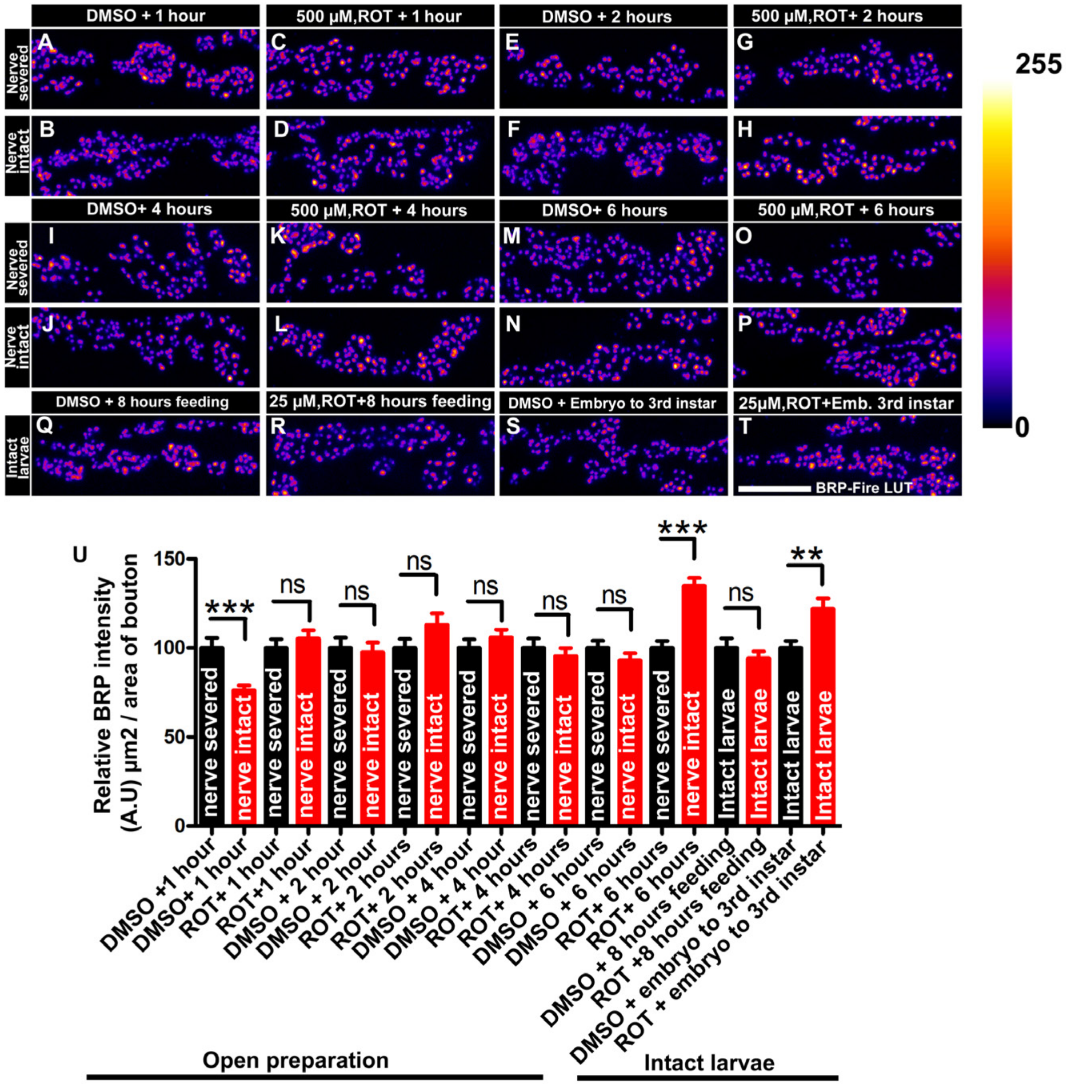


**Figure S12, related to Figure 4: Rotenone incubation increases levels of active zone material at the NMJs**

(A) Representative images of the A2 hemi segment of muscle 6/7 NMJs in (A) (*w*$^{1118}$+DMSO, 1 hour, nerve severed), (B) (*w*$^{1118}$ + DMSO, 1 hour, nerve intact), (C) (*w*$^{1118}$+500 μM ROT, 1 hour, nerve severed), (D) (*w*$^{1118}$+ROT, 1 hour, nerve intact), (E) (*w*$^{1118}$+DMSO, 2 hours, nerve severed), (F) (*w*$^{1118}$+DMSO, 2 hours, nerve intact), (G) (*w*$^{1118}$+500 μM ROT, 2 hours, nerve severed), (H) (*w*$^{1118}$+ROT, 2 hours, nerve intact), (I) (*w*$^{1118}$+DMSO, 4 hours, nerve severed), (J) (*w*$^{1118}$+DMSO, 4 hours, nerve intact), (K) (*w*$^{1118}$+500 μM ROT, 4 hours, nerve severed), (L) (*w*$^{1118}$+ROT, 4 hours, nerve intact), (M) (*w*$^{1118}$+DMSO, 6 hours, nerve severed), (N) (*w*$^{1118}$+DMSO, 6 hours, nerve intact), (O) (*w*$^{1118}$ +500 μM ROT, 6 hours, nerve severed), (P) (*w*$^{1118}$+500 μM ROT, 6 hours, nerve intact), (Q) (*w*$^{1118}$ + DMSO, 8 hours feeding, intact larvae), (R) (*w*$^{1118}$+25 μM ROT, 8 hours feeding, intact larvae), (S) (*w*$^{1118}$ +DMSO, embryo to 3$^{rd}$ instar) and (T) (*w*$^{1118}$+25 μM ROT, embryo to 3$^{rd}$ instar) larvae immunostained with antibodies against the active zone scaffold Bruchpilot (BRP:fire-LuT) to label the active zones. Scale bar: 5 μm. Note that the incubation or feeding larvae with rotenone elevate BRP levels at the NMJs in *w*$^{1118}$ +500 μM ROT, 6 hours nerve intact and *w*$^{1118}$ +25 μM ROT, embryo to 3$^{rd}$ instar compared to control nerve severed larvae. (U) Histograms show the quantification of BRP intensity in μm$^2$ of bouton at muscle 6/7 in the indicated genotypes. At least 8 NMJs of each genotype were used for quantification. ***$p$=0.0003 (*w*$^{1118}$+DMSO, 1 hour, nerve severed vs *w*$^{1118}$ + DMSO, 1 hour, nerve intact), ***$p$ <0.0001, **$p$=0.0019 (*w*$^{1118}$ +DMSO, embryo to 3$^{rd}$ instar vs *w*$^{1118}$+25 μM ROT, embryo to 3$^{rd}$ instar). Error bars signify the standard error of the mean. Statistical analysis is based on the Student's t-test for pairwise comparison among the samples. Raw data for this figure are available in the S2 Data Excel file, tab Figure_S12.

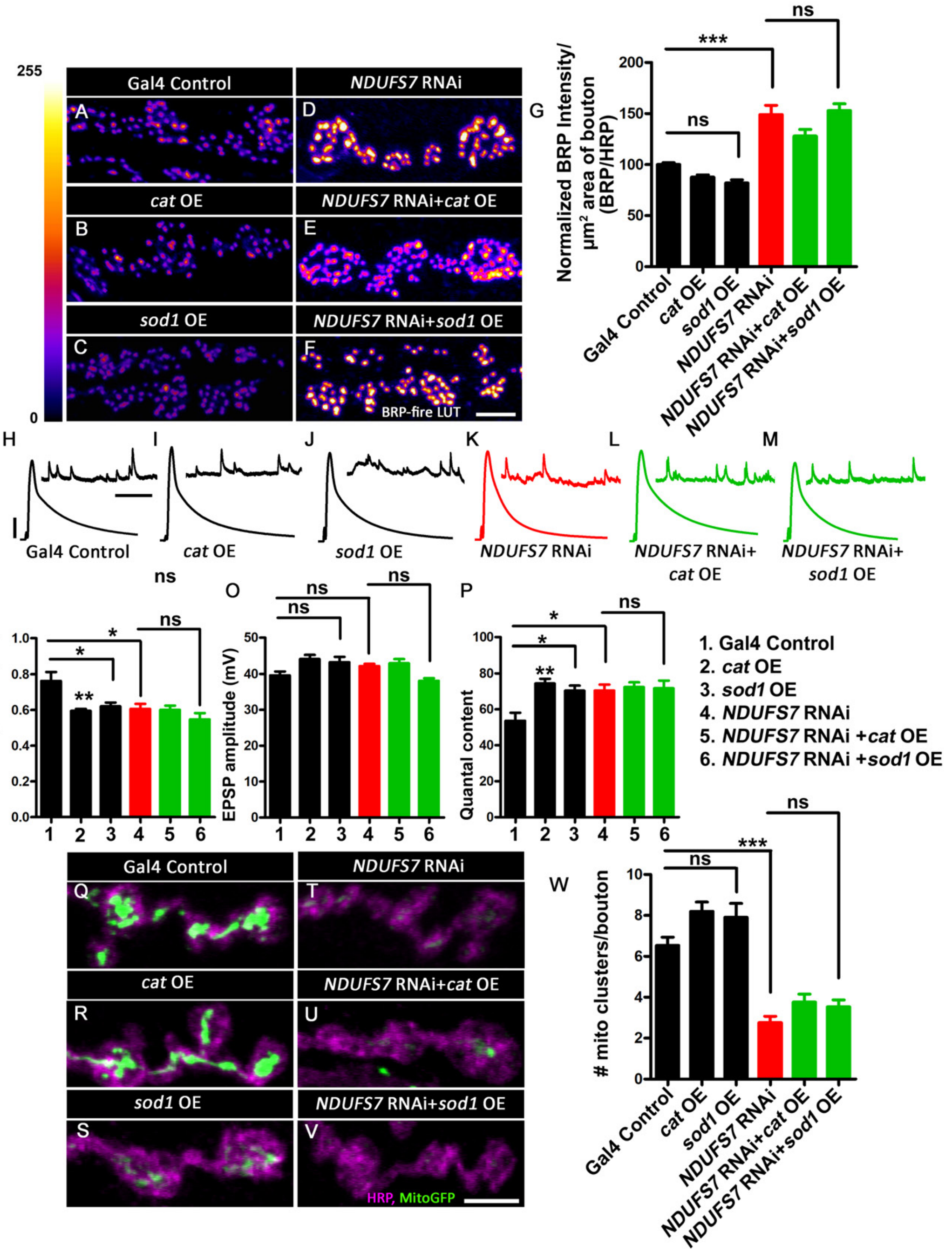


**Figure S13, related to Figure 4: Both *Catalase* and *Sod1* overexpression fail to reverse synaptic and mitochondrial phenotypes in *NDUFS7* RNAi-depleted animals.**

(A) Representative images of the A2 hemi segment of muscle 6/7 NMJs in *UAS-mitoGFP, D42-Gal4/*+, (B) *UAS-cat/+; UAS-mitoGFP, D42-Gal4/*+, (C) *UAS-sod1/+; UAS-mitoGFP, D42-Gal4/*+, (D) *NDUFS7 [RNAi]/+; UAS-mitoGFP, D42-Gal4/*+, (E) *NDUFS7 [RNAi]/UAS-cat; UAS-mitoGFP, D42-Gal4/*+ and (F) *NDUFS7 [RNAi]/UAS-sod1; UAS-mitoGFP, D42-Gal4/*+ larvae immunostained with antibodies against the active zone scaffold Bruchpilot (BRP:fire-LuT) to label the active zones. BRP levels are upregulated at the NMJs in *NDUFS7* [RNAi] depleted flies, while overexpression of ROS scavenger *cat* or *sod1* in the neuron fails to restore BRP to the control level. (A-F) Scale bar: 2.5 µm. (F-G) Histograms showing quantification of BRP intensity (F) and density (G) in µm$^2$ area of bouton at muscle 6/7 in the genotypes mentioned above. At least 8 NMJs of each genotype were used for quantification. ****p* <0.0001. Error bars denote mean ± s.e.m. Statistical analysis based on one-way ANOVA followed by post-hoc Tukey's multiple-comparison test. (H-P) Representative traces, quantification of mEPSPs, EPSPs and quantal content in the indicated genotypes. Scale bars for EPSPs (mEPSP) are x=50 ms (1000 ms) and y= 10 mV (1 mV). EPSP amplitudes were maintained in *NDUFS7* [RNAi]-depleted flies due to the induction of BRP; however, NMJs with *sod1* rescued *NDUFS7* [RNAi] in neurons showed diminished evoked release when compared with *NDUFS7* [RNAi]. Minimum 7 NMJs recordings of each genotype were used for quantification. mEPSP amplitude: ***p*=0.002 (Control vs UAS-*cat),* **p*=0.009 *(*Control vs UAS-*sod1*), **p*=0.018 (Control vs *NDUFS7* [RNAi]); Quantal content: ***p*=0.0009 (Control vs UAS-*cat),* **p*=0.004 *(*Control vs UAS-*sod1*), **p*=0.010 (Control vs *NDUFS7* [RNAi]); ns, not significant. Statistical analysis based on one-way ANOVA followed by posthoc Tukey's multiple-comparison test. Error bars denote the standard error of the mean. (Q) Representative images of the A2 hemi segment of muscle 6/7 NMJs in *UAS-mitoGFP, D42-Gal4/*+, (R) *UAS-cat/*+*; UAS-mitoGFP, D42-Gal4/*+, (S) *UAS-sod1/*+*; UAS-mitoGFP, D42-Gal4/*+, (T) *NDUFS7 [RNAi]/*+*; UAS-mitoGFP, D42-Gal4/*+, (U) *NDUFS7 [RNAi]/UAS-cat; UAS-mitoGFP, D42-Gal4/*+ and (V) *NDUFS7 [RNAi]/UAS-sod1; UAS-mitoGFP, D42-Gal4/*+ larvae immunostained with antibodies against HRP(magenta) and GFP (mito-GFP:green) to label neurons and mitochondria. *NDUFS7*-depleted and *cat* or *sod1* rescued *NDUFS7* [RNAi] animals harbor fewer mitochondria at the terminals than control animals. (Q-V) Scale bar: 5 µm. (W) Histograms showing quantification of mitochondrial clusters in the above-indicated genotypes. At least 8 NMJs of each genotype were used for quantification. ****p*<0.0001. Error bars represent mean ± s.e.m. Statistical analysis based on one-way ANOVA followed by post-hoc Tukey's multiple-comparison test. Raw data for this figure are available in the S2 Data Excel file, tab Figure_S13.

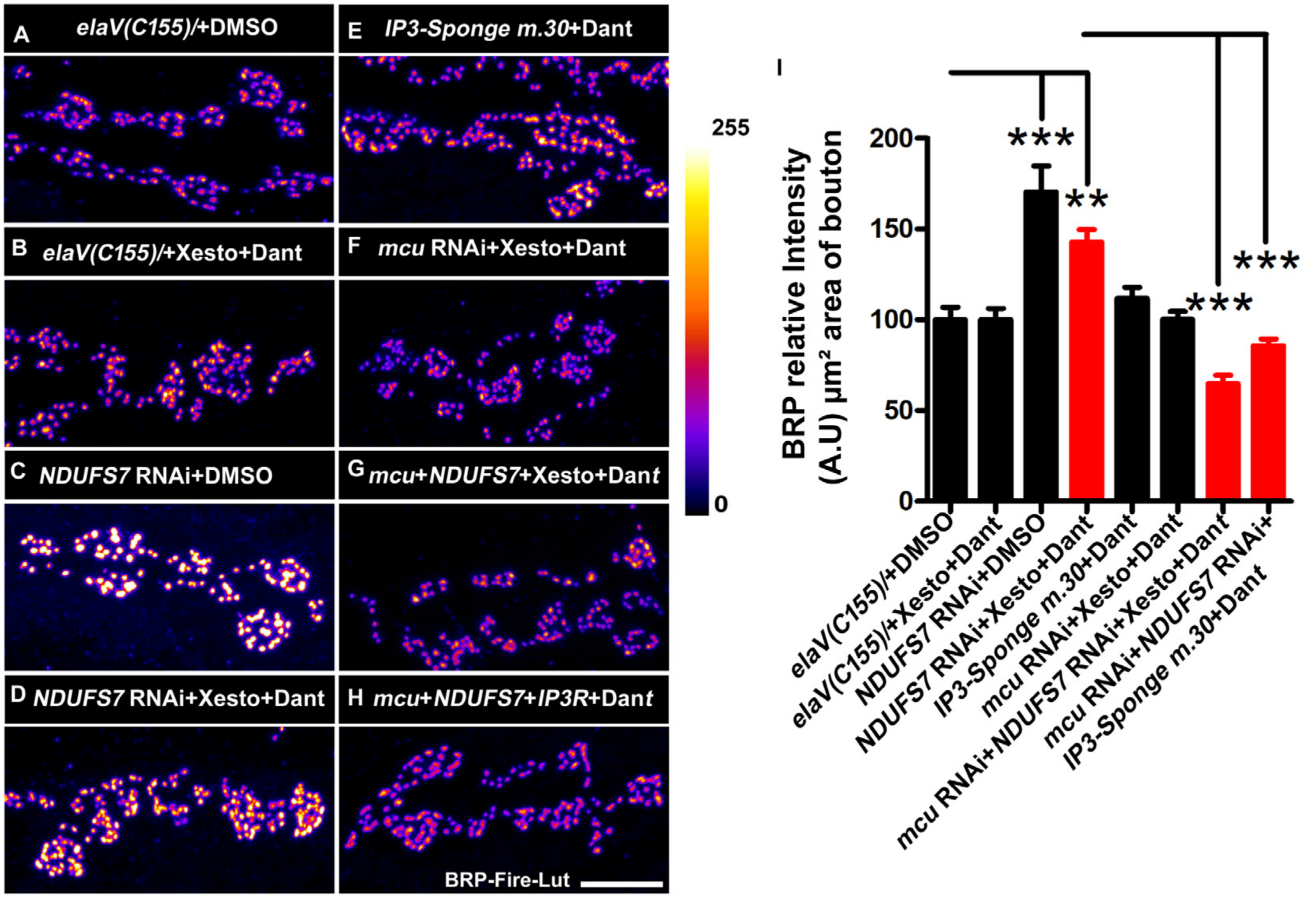


**Figure S14, related to Figure 5: Loss of MCI modulates activities of intracellular calcium channels and regulates levels of active zone materials to stabilize synaptic strength**

(A) Representative images of the A2 hemi segment of muscle 6/7 NMJs in *elaV(C155)/*+DMSO, (B) *elaV(C155)/*+Xesto+Dant, (C) *elaV(C155)/+; UAS-NDUFS7[RNAi]/*+DMSO, (D) *elaV(C155)/+; UAS-NDUFS7[RNAi]/***+**Xesto+Dant, (E) *elaV(C155)/+UAS-ip3-sponge m.30/*+, (F) *elaV(C155)/+; mcu RNAi/+,* (G) *elaV(C155)/+;mcu RNAi/UAS-NDUFS7[RNAi]*+ Xesto + Dant and (H) *elaV(C155)/+;mcu RNAi/UAS-NDUFS7[RNAi]/*+; *UAS-ip3-sponge.m30/*+Dant larvae immunostained with antibodies against the active zone scaffold Bruchpilot (BRP:fire-LuT) to label the active zones. While BRP levels are upregulated at the NMJs in *UAS-NDUFS7[RNAi]* + DMSO and *UAS-NDUFS7[RNAi]* + Xesto + Dant flies, blocking ER calcium release and import to the mitochondria reduces BRP levels below the control level. (A-H) Scale bar: 5 µm. (I) Histograms showing quantification of BRP intensity in the µm$^2$ area of bouton at muscle 6/7 in the above genotypes. At least 8 NMJs of each genotype were used for quantification. ***$p$<0.0001 **$p$=0.0002, (BRP levels: *elaV(C155)/*+DMSO *vs elaV(C155)/+; UAS-NDUFS7[RNAi]/*+Xesto+Dant, Error bars denote mean ± s.e.m. Statistical analysis based on one-way ANOVA followed by post-hoc Tukey's multiple-comparison test. Raw data for this figure are available in the S2 Data Excel file, tab Figure_S14.

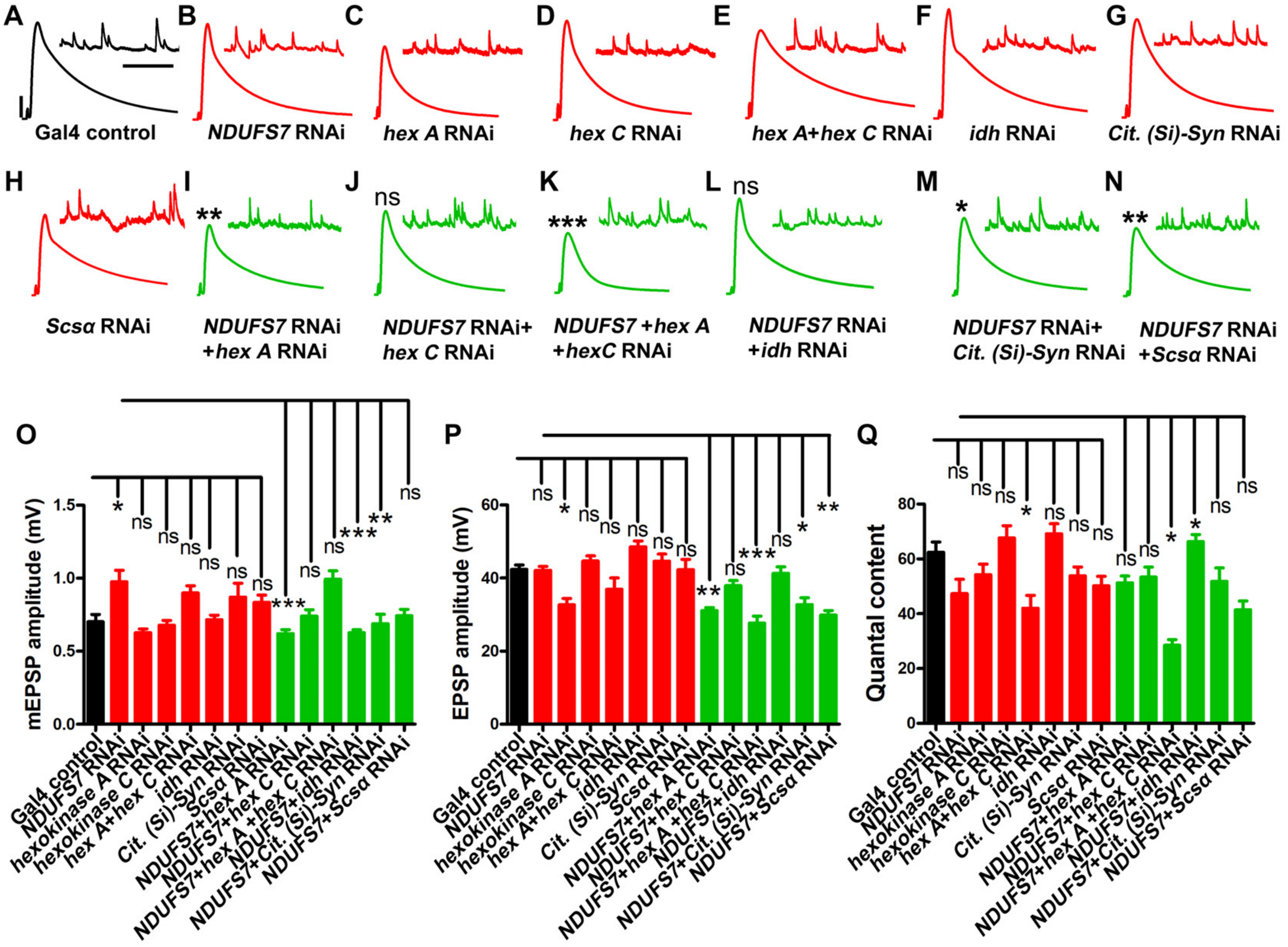


**Figure S15, related to Figure 6: Combined genetic loss in neurons of an MCI subunit and glycolysis or TCA cycle genes impairs evoked neurotransmission.**

(A-N) Example electrophysiological traces of Gal4 driver control (*elaV(C155)-Gal4/+*) or experimental genotypes tested for potential combinatorial effects of losing *NDUFS7* gene function and/or the function of glycolysis or TCA cycle genes, including: *hexokinase A (hex-A)*, *hexokinase C (hex-C)*, *Citrate (Si) Synthase I*, *Isocitrate dehydrogenase* (*Idh*), and *Succinyl-coenzyme A synthetase α subunit 1* (*Scsa1*). Scale bars for EPSPs (mEPSP) are x=50 ms (1000 ms) and y= 10 mV (1 mV). (O-Q) Data histograms and statistical analyses for these same genotypes (knockdowns all in the *elaV(C155)-Gal4/+* genetic background; see Table 17 for full genotypes), including quantal size (mEPSP), evoked excitation (EPSP) and quantal content. Error bars denote mean ± s.e.m. Statistical analysis based on one-way ANOVA followed by post-hoc Tukey's multiple-comparison test. *$p < 0.05$, **$p < 0.01$, ***$p < 0.001$. Raw data for this figure are available in the S2 Data Excel file, tab Figure_S15.

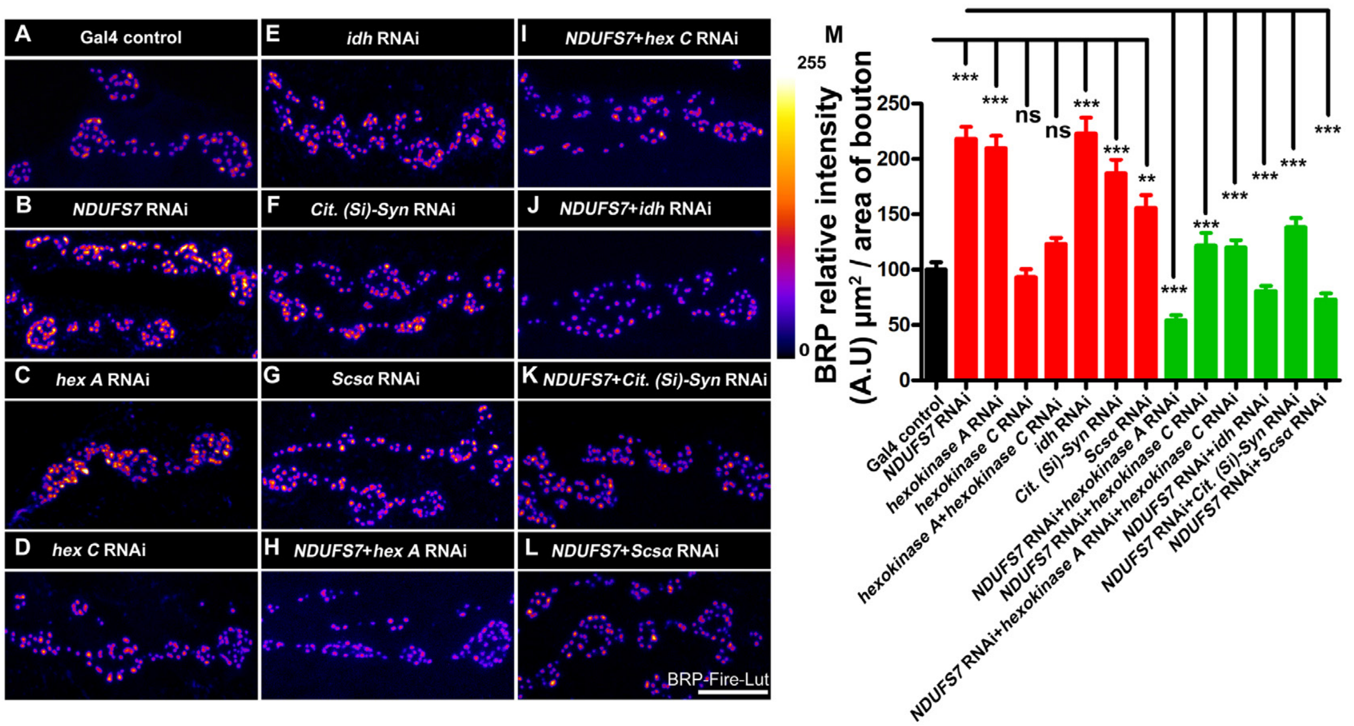


**Figure S16, related to Figure 6: Loss in neurons of glycolysis or TCA cycle genes reverses the active zone enhancements normally induced by *NDUFS7* loss.**

(A-L) Representative images of the A2 hemi segment of muscle 6/7 NMJs, examining active zone material through anti-Bruchpilot immunostaining. Analysis for Gal4 driver control (*elaV(C155)-Gal4/+*) or experimental genotypes tested for potential combinatorial effects of losing *NDUFS7* gene function and/or the function of glycolysis or TCA cycle genes, including *hexokinase A (hex-A)*, *hexokinase C (hex-C)*, *Citrate (Si) Synthase I*, *Isocitrate dehydrogenase* (*Idh*), and *Succinyl-coenzyme A synthetase α subunit 1* (*Scsa1*). Scale bar: 5 μm. Full genotypes in Table 18. (M) Histograms showing quantification of BRP intensity in the $\mu m^2$ area of bouton at muscle 6/7 in the above genotypes. At least 8 NMJs of each genotype were used for quantification. Error bars signify mean ± s.e.m. Raw data for this figure are available in the S2 Data Excel file, tab Figure_S16.

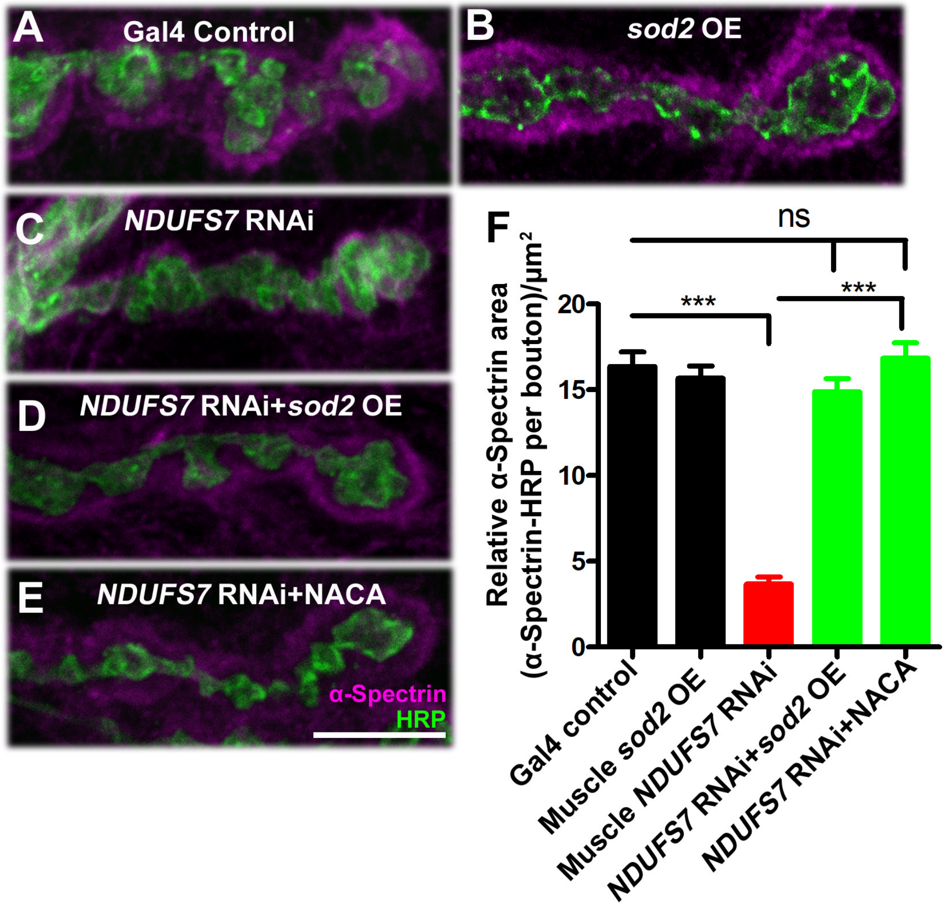


**Figure S17, related to Figure 7: *NDUFS7* subunit in muscle affects the organization of α-Spectrin scaffold at the NMJs**

(A-E) Representative confocal images of third instar larval NMJs in (A) Muscle-Gal4 control (*BG57-Gal4/+*), (B) Muscle Gal4 driven *UAS-Sod2* (*UAS-Sod2*/*+; BG-57-Gal4/+*), (C) Muscle *UAS-NDUFS7[RNAi]* (*UAS-NDUFS7[RNAi]/+; BG-57-Gal4/+*), (D) *Sod2* muscle rescue (*UAS-Sod2/UAS-NDUFS7[RNAi]; BG-57-Gal4/BG-57-Gal4*) and (E) (*UAS-NDUFS7[RNAi]/+; BG57-Gal4/+*NACA) synapses, immunostained with anti-HRP (green) and α-Spectrin (magenta) antibodies. Scale bar: 5 μm. (N) Histogram showing relative α-Spectrin area in the indicated genotypes. Compared with controls, *UAS-NDUFS7[RNAi]/+; BG-57-Gal4/+)* NMJs show a significant reduction in α-Spectrin area, which is rescued upon muscle overexpression of a *Sod2* transgene or feeding larvae with NACA. ***$p$<0.0001; ns, not significant. Error bars signify mean ± s.e.m. Statistical analysis based on one-way ANOVA with post-hoc Tukey's test for multiple comparisons. Raw data for this figure are available in the S2 Data Excel file, tab Figure_S17.

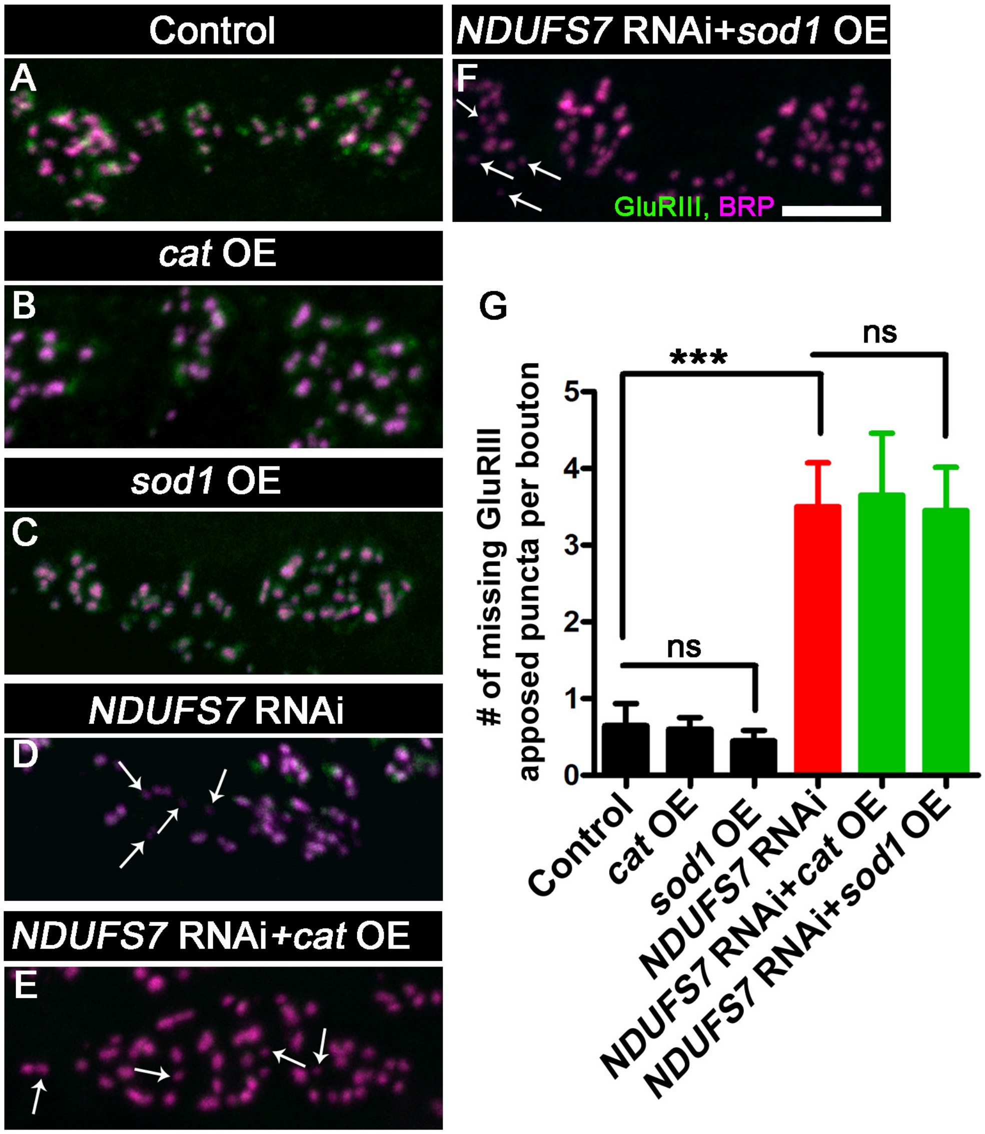


**Figure S18, related to Figure 8: Both *Catalase* and *Sod1* overexpression fail to restore defective glutamate receptor-active zone apposition in *NDUFS7*-deficient animals**

The NDUFS7 subunit in muscle affects the organization of the GluR cluster in *Drosophila*. Representative confocal images of boutons at third instar larval NMJ synapse in (A) Muscle-Gal4 control (*BG57-Gal4/+)*, (B) Muscle Gal4 driven UAS-*cat* (*UAS-cat/+; BG57-Gal4/+)*, (C) Muscle Gal4 driven UAS-*sod1* (*UAS-sod1/+; BG57-Gal4/+*), (D) Muscle *NDUFS7[RNAi]* (*NDUFS7[RNAi]/+; BG57-Gal4/+*), (E) *Cat* muscle rescue (*UAS-cat/NDUFS7 [RNAi]; BG57-Gal4/BG57-Gal4*) and (F) *Sod1* muscle rescue (*UAS-sod1/NDUFS7 [RNAi]; BG57-Gal4/BG57-Gal4*) animals immunolabeled with active zone marker BRP (magenta) and anti-GluRIII (green) antibodies. Scale bar: 2.5 µm. Note that GluRIII apposed clusters with BRP are missing in the *NDUFS7* [RNAi]-depleted, *cat* and *sod1* rescued animals (marked in arrow) compared to the control. (G) Histograms showing quantification of the number of missing BRP-GluRIII apposed puncta per bouton in the indicated genotypes. ***$p<0.0001$. Error bars represent mean ±s.e.m. Statistical analysis based on one-way ANOVA with post-hoc Tukey's test for multiple comparisons. Raw data for this figure are available in the S2 Data Excel file, tab Figure_S18.
