## Supplemental Tables for "Mitochondrial Complex I and ROS control neuromuscular function through opposing pre- and postsynaptic mechanisms"

This is Supplemental Document S2 for Mallik *et al*. It contains the following:

- S1-S21 Tables: Summary data for graphs in all Figures and Supplemental Figures.
- S22 Table: Detailed information about reagents and materials for items described in the STAR METHODS section.

**S1 Table, related to Figure 2:** Mitochondrial branch length (µm) and cluster number (µm^2^ area of the axon).

| **Genotype** | **Mitochondria branch**  **length (µm) in VNC** | **# clusters (per µm^2^ area**  **of a nerve at A5 hemisegment)** |
| --- | --- | --- |
| *UAS-mitoGFP, D42-Gal4/+* | 1.74 ± 0.07, n = 50 | 0.79 ± 0.03, n = 30 |
| *UAS-drp1[RNAi]/+; UAS-mitoGFP, D42-Gal4/+* | 1.69 ± 0.08, n = 50 | 0.80 ± 0.02, n = 30 |
| *UAS-NDUFS7[RNAi]/+; UAS-mitoGFP, D42-Gal4/+* | 0.96 ± 0.04, n = 50 | 0.27 ± 0.01, n = 30 |
| *UAS-mitoGFP, D42-Gal4/UAS-marf[RNAi]* | 0.89 ± 0.03, n = 50 | 0.20 ± 0.02, n = 30 |
| *UAS-NDUFS7[RNAi]/+; UAS-mitoGFP, D42-Gal4/UAS-marf[RNAi]* | 0.83 ± 0.03, n = 50 | 0.20 ± 0.01, n = 30 |
| *UAS-NDUFS7 [RNAi]/UAS-drp1[RNAi]; UAS-mitoGFP, D42-Gal4/+* | 0.91 ± 0.03, n = 50 | 0.32 ± 0.02, n = 30 |

*D42-Gal4* is a motor neuron driver. Values represent mean ± s.e.m.

**S2 Table, related to Figure 2 and Figure S4:** mitochondrial branch length (µm) and cluster number (per µm^2^ area of the axon).

| **Genotype** | **Mito branch length (µm) in VNC** | **# clusters (per µm^2^ area of nerve A5 hemi segment)** |
| --- | --- | --- |
| *D42-Gal4, UAS-mitoGFP/+* | 1.88 ± 0.07, n = 50 | 0.94 ± 0.05, n = 30 |
| *UAS-Sod2/+; D42-Gal4, UAS-mitoGFP/+* | 1.75 ± 0.06, n = 50 | 1.10 ± 0.06, n = 35 |
| *UAS-NDUFS7 [RNAi]/+;*  *D42-Gal4, UAS-mitoGFP/+* | 0.99 ± 0.03, n = 50 | 0.21 ± 0.01, n = 30 |
| *UAS-NDUFS7 [RNAi]/+;*  *D42-Gal4,UAS-mitoGFP/+* with NACA | 1.63 ± 0.06, n = 50 | 0.59 ± 0.04, n = 30 |
| *UAS-NDUFS7 [RNAi]/UAS-Sod2;*  *D42-Gal4,UAS-mitoGFP/+* | 1.94 ± 0.08, n = 50 | 0.57 ± 0.03, n = 30 |
| *UAS-ND-30*[RNAi]/+;  *D42-Gal4, UAS-mitoGFP/+* | 0.89 ± 0.03, n = 50 | 0.32 ± 0.01, n = 30 |
| *UAS-Sod2*/*UAS-ND-30[RNAi]*;  *D42-Gal4, UAS-mitoGFP/+* | 1.78 ± 0.06, n = 50 | 0.83 ± 0.04, n = 30 |

*D42-Gal4* is a motor neuron driver. Values represent mean ± s.e.m.

**S3 Table, related to Figure 2 and S5:** mitochondrial branch length (µm) and cluster number (per µm^2^ area of the axon).

| **Genotype** | **Mitochondrial branch**  **length (µm) in VNC** | **# clusters (per µm^2^ area of axon at A5 hemi segment)** |
| --- | --- | --- |
| *UAS-mitoGFP, D42-Gal4/+* | 1.21 ± 0.05, n=40 | 1.43 ± 0.06, n=30 |
| *UAS-Cat/+;*  *UAS-mitoGFP, D42-Gal4/+* | 1.22 ± 0.05, n=40 | 1.50 ± 0.09, n=30 |
| *UAS-Sod1/+;*  *UAS-mitoGFP, D42-Gal4/+* | 1.25 ± 0.03, n=40 | 1.51 ± 0.11, n=30 |
| *UAS-NDUFS7[RNAi]/+;*  *UAS-mitoGFP, D42-Gal4/+* | 0.72 ± 0.03, n=40 | 0.68 ± 0.04, n=30 |
| *UAS-NDUFS7[RNAi]/UAS-Cat;*  *UAS-mitoGFP, D42-Gal4/+* | 0.65 ± 0.02, n=40 | 0.54 ± 0.05, n=30 |
| *UAS-NDUFS7[RNAi]/UAS-Sod1;*  *UAS-mitoGFP, D42-Gal4/+* | 0.69 ± 0.02, n=40 | 0.66 ± 0.04, n=30 |

*D42-Gal4* is a motor neuron driver. Values represent mean ± s.e.m.

**S4 Table, related to Figure 2:** mitochondrial cluster number (per µm^2^ area of bouton).

| **Genotype** | **# clusters (per µm^2^ area of bouton)** |
| --- | --- |
| *UAS-mitoGFP, D42-Gal4/+* | 4.67 ± 0.26, n = 40 |
| *UAS-Sod2/+; UAS-mitoGFP, D42-Gal4/+* | 5.2 ± 0.20, n = 40 |
| *UAS-NDUFS7[RNAi]/+;*  *UAS-mitoGFP, D42-Gal4/+* | 1.40 ± 0.15, n = 40 |
| *UAS-NDUFS7[RNAi]/UAS-Sod2;*  *UAS-mitoGFP, D42-Gal4/+* | 2.07 ± 0.15, n = 40 |
| *UAS-drp1[RNAi]/+; UAS-mitoGFP, D42-Gal4/+* | 4.65 ± 0.24, n = 40 |
| *UAS-drp1[RNAi]/UAS-Sod2;*  *UAS-mitoGFP, D42-Gal4/+* | 5.47 ± 0.22, n = 40 |
| *UAS-drp1[RNAi]/UAS-NDUFS7[RNAi];*  *UAS-mitoGFP, D42-Gal4/+* | 1.15 ± 0.19, n = 40 |
| *UAS-mitoGFP, D42-Gal4/UAS-marf[RNAi]* | 1.50 ± 0.22, n = 40 |
| *UAS-Sod2/+;*  *UAS-mitoGFP, D42-Gal4/UAS-marf[RNAi]* | 2.40 ± 0.19, n = 40 |
| *UAS-NDUFS7[RNAi]/+;*  *UAS-mitoGFP, D42-Gal4/UAS-marf[RNAi]* | 0.62 ± 0.13, n = 40 |
| *UAS-NDUFS7[RNAi]/UAS-Sod2;*  *UAS-mitoGFP, D42-Gal4/UAS-marf[RNAi]* | 0.92 ± 0.14, n = 40 |

*D42-Gal4* is a motor neuron driver. Values represent mean ± s.e.m.

**S5 Table, related to Figures S2 and S6:** MitoSOX intensity as an indicator of mitochondrial ROS.

| **Genotype** | **MitoSOX Intensity in AU (µm^2^ area of indicated tissue)** |
| --- | --- |
| **Ventral nerve cord (VNC)** (MCI genetic manipulation) | |
| *D42-Gal4, UAS-mitoGFP/+* | 100.0 ± 4.47, n = 30 |
| *UAS-NDUFS7[RNAi]/+;*  *D42-Gal4, UAS-mitoGFP/+* | 352.1 ± 8.44, n = 30 |
| *UAS-NDUFS7[RNAi]/+;*  *D42-Gal4, UAS-mitoGFP/+* with NACA | 83.50 ± 5.18, n = 30 |
| *UAS-NDUFS7[RNAi]/UAS-Sod2;*  *UAS-mitoGFP, D42-Gal4/+* | 94.21 ± 6.98, n = 30 |
| **Muscle** | |
| *UAS-mitoGFP/+; BG57-Gal4/+* | 100.0 ± 8.12, n = 30 |
| *UAS-NDUFS7[RNAi]/UAS-mitoGFP;*  *BG57-Gal4/+* | 267.6 ± 14.62, n = 30 |
| *UAS-NDUFS7[RNAi]/UAS-mitoGFP;*  *BG57-Gal4/+* with NACA | 93.46 ± 8.10, n = 30 |
| *UAS-NDUFS7[RNAi]/UAS-Sod2; BG57-Gal4/+* | 92.34 ± 5.31, n = 30 |
| **Ventral nerve cord (VNC)** (Mitofusin or pharmacology against MCI) | |
| *D42-Gal4, UAS-mitoGFP/+* | 100.0 ± 13.84, n = 30 |
| *D42-Gal4, UAS-mitoGFP/UAS-marf[RNAi]* | 1067 ± 91.59, n = 37 |
| *D42-Gal4, UAS-mitoGFP/+* with 25µM ROT | 405.3 ± 40.72, n = 22 |
| *UAS-NDUFS7[RNAi]/+;*  *D42-Gal4, UAS-mitoGFP/+* | 645.5 ± 37.13, n = 47 |
| **Axon** | |
| *D42-Gal4, UAS- mitoGFP/+* | 100.0 ± 12.28, n = 30 |
| *D42-Gal4,UAS-mitoGFP/UAS-marf[RNAi]* | 325.9 ± 23.02, n = 48 |
| *D42-Gal4,UAS- mitoGFP/+* with 25µM ROT | 490.0 ± 52.21, n = 40 |
| *UAS-NDUFS7[RNAi]/+;*  *D42-Gal4, UAS-mitoGFP/+* | 919.6 ± 66.61, n = 65 |
| **Bouton** | |
| *D42-Gal4, UAS-mitoGFP/+* | 100.0 ± 10.49, n = 50 |
| *D42-Gal4,UAS-mitoGFP/UAS-marf[RNAi]* | 254.7 ± 46.29, n = 53 |
| *D42-Gal4,UAS- mitoGFP/+* with 25µM ROT | 271.5 ± 24.21, n = 56 |
| *NDUFS7 [RNAi]/+; D42-Gal4, UAS-mitoGFP/+* | 223.2 ± 19.16, n = 59 |

NACA – N-Acetyl Cysteine Amide; ROT – Rotenone. *D42-Gal4* is a motor neuron driver. *BG57-Gal4* is a muscle driver. Values represent mean ± s.e.m.

**S6 Table, related to Figure S3:** MitoSOX intensity as an indicator of mitochondrial ROS for genetic combinations, as analyzed in the ventral nerve cord, muscle, axon, or boutons.

| **Genotype** | **MitoSOX Intensity in AU**  **(µm^2^ area of VNC, axon,**  **boutons and muscle** |
| --- | --- |
| **Ventral nerve cord (VNC)** | |
| *UAS-mitoGFP, D42-Gal4/+* | 100.0 ± 9.98, n=20 |
| *UAS-mitoGFP, D42-Gal4/UAS-Sod2[RNAi]* | 175.6 ± 15.52, n=22 |
| **Muscle** | |
| *UAS-mitoGFP/+; BG57-Gal4/+* | 100.0 ± 6.01, n=41 |
| *UAS-mitoGFP/+; BG57-Gal4/UAS-Sod2[RNAi]* | 145.5 ± 9.42, n=40 |
| **Axon** | |
| *UAS-mitoGFP, D42-Gal4/*+ | 100.0 ± 6.99, n=28 |
| *UAS-mitoGFP, D42-Gal4/UAS-Sod2[RNAi]* | 143.4 ± 17.07, n=27 |
| **Bouton** | |
| *UAS-mitoGFP, D42-Gal4/*+ | 100.0 ± 10.56, n=25 |
| *UAS-mitoGFP, D42-Gal4/UAS-Sod2[RNAi]* | 266.8 ± 14.67, n=25 |

*D42-Gal4* is a motor neuron driver. *BG57-Gal4* is a muscle driver. Values represent mean ± s.e.m.

**S7 Table, related to Figures 3, 8, S7, S17:** Cell and synapse-level phenotypes from tissue-specific losses of MCI.

| **Genotype** | **% of A5 axons showing CSP accumulation** | **% Futsch positive loops** | **Genotype** | **α-Spectrin**  **levels (AU)** | **# missing apposed BRP- puncta (per bouton)** |
| --- | --- | --- | --- | --- | --- |
| *D42-Gal4/+* | 12.50 ± 12.50,  n = 8 | 36.47 ± 2.88,  n = 8 | *BG57-Gal4/+* | 16.35 ± 0.84,  n = 30 | GluRIII: 0.75 ± 0.23, n = 20  IIA: 1.09 ± 0.23,  n = 21 |
| *UAS-Sod2/+;*  *D42-Gal4/+* | 16.67 ± 16.67,  n = 6 | 50.72 ± 3.95,  n = 8 | *NDUFS7 [RNAi]/+; BG57-Gal4/+* | 15.67 ± 0.72,  n = 30 | GluRIII: 3.50 ± 0.62, n = 20  IIA: 4.19 ± 0.62,  n = 21 |
| *UAS-NDUFS7 [RNAi]/+; D42-Gal4/+* | 87.50 ± 12.50,  n = 8 | 21.54 ± 1.97,  n = 8 | *UAS-Sod2*/*+;*  *BG57-Gal4/+* | 3.67 ± 0.41,  n = 30 | GluRIII: 0.30 ± 0.12, n = 20  IIA: 0.66 ± 0.21,  n = 21 |
| *UAS-NDUFS7 [RNAi]/+; D42-Gal4/+*  with NACA | 16.67 ± 16.67,  n = 6 | 40.52 ± 4.36,  n = 8 | *NDUFS7 RNAi/+;BG57-Gal4/+*  with NACA | 14.89 ± 0.77,  n = 30 | GluRIII: 0.50 ± 0.22, n = 20  IIA: 0.52 ± 0.13,  n = 21 |
| *NDUFS7 [RNAi]/*  *UAS-Sod2;*  *D42-Gal4/+* | 16.67 ± 11.24,  n = 12 | 38.97 ± 3.34,  n = 8 | *UAS-Sod2/NDUFS7 [RNAi]; BG57-Gal4/BG57-Gal4* | 16.84 ± 0.91,  n = 30 | GluRIII: 0.40 ± 0.13, n = 20  IIA: 0.61 ± 0.20,  n = 21 |

For presynaptic genotypes (left): % of A5 axons showing accumulation of CSP, % of Futsch positive loops. For postsynaptic genotypes: α-Spectrin levels and number of missing BRP puncta apposed to glutamate receptor cluster (GluRIII is also known as GluRIIC; IIA denotes GluRIIA). *D42-Gal4* is a motor neuron driver. *BG57-Gal4* is a muscle driver. Values represent mean ± s.e.m.

**S8 Table, related to Figures S8, S9, and S18:** Cell and synapse-level phenotypes from tissue-specific loss of MCI.

| **Genotype** | **% of A5 axon**  **showing CSP**  **accumulation** | **% Futsch**  **positive loops** | **Genotype** | **# missing**  **BRP-GluRIII apposed**  **puncta per bouton** |
| --- | --- | --- | --- | --- |
| *UAS-mitoGFP,*  *D42-Gal4/+* | Normal:  88.89 ± 12.50, n=9  Accumulation:  11.11 ± 11.11, n=9 | 68.75 ± 3.13,  n=14 | *BG57-Gal4/+* | GluRIII: 0.65 ± 0.28, n=20 |
| *UAS-Cat/+;*  *UAS-mitoGFP,D42-Gal4* | Normal:  83.89 ± 11.11, n=9  Accumulation:  11.11 ± 11.11, n=9 | 70.93 ± 3.76,  n=10 | *UAS-Cat/+;*  *BG57-Gal4/+* | GluRIII: 0.60 ± 0.15, n=20 |
| *UAS-Sod1/+;*  *UAS-mitoGFP,D42-Gal4* | Normal:  92.31 ± 7.69, n=13  Accumulation:  7.69 ± 7.69, n=13 | 71.74 ± 2.51,  n=9 | *UAS-Sod1/+;*  *BG57-Gal4/+* | GluRIII: 0.45 ± 0.13, n=20 |
| *NDUFS7 [RNAi]/+;*  *UAS-mitoGFP,D42-Gal4/+* | Normal:  22.22 ± 14.70, n=9  Accumulation:  77.78 ± 14.70, n=9 | 33.03 ± 2.48,  n=10 | *NDUFS7 [RNAi]/+;*  *BG57-Gal4/+* | GluRIII: 3.50 ± 0.57, n=20 |
| *NDUFS7 [RNAi]/UAS-Cat;*  *UAS-mitoGFP, D42-Gal4/+* | Normal:  20.00 ± 13.33, n=10  Accumulation:  80.00 ± 13.33, n=10 | 40.32 ± 2.85,  n=10 | *NDUFS7 [RNAi]/UAS-Cat;*  *BG57-Gal4/BG57-Gal4* | GluRIII: 3.65 ± 0.80, n=20 |
| *NDUFS7 [RNAi]/UAS-Sod1; UAS-mitoGFP,D42-Gal4/+* | Normal:  22.22 ± 14.70, n=9  Accumulation:  77.78 ± 14.70, n=9 | 39.88 ± 1.84,  n=10 | *NDUFS7 RNAi/UAS-Sod1;*  *BG57-Gal4/BG57-Gal4* | GluRIII: 3.45 ± 0.56, n=20 |

For presynaptic genotypes (left): % of A5 axons showing accumulation of CSP, % of Futsch positive loops. For postsynaptic genotypes: Number of missing BRP puncta apposed to glutamate receptor cluster (GluRIII is also known as GluRIIC. *D42-Gal4* is a motor neuron driver. *BG57-Gal4* is a muscle driver. Values represent mean ± s.e.m.

**S9 Table, related to Figure S10:** Electrophysiological parameters when NDUFS7 and Mitofusin (*marf*) functions are impaired – mEPSPs, EPSPs, mEPSP frequencies, and QC.

| **Genotype** | **mEPSP amp (mV)** | **mEPSP freq (Hz)** | **EPSP amplitude (mV)** | **Quantal content (QC)** |
| --- | --- | --- | --- | --- |
| *UAS-mitoGFP; D42-Gal4/+* | 1.06 ± 0.05,  n = 10 | 4.12 ± 0.72,  n = 10 | 49.27 ± 0.99,  n = 10 | 47.30 ± 2.49,  n = 10 |
| *UAS-Sod2/+;*  *UAS-mitoGFP, D42-Gal4/+* | 0.86 ± 0.08,  n = 9 | 2.77 ± 0.51,  n = 9 | 43.53 ± 2.55,  n = 10 | 53.16 ± 4.43,  n = 9 |
| *NDUFS7 [RNAi]/+; UAS-mitoGFP, D42-Gal4/+* | 0.79 ± 0.02,  n = 13 | 3.30 ± 0.41,  n = 13 | 45.49 ± 1.18,  n = 13 | 57.79 ± 2.39,  n = 13 |
| *UAS-mitoGFP, D42-Gal4/UAS-marf[RNAi]* | 0.63 ± 0.05,  n = 11 | 4.99 ± 0.76,  n = 11 | 39.68 ± 1.39,  n = 12 | 65.35 ± 5.30,  n = 11 |
| *UAS-Sod2/+;*  *UAS-mitoGFP, D42-Gal4/UAS-marf[RNAi]* | 0.83 ± 0.06,  n = 8 | 5.01 ± 0.74,  n = 8 | 37.45 ± 2.08,  n = 9 | 47.52 ± 4.55,  n = 8 |
| *UAS- NDUFS7 [RNAi]/+;*  *UAS- mitoGFP, D42-Gal4/UAS-marf[RNAi]* | 0.90 ± 0.07,  n = 10 | 6.23 ± 0.66,  n = 10 | 35.44 ± 1.80,  n = 12 | 40.82 ± 3.70,  n = 10 |
| *UAS-Sod2/UAS-NDUFS7 [RNAi];*  *UAS-mitoGFP, D42-Gal4/UAS-marf[RNAi]* | 0.66 ± 0.04,  n = 9 | 5.26 ± 0.70,  n = 9 | 35.50 ± 1.9,  n = 10 | 56.00 ± 4.51,  n = 9 |

*D42-Gal4* is a motor neuron driver. Values represent mean ± s.e.m.

**S10 Table, related to Figure S11:** Electrophysiological properties for motor neuron MCI loss (or rescue) conditions.

| **Recording Condition** | **Genotype** | **Electrophysiological Data** |
| --- | --- | --- |
| 0.15 mM Ca^2+^ | *D42-Gal4/+* | EPSP: 17.61 ± 4.16 mV, n = 10 |
|  | *UAS-NDUFS7 [RNAi]/+;*  *D42-Gal4/+* | EPSP: 9.52 ± 2.01 mV, n = 11 |
| Failure analysis:  0.1 mM Ca^2+^  Data: % failures  n = number of muscles | *D42-Gal4/+* | 18.08% ± 7.59, n = 13 |
|  | *UAS-Sod2/+;D42-Gal4/+* | 27.00% ± 8.49, n = 8 |
|  | *NDUFS7 [RNAi]/+; D42-Gal4/+* | 1.87% ± 0.44, n = 8 |
|  | *NDUFS7 [RNAi]/+; D42-Gal4/+*NACA | 60.22% ± 15.16, n = 9 |
|  | *NDUFS7 [RNAi]/UAS-Sod2; D42-Gal4/+* | 20.13% ± 8.18, n = 15 |
| Paired pulse ratios  (EPSP2/EPSP1)  0.4 mM Ca^2+^ | *D42-Gal4/+* | 0.88 ± 0.06, n = 8 |
|  | *NDUFS7 [RNAi]/+; D42-Gal4/+* | 0.82 ± 0.01, n = 10 |
| Paired pulse ratios  (EPSP2/EPSP1)  1.5 mM Ca^2+^ | *D42-Gal4/+* | 0.88 ± 0.02, n = 12 |
|  | *NDUFS7 [RNAi]/+; D42-Gal4/+* | 0.84 ± 0.01, n = 11 |

*D42-Gal4* is a motor neuron driver. Values represent mean ± s.e.m.

**S11 Table, related to Figure 4:** BRP intensity, BRP density, electrophysiological parameters, and the number of mitochondrial clusters for genetic combinations shown in Figure 4.

| **Genotype** | **BRP Intensity in AU (µm^2^ area of bouton)** | **BRP density (per µm^2^ area**  **of bouton)** |
| --- | --- | --- |
| *UAS-mitoGFP, D42-Gal4/+* | 100.0 ± 7.83, n = 40 | 2.28 ± 0.06, n = 29 |
| *UAS-Sod2/+;*  *UAS-mitoGFP, D42-Gal4/+* | 84.29 ± 5.50, n = 38 | 2.29 ± 0.09, n = 30 |
| *UAS-NDUFS7 [RNAi]/+;*  *UAS-mitoGFP, D42-Gal4/+* | 195.2 ± 22.90, n = 42 | 1.78 ± 0.09, n = 30 |
| *NDUFS7 [RNAi]/+; UAS-mitoGFP, D42-Gal4/+*  with NACA | 110.8 ± 13.72, n = 41 | 2.22 ± 0.06, n = 30 |
| *NDUFS7 [RNAi]/UAS-Sod2;*  *UAS-mitoGFP, D42-Gal4/+* | 98.29 ± 7.96, n = 40 | 2.24 ± 0.10, n = 30 |
| **Genotype** | **mEPSP amp (mV), mEPSP freq (Hz) EPSP (mV), and QC** | **# mito clusters (per µm^2^  area of bouton)** |
| *UAS-mitoGFP, D42-Gal4/+* | mEPSP amp: 0.98 ± 0.04, n = 8,  mEPSP freq: 2.51 ± 0.12, n = 8  EPSP amp: 43.29 ± 1.75, n = 8  QC: 44.21 ± 0.72, n = 8 | 12.42 ± 1.03, n = 38 |
| *UAS-Sod2/+;*  *UAS-mitoGFP, D42-Gal4/+* | mEPSP amp: 0.58 ± 0.023, n = 8,  mEPSP freq: 2.82 ± 0.56, n = 8  EPSP amp: 40.78 ± 0.50, n = 8  QC: 70.66 ± 3.02, n = 8 | 12.23 ± 0.78, n = 39 |
| *UAS-NDUFS7 RNAi]/+;*  *UAS-mitoGFP, D42-Gal4/+* | mEPSP amp: 0.75 ± 0.03, n = 7,  mEPSP freq: 4.92 ± 0.81, n = 7  EPSP amp: 43.67 ± 1.55, n = 7  QC: 58.31 ± 3.92, n = 7 | 4.84 ± 0.44, n = 38 |
| *UAS-NDUFS7[RNAi]/UAS-Sod2;*  *UAS-mitoGFP, D42-Gal4/+* | mEPSP amp: 0.65 ± 0.02, n = 14,  mEPSP freq: 3.18 ± 0.32, n = 14  EPSP amp: 37.62 ± 1.28, n = 14  QC: 58.06 ± 2.76, n = 14 | 5.48 ± 0.48, n = 41 |

*D42-Gal4* is a motor neuron driver. Values represent mean ± s.e.m.

**S12 Table, related to Figure S12:** Analysis of BRP intensity for various rotenone applications.

| **Genotype** | **BRP Intensity in AU**  **(µm^2^ area of a bouton)** |
| --- | --- |
| *w^1118^ +* DMSO, nerve severed, 1 hour | 100.0 ± 5.73, n = 75 |
| *w^1118^ +* DMSO, nerve intact, 1 hour | 76.18 ± 2.79, n = 74 |
| *w^1118^ +* 500 µM ROT, nerve severed, 1 hour | 100.0 ± 4.98, n = 68 |
| *w^1118^ +* 500 µM ROT, nerve intact, 1 hour | 105.3 ± 4.65, n = 77 |
| *w^1118^ +* DMSO, nerve severed, 2 hours | 100.0 ± 5.86, n = 80 |
| *w^1118^ +* DMSO, nerve intact, 2 hours | 97.54 ± 5.52, n = 80 |
| *w^1118^ +* 500 µM ROT, nerve severed, 2 hours | 100.0 ± 5.09, n = 75 |
| *w^1118^ +* 500 µM ROT, nerve intact, 2 hours | 113.1 ± 6.34, n = 81 |
| *w^1118^ +* DMSO, nerve severed, 4 hours | 100.0 ± 4.79, n = 78 |
| *w^1118^ +* DMSO, nerve intact, 4 hours | 105.9 ± 4.34, n = 70 |
| *w^1118^ +* 500 µM ROT, nerve severed, 4 hours | 100.0 ± 5.31, n = 60 |
| *w^1118^ +* 500 µM ROT, nerve intact, 4 hours | 95.41 ± 4.58, n = 78 |
| *w^1118^ +* DMSO, nerve severed, 6 hours | 100.0 ± 4.08, n = 71 |
| *w^1118^ +* DMSO, nerve intact, 6 hours | 92.90 ± 4.14, n = 70 |
| *w^1118^ +* 500 µM ROT, nerve severed, 6 hours | 100.0 ± 3.90, n = 90 |
| *w^1118^ +* 500 µM ROT, nerve intact, 6 hours | 134.8 ± 4.53, n = 80 |
| *w^1118^ +* DMSO+8 hours feeding, intact larvae | 100.0 ± 5.40, n = 80 |
| *w^1118^ +* 25µM ROT+8 hours feeding, intact larvae | 94.26 ± 3.84, n = 81 |
| *w^1118^ +* DMSO+Embryo to 3^rd^ instar | 100.0 ± 3.88, n = 74 |
| *w^1118^ +* 25µM ROT+Embryo to 3^rd^ instar | 121.9 ± 5.86, n = 64 |

ROT denotes rotenone; DMSO denotes dimethyl sulfoxide (carrier control). Values represent mean ± s.e.m.

**S13 Table, related to Figures 4, S13:** BRP intensity, electrophysiological parameters, and the number of mitochondrial clusters for genetic combinations shown.

| **Genotype** | **BRP Intensity in AU**  **(µm^2^ area of a bouton)** | **# mito clusters (per µm^2^ area of a bouton)** |
| --- | --- | --- |
| *UAS-mitoGFP,*  *D42-Gal4/+* | 100.0 ± 1.91,n=26 | 6.53 ± 0.40,  n=30 |
| *UAS-Cat/+;*  *UAS-mitoGFP,D42-Gal4* | 87.80 ± 2.07, n=21 | 8.20 ± 0.45,  n=30 |
| *UAS-Sod1/+;*  *UAS-mitoGFP,D42-Gal4* | 81.85 ± 3.15, n=22 | 7.90 ± 0.68,  n=30 |
| *NDUFS7 [RNAi]/+;*  *UAS-mitoGFP, D42-Gal4/+* | 148.9 ± 9.12, n=24 | 2.76 ± 0.30,  n=30 |
| *NDUFS7 [RNAi]/UAS-Cat;*  *UAS-mitoGFP, D42-Gal4/+* | 128.0 ± 6.42, n=25 | 3.76 ± 0.38,  n=30 |
| *NDUFS7 [RNAi]/UAS-Sod1;*  *UAS-mitoGFP, D42-Gal4/+* | 152.9 ± 6.62, n=26 | 3.53 ± 0.33,  n=30 |
| **Genotype** | **mEPSP, EPSP (mV)** | **Quantal content** |
| *UAS-mitoGFP,*  *D42-Gal4/+* | mEPSP amplitude: 0.76 ± 0.05, n=7,  EPSP amplitude: 39.60 ± 1.07, n=7,  mEPSP frequency: 2.74 ± 0.40 Hz, n=7 | 53.61 ± 4.48,  n=7 |
| *UAS-Cat/+;*  *UAS-mitoGFP,*  *D42-Gal4/+* | mEPSP amplitude: 0.59 ± 0.01, n=9,  EPSP amplitude: 44.15 ± 1.15, n=9,  mEPSP frequency: 2.76 ± 0.62 Hz, n=9 | 74.38 ± 2.65,  n=9 |
| *UAS-Sod1/+;*  *UAS-mitoGFP,*  *D42-Gal4/+* | mEPSP amplitude: 0.62 ± 0.02, n=11,  EPSP amplitude: 43.27 ± 1.51, n=11,  mEPSP frequency: 1.60 ± 0.26 Hz, n=11 | 70.34 ± 2.87,  n=11 |
| *NDUFS7 [RNAi]/+;*  *UAS-mitoGFP, D42-Gal4/+* | mEPSP amplitude: 0.60 ± 0.028, n=7,  EPSP amplitude: 42.18 ± 0.61, n=7,  mEPSP frequency: 6.56 ± 0.87 Hz, n=7 | 70.43 ± 3.32, n=7 |
| *NDUFS7 [RNAi]/UAS-Cat;*  *UAS-mitoGFP, D42-Gal4/+* | mEPSP amplitude: 0.60 ± 0.023, n=12,  EPSP amplitude: 42.96 ± 1.16, n=12,  mEPSP frequency: 4.07 ± 0.67 Hz, n=12 | 72.33 ± 2.71, n=12 |
| *NDUFS7 [RNAi]/UAS-Sod1;*  *UAS-mitoGFP,D42-Gal4/+* | mEPSP amplitude: 0.54 ± 0.03, n=9,  EPSP amplitude: 38.11 ± 0.69, n=9,  mEPSP frequency: 2.65 ± 0.57 Hz, n=9 | 71.75 ± 4.27, n=9 |

*D42-Gal4* is a motor neuron driver. Values represent mean ± s.e.m.

**S14 Table, related to Figure 5:** Electrophysiological properties for various genotypes: mEPSP amplitude, EPSP amplitude, mEPSP frequency, and QC.

| **Genotype** | **mEPSP amplitude (mV)** | **EPSP**  **amplitude (mV)** | **mEPSP frequency (Hz)** | **Quantal Content (QC)** |
| --- | --- | --- | --- | --- |
| *elaV-Gal4(C155/+)* | 0.63 ± 0.01,  n = 7 | 38.22 ± 1.50,  n = 7 | 1.90 ± 0.38,  n = 7 | 60.65 ± 2.62,  n = 7 |
| *elaV(C155)-Gal4/+* with XestC (20µM) & Dantrolene (10µM) | 0.72 ± 0.05,  n = 9 | 38.21 ± 1.29,  n = 9 | 2.81 ± 0.58,  n = 9 | 54.57 ± 3.44,  n = 9 |
| *elaV(C155)-Gal4/+;*  *UAS-NDUFS7 [RNAi]/+* | 0.79 ± 0.03,  n = 10 | 39.96 ± 1.44,  n = 10 | 3.83 ± 0.40,  n = 10 | 51.45 ± 3.28,  n = 10 |
| *elaV(C155)-Gal4/+; UAS-NDUFS7 [RNAi]/+*  with XestC (20µM) & Dantrolene  (10µM) | 0.60 ± 0.04,  n = 8 | 35.97 ± 1.05,  n = 8 | 3.63 ± 0.58,  n = 8 | 61.19 ± 5.29,  n = 8 |
| *elaV(C155)-Gal4/+; UAS-CalX[RNAi]/+* | 0.80 ± 0.03,  n = 8 | 35.45 ± 1.26,  n = 8 | 1.61 ± 0.18,  n = 8 | 44.42 ± 2.59,  n = 8 |
| *elaV(C155)-Gal4/+; UAS-NDUFS7 [RNAi]/+;*  *UAS-CalX[RNAi]/+* | 0.75 ± 0.03,  n = 10 | 34.04 ± 1.72,  n = 10 | 2.68 ± 0.40,  n = 10 | 46.10 ± 3.10,  n = 10 |
| *elaV(C155)-Gal4/+; UAS-IP_3_-Sponge.m30/+*  + DMSO | 0.64 ± 0.04,  n = 8 | 33.06 ± 2.84,  n = 8 | 2.61 ± 0.51,  n = 8 | 53.31 ± 6.23,  n = 8 |
| *elaV(C155)-Gal4/+; UAS-IP_3_-Sponge.m30/+*  with XestC (20µM) & Dantrolene (10µM) | 0.79 ± 0.03,  n = 8 | 38.37 ± 0.96,  n = 8 | 2.13 ± 0.36,  n = 8 | 48.84 ± 2.96,  n = 8 |
| *elaV(C155)-Gal4/+; UAS-NDUFS7 [RNAi]/+;*  *IP3-Sponge.m30/+* | 0.67 ± 0.02,  n = 10 | 35.55 ± 1.53,  n = 10 | 3.70 ± 0.33,  n = 10 | 53.60 ± 3.17,  n = 10 |
| *elaV(C155)-Gal4/+; UAS-NDUFS7 [RNAi]/+;*  *UAS-IP_3_-Sponge.m30/+*  with Dantrolene (10µM) | 0.58 ± 0.02,  n = 10 | 33.07 ± 1.09,  n = 10 | 3.99 ± 0.50,  n = 10 | 57.65 ± 3.22,  n = 10 |
| *elaV(C155)-Gal4/+; UAS-mcu[RNAi]/+* | 0.77 ± 0.04,  n = 9 | 34.87 ± 1.24,  n = 9 | 3.39 ± 0.33,  n = 9 | 46.49 ± 3.16,  n = 9 |
| *elaV(C155)-Gal4/+; UAS-mcu[RNAi]/+*  with XestC (20µM) & Dantrolene (10µM) | 0.70 ± 0.08,  n = 9 | 38.58 ± 1.25,  n = 9 | 2.44± 0.21,  n = 9 | 59.15 ± 5.97,  n = 9 |
| *elaV(C155)-Gal4/+; UAS-NDUFS7 [RNAi]/*  *UAS-mcu[RNAi]* | 0.70 ± 0.05,  n = 9 | 32.26 ± 1.12,  n = 10 | 3.83 ± 0.46,  n = 9 | 48.87 ± 3.84,  n = 9 |
| *elaV(C155)-Gal4/+; UAS-NDUFS7 RNAi]/*  *UAS-mcu[RNAi]* with XestC (20µM)  & Dantrolene (10µM) | 0.55 ± 0.04,  n = 9 | 28.68 ± 1.63,  n = 11 | 3.18 ± 0.40,  n = 9 | 55.14 ± 7.01,  n = 9 |
| *elaV(C155)-Gal4/+; UAS-NDUFS7 [RNAi]/*  *UAS-mcu[RNAi]/IP_3_-Sponge.m30*  with Dantrolene (10µM) | 0.56 ± 0.05,  n = 8 | 24.96 ± 2.91,  n = 8 | 3.24 ± 0.42,  n = 8 | 46.83 ± 6.20,  n = 8 |
| *elaV(C155)-Gal4/+*  + DMSO and wash | 0.68 ± 0.01,  n = 12 | 40.92 ± 2.38,  n = 12 | 2.56 ± 0.35,  n = 12 | 60.68 ± 4.04,  n = 12 |
| *elaV(C155)-Gal4/+*  + BAPTA-AM (20µM) and wash | 0.66 ± 0.02,  n = 13 | 40.38 ± 1.68,  n = 13 | 2.21 ± 0.14,  n = 13 | 61.94 ± 3.92,  n = 13 |
| *elaV(C155)-Gal4/+;*  *UAS-NDUFS7 [RNAi]/+*  + DMSO and wash | 0.59 ± 0.02,  n = 14 | 43.39 ± 1.86,  n = 14 | 3.28 ± 0.24,  n = 14 | 74.81 ± 4.42,  n = 14 |
| *elaV(C155)-Gal4/+;*  *UAS-NDUFS7 [RNAi]/+*  + BAPTA-AM (20µM) and wash | 0.69 ± 0.06,  n = 11 | 31.65 ± 2.92,  n = 11 | 2.63 ± 0.43,  n = 11 | 49.79 ± 6.74,  n = 11 |

*elaV(C155)-Gal4* is a pan-neuronal driver. Values represent mean ± s.e.m.

**S15 Table, related to Figure S14:** Active zone marker intensity for genotypes in Figure S14.

| **Genotype** | **BRP Intensity in AU**  **(µm^2^ area of a bouton)** |
| --- | --- |
| *elaV(C155)-Gal4/*+DMSO | 100.0 ± 6.96, n = 67 |
| *elaV(C155)-Gal4/*+Xesto+Dant | 100.1 ± 6.22, n = 86 |
| *elaV(C155)-Gal4/+; NDUFS7 [RNAi]/+*DMSO | 170.4 ± 14.26, n = 100 |
| *elaV(C155)-Gal4/+; NDUFS7 [RNAi]/****+***Xesto+Dant | 142.7 ± 7.03, n = 100 |
| *elaV(C155)-Gal4/+; UAS-IP_3_-sponge m.30/*+ | 111.8 ± 6.02, n = 88 |
| *elaV(C155)-Gal4/+; mcu[RNAi]/+* | 100.3 ± 4.28, n = 111 |
| *elaV(C155)-Gal4/+; UAS-mcu[RNAi]/NDUFS7 [RNAi]*+Xesto+Dant | 64.78 ± 4.59, n = 83 |
| *elaV(C155)/+; UAS-mcu[RNAi]/NDUFS7 [RNAi]/*+;  *UAS-IP_3_-sponge.m30/+*Dant | 85.73 ± 3.70, n = 100 |

*elaV(c155)-Gal4* is a pan-neuronal driver. Xesto denotes XestosponginC. Dant denotes Dantrolene. Values represent mean ± s.e.m.

**S16 Table 16, related to Figure 6:** Electrophysiological properties and active zone marker intensity measurements for conditions depicted in Figure 6.

| **Genotype** | **mEPSP**  **amplitude**  **(mV)** | **mEPSP**  **frequency**  **(Hz)** | **EPSP**  **amplitude**  **(mV)** | **Quantal**  **content**  **(QC)** | **BRP Intensity**  **in AU**  **(µm^2^ area**  **of a bouton)** |
| --- | --- | --- | --- | --- | --- |
| *elaV(C155)-Gal4/+*  +DMSO | 0.60 ± 0.02,  n = 10 | 4.34 ± 0.74,  n = 10 | 41.52 ± 0.99,  n = 10 | 68.70 ± 2.29,  n = 10 | 100.0 ± 6.79,  n = 69 |
| *elaV(C155)-Gal4/+*  +LDA | 0.69 ± 0.04,  n = 9 | 4.93 ± 0.54,  n = 9 | 41.22 ± 1.29,  n = 10 | 60.59 ± 3.38,  n = 9 | 193.1 ± 10.41,  n = 94 |
| *elaV(C155)-Gal4/+*  *UAS-NDUFS7 [RNAi]/+*  +DMSO | 0.61 ± 0.03,  n = 9 | 2.79 ± 0.27,  n = 9 | 40.46 ± 0.82,  n = 9 | 66.97 ± 3.72,  n = 9 | 100.0 ± 4.65,  n = 75 |
| *elaV(C155)-Gal4/+*  *UAS-NDUFS7 [RNAi]/+*  +LDA | 0.58 ± 0.02,  n = 11 | 3.15± 0.25,  n = 11 | 35.71 ± 0.56,  n = 11 | 62.25 ± 2.95,  n = 11 | 81.05 ± 5.37,  n = 71 |
| *elaV(C155)-Gal4/+*  +2-Deoxy-D-glucose | 0.96 ± 0.04,  n = 9 | 1.94 ± 0.22,  n = 9 | 41.09 ± 1.66,  n = 9 | 42.70 ± 1.52,  n = 9 | 100.0 ± 4.71,  n = 74 |
| *elaV(C155)-Gal4/+*  *UAS-NDUFS7 [RNAi]/+*  +2-Deoxy-D-glucose | 0.66 ± 0.07,  n = 8 | 3.74 ± 1.02,  n = 8 | 32.27 ± 1.72,  n = 8 | 52.65 ± 5.82,  n = 8 | 78.69 ± 3.80,  n = 84 |

*elaV(C155)-Gal4* is a pan-neuronal driver. LDA denotes lonidamine. Values represent mean ± s.e.m.

**S17 Table, related to Figure S15:** Electrophysiological parameters for conditions depicted in Figure S15.

| **Genotype** | **mEPSP**  **amplitude**  **(mV)** | **mEPSP**  **frequency**  **(Hz)** | **EPSP**  **amplitude**  **(mV)** | **Quantal**  **Content**  **(QC)** |
| --- | --- | --- | --- | --- |
| *elaV(C155)-Gal4/+* | 0.70 ± 0.04,  n = 10 | 2.72 ± 0.52,  n = 10 | 42.40 ± 1.19,  n = 10 | 62.46 ± 3.71,  n = 10 |
| *elaV(C155)-Gal4/+;*  *UAS-NDUFS7 [RNAi]/+* | 0.97 ± 0.07,  n = 10 | 2.92 ± 0.46,  n = 10 | 42.11 ± 1.07,  n = 11 | 47.33 ± 5.26,  n = 10 |
| *elaV(C155)-Gal4/+;*  *UAS-hex A[RNAi]/+* | 0.62 ± 0.02,  n = 10 | 1.64 ± 0.17,  n = 10 | 32.70 ± 1.75,  n = 11 | 54.26 ± 3.85,  n = 10 |
| *elaV(C155)-Gal4/+;*  *UAS-hex C[RNAi]/+* | 0.67 ± 0.03,  n = 10 | 1.84 ± 0.21,  n = 10 | 44.69 ± 1.36,  n = 10 | 67.60 ± 4.48,  n = 10 |
| *elaV(C155)-Gal4/+; UAS-hex C[RNAi]/+; UAS-hex A[RNAi]/+* | 0.90 ± 0.04,  n = 11 | 1.23 ± 0.09,  n = 11 | 36.94 ± 3.09,  n = 13 | 42.02 ± 4.65,  n = 11 |
| *elaV(C155)-Gal4/+;*  *UAS-idh[RNAi]/+* | 0.71 ± 0.03,  n = 10 | 1.79 ± 0.27,  n = 10 | 48.58 ± 1.60,  n = 10 | 69.15 ± 3.67,  n = 10 |
| *elaV(C155)Gal4/+;*  *UAS-Cit.(Si)-Syn[RNAi]/+* | 0.87 ± 0.09,  n = 9 | 1.67 ± 0.14,  n = 9 | 44.61 ± 1.98,  n = 9 | 53.81 ± 3.20,  n = 9 |
| *elaV(C155)-Gal4/+;*  *UAS-Scsα[RNAi]/+* | 0.83 ± 0.04,  n = 8 | 1.82 ± 0.37,  n = 8 | 42.33 ± 2.81,  n = 9 | 50.19 ± 3.47,  n = 8 |
| *elaV(C155)-Gal4/+;*  *UAS-NDUFS7 [RNAi]/+;*  *UAS-hex A[RNAi]/+* | 0.62 ± 0.02,  n = 12 | 2.99 ± 0.33,  n = 12 | 31.10 ± 0.81,  n = 13 | 51.33 ± 2.43,  n = 12 |
| *elaV(C155)-Gal4/+;*  *UAS-NDUFS7 [RNAi]/+;*  *UAS-hex C[RNAi]/+* | 0.74 ± 0.04,  n = 12 | 3.39 ± 0.33,  n = 12 | 37.93 ± 1.37,  n = 12 | 53.38 ± 3.66,  n = 12 |
| *elaV(C155)-Gal4/+;*  *UAS-NDUFS7 RNAi]/UAS-hex C[RNAi]; UAS-hex A[RNAi]/+* | 0.99 ± 0.05,  n = 12 | 2.60 ± 0.48,  n = 12 | 27.70 ± 1.86,  n = 13 | 28.48 ± 2.10,  n = 12 |
| *elaV(C155)-Gal4/+;*  *UAS-NDUFS7 [RNAi]/+;*  *UAS-idh[RNAi]/+* | 0.62 ± 0.01,  n = 12 | 3.40 ± 0.49,  n = 12 | 41.34 ± 1.72,  n = 12 | 66.30 ± 2.58,  n = 12 |
| *elaV(C155)-Gal4/+;*  *UAS-NDUFS7 [RNAi]/+;*  *UAS-Cit.(Si)-Syn[RNAi]/+* | 0.68 ± 0.06,  n = 14 | 3.95 ± 0.72,  n = 14 | 32.73 ± 1.87,  n = 14 | 51.82 ± 4.85,  n = 14 |
| *elaV(C155)-Gal4/+;*  *UAS-NDUFS7 [RNAi]/+;*  *UAS-Scsα[RNAi]/+* | 0.74 ± 0.04,  n = 8 | 3.09 ± 0.66,  n = 8 | 29.90 ± 1.20,  n = 8 | 41.45 ± 3.17,  n = 8 |

*elaV(C155)-Gal4* is a pan-neuronal driver. Values represent mean ± s.e.m.

**S18 Table, related to Figure S10:** Active zone intensity data for genotypes in Figure S16.

| **Genotype** | **BRP Intensity in AU**  **(µm^2^ area of a bouton)** |
| --- | --- |
| *elaV(C155)-Gal4/+* | 100.0 ± 6.74, n = 68 |
| *elaV(C155)-Gal4/+; UAS-NDUFS7 [RNAi]/+* | 218.3 ± 10.89, n = 96 |
| *elaV(C155)-Gal4/+; UAS-hex A[RNAi]/+* | 209.6 ± 11.43, n = 69 |
| *elaV(C155)-Gal4/+; UAS-hex C[RNAi]/+* | 93.20 ± 7.22, n = 68 |
| *elaV(C155)-Gal4/+;*  *UAS-hex C[RNAi]/+; UAS-hex A[RNAi]/+* | 123.3 ± 5.68, n = 84 |
| *elaV(C155)-Gal4/+; UAS-idh[RNAi]/+* | 223.0 ± 14.31, n = 98 |
| *elaV(C155)-Gal4/+; UAS-Cit.(Si)-Syn[RNAi]/+* | 187.1 ± 12.40, n = 79 |
| *elaV(C155)-Gal4/+; UAS-Scsα[RNAi]/+* | 155.8 ± 11.47, n = 86 |
| *elaV(C155)-Gal4/+; UAS-NDUFS7 [RNAi]/+;*  *UAS-hex A[RNAi]/+* | 54.37 ± 4.41, n = 63 |
| *elaV(C155)-Gal4/+; UAS-NDUFS7 [RNAi]/UAS-hex C[RNAi]* | 121.7 ± 11.47, n = 68 |
| *elaV(C155)-Gal4/+; UAS-NDUFS7 [RNAi]/UAS-hex C[RNAi]/+; UAS-hex A[RNAi]/+* | 120.2 ± 6.61, n = 84 |
| *elaV(C155)-Gal4/+; UAS-NDUFS7 RNAi]/+;*  *UAS-idh[RNAi]/+* | 81.01 ± 4.54, n = 87 |
| *elaV(C155)-Gal4/+; UAS-NDUFS7 [RNAi]/+;*  *UAS-Cit.(Si)-Syn[RNAi]/+* | 138.2 ± 8.38, n = 90 |
| *elaV(C155)-Gal4/+; UAS-NDUFS7 [RNAi]/+;*  *UAS-Scsα[RNAi]/+* | 73.09 ± 5.49, n = 75 |

*elaV(C155)-Gal4* is a pan-neuronal driver. Values represent mean ± s.e.m.

**S19 Table, related to Figure 7:** NMJ developmental parameters for genotypes shown in Figure 7.

| **Genotype** | **# boutons** | **Muscle area (µm^2^)** | **# of branches** | **Relative Dlg**  **area (µm^2^)** | **Bouton area (µm^2^)** |
| --- | --- | --- | --- | --- | --- |
| *BG57-Gal4/+* | 94.38 ± 5.09,  n = 8 | 59050 ± 2845,  n = 8 | 12.00 ± 1.01,  n = 8 | 8.39 ± 0.70,  n = 30 | 19.20 ± 1.15,  n = 30 |
| *UAS-Sod2/+;*  *BG57-Gal4/+* | 83.38 ± 4.76,  n = 8 | 56340 ± 1447,  n = 8 | 8.12 ± 0.78,  n = 8 | 6.25 ± 0.38,  n = 30 | 18.11 ± 0.84,  n = 30 |
| *UAS-Sod1/+;*  *BG57-Gal4/+* | 83.89 ± 5.39,  n = 9 | 54490 ± 2001,  n = 9 | 7.55 ± 0.70,  n = 9 | 7.05 ± 0.69,  n = 29 | 22.44 ± 1.30,  n = 29 |
| *UAS-Cat/+;*  *BG57-Gal4/+* | 79.50 ± 5.80,  n = 8 | 56190 ± 2297,  n = 8 | 8.50 ± 0.50,  n = 8 | 9.60 ± 0.77,  n = 30 | 26.26 ± 1.16,  n = 30 |
| *UAS-NDUFS7 RNAi/+;*  *BG57-Gal4/+* | 61.88 ± 5.57,  n = 8 | 34250 ± 2546,  n = 8 | 7.00 ± 0.70,  n = 8 | 2.36 ± 0.23,  n = 30 | 10.40 ± 0.59,  n = 30 |
| *UAS-NDUFS7 [RNAi]/+;*  *BG57-Gal4/+*  *+*0.5mM NACA | 92.38 ± 5.09,  n = 8 | 35990 ± 1377,  n = 8 | 8.62 ± 0.56,  n = 8 | 5.63 ± 0.42,  n = 30 | 15.27 ± 0.93,  n = 30 |
| *UAS-NDUFS7 RNAi]/*  *UAS-Sod2;*  *BG57-Gal4/*+ | 87.88 ± 5.83,  n = 8 | 35850 ± 1967,  n = 8 | 9.00 ± 0.68,  n = 8 | 7.55 ± 0.43,  n = 34 | 15.59 ± 0.75,  n = 34 |
| *UAS-NDUFS7 [RNAi]/*  *UAS-Sod1;*  *BG57-Gal4/*+ | 58.75 ± 4.17,  n = 8 | 31620 ± 2084,  n = 8 | 5.75 ± 0.31,  n = 8 | 3.53 ± 0.50,  n = 30 | 14.36 ± 0.93,  n = 30 |
| *UAS-NDUFS7 [RNAi]/*  *UAS-Cat; BG57-Gal4/+* | 58.25 ± 3.80,  n = 8 | 31180 ± 1069,  n = 8 | 6.37 ± 0.26,  n = 8 | 3.91 ± 0.35,  n = 30 | 14.52 ± 0.73,  n = 30 |
| *elaV(C155)-Gal4/+* | 89.75 ± 5.00,  n = 8 | 70710 ± 4063,  n = 9 | 7.00 ± 0.37,  n = 8 | 5.05 ± 0.37,  n = 22 | 10.82 ± 0.78,  n = 22 |
| *elaV(C155)-Gal4/+;*  *UAS-NDUFS7 [RNAi]/+* | 113.8 ± 7.81,  n = 8 | 74980 ± 2586,  n = 8 | 9.25 ± 0.83,  n = 8 | 4.23 ± 0.45  n = 25 | 10.10 ± 0.62,  n = 25 |
| *D42-Gal4/+* | 98.22 ± 9.30,  n = 9 | 70850 ± 2785,  n = 9 | 7.88 ± 1.04,  n = 9 | 5.51 ± 0.40,  n = 25 | 8.59 ± 0.38,  n = 25 |
| *UAS-NDUFS7 [RNAi]/+;*  *D42-Gal4/+* | 121.4 ± 4.58,  n = 8 | 68390 ± 3436,  n = 8 | 9.87 ± 0.93,  n = 8 | 4.92 ± 0.45,  n = 25 | 7.95 ± 0.53,  n = 25 |

*BG57-Gal4* is a muscle driver. *elaV(C155)-Gal4* is a pan-neuronal driver. *D42-Gal4* is a motor neuron driver. Values represent mean ± s.e.m.

**S20 Table, related to Figure 9:** Electrophysiological parameters for genotypes and conditions in Figure 9.

| **Genotype** | **mEPSP**  **amplitude (mV)** | **mEPSP**  **frequency (Hz)** | **EPSP**  **amplitude (mV)** | **Quantal**  **Content (QC)** |
| --- | --- | --- | --- | --- |
| *BG57-Gal4/+* | 0.77 ± 0.02,  n = 8 | 4.54 ± 0.36,  n = 8 | 37.33 ± 1.60,  n = 8 | 48.63 ± 2.60,  n = 8 |
| *NDUFS7 [RNAi]/+;*  *BG57-Gal4/+* | 0.63 ± 0.05,  n = 7 | 5.45 ± 0.76,  n = 7 | 25.49 ± 1.31,  n = 7 | 41.97 ± 4.12,  n = 7 |
| *NDUFS7 [RNAi]/UAS-Cat;*  *BG57-Gal4/+* | 0.79 ± 0.06,  n = 8 | 9.27 ± 0.74,  n = 8 | 25.47 ± 2.20,  n = 8 | 33.07 ± 3.39,  n = 8 |
| *NDUFS7 [RNAi]/ UAS-Sod1*  *BG57-Gal4/*+ | 0.77 ± 0.07,  n = 8 | 5.28 ± 0.73,  n = 8 | 26.34 ± 1.82,  n = 8 | 37.15 ± 5.05,  n = 8 |
| *NDUFS7 [RNAi]/UAS-Sod2*  *BG57-Gal4/*+ | 0.77 ± 0.07,  n = 6 | 6.99 ± 1.21,  n = 6 | 33.26 ± 1.08,  n = 6 | 38.24 ± 6.79,  n = 6 |
| *BG57-Gal4/+*  +10% EtOH | 0.77 ± 0.04,  n = 9 | 4.79 ± 0.36,  n = 9 | 37.77 ± 2.00,  n = 9 | 50.17 ± 3.92,  n = 9 |
| *NDUFS7 [RNAi]/+;*  *BG57-Gal4/+* 10% EtOH | 0.62 ± 0.05,  n = 12 | 7.12 ± 1.30,  n = 12 | 27.49 ± 1.20,  n = 10 | 43.65 ± 4.06,  n = 10 |
| *NDUFS7 RNAi]/+;*  *BG57-Gal4/+*  + 0.5mM Curcumin | 0.69 ± 0.06,  n = 9 | 6.07 ± 0.85,  n = 9 | 29.74 ± 1.28,  n = 9 | 46.00 ± 5.05,  n = 9 |
| *NDUFS7 [RNAi]/+; BG57-Gal4/+*  0.5mM NACA | 0.88 ± 0.037,  n = 9 | 5.34 ± 0.88,  n = 9 | 35.12 ± 1.23,  n = 9 | 39.88 ± 1.37,  n = 9 |
| *ND-30* mutant  (*ND-30 ^epgy^/Df*) | 0.50 ± 0.02,  n = 10 | 6.74 ± 1.11,  n = 10 | 20.27 ± 1.72,  n = 10 | 41.22 ± 4.34,  n = 10 |
| *ND-30 ^epgy^/Df*  *+* 0.5 mM NACA | 0.71 ± 0.06,  n = 8 | 6.50 ± 1.17,  n = 8 | 30.32 ± 1.36,  n = 8 | 44.59 ± 4.20,  n = 8 |
| *Sod2* muscle rescue  *UAS-Sod2/+;*  *ND-30 ^epgy^/Df, BG57-Gal4* | 0.56 ± 0.04,  n = 8 | 5.07 ± 0.34,  n = 8 | 36.19 ± 1.08,  n = 8 | 66.13 ± 4.42,  n = 8 |
| *BG57-Gal4/+*  + 0.5% DMSO | 0.92 ± 0.03,  n = 9 | 4.43 ± 0.44,  n = 9 | 40.09 ± 1.95,  n = 9 | 43.04 ± 0.79,  n = 9 |
| *BG57-Gal4/+*  + 50µM rotenone | 0.94 ± 0.08,  n = 9 | 4.36 ± 0.71,  n = 9 | 31.96 ± 1.43,  n = 9 | 35.50 ± 3.08,  n = 9 |
| *UAS-Cat/+; BG57-Gal4/+*  + 50µM rotenone | 0.72 ± 0.03,  n = 11 | 5.10 ± 0.52,  n = 11 | 31.58 ± 2.40,  n = 11 | 44.34 ± 3.67,  n = 11 |
| *UAS-Sod1/+; BG57-Gal4/+*  + 50µM rotenone | 0.82 ± 0.03,  n = 8 | 4.53 ± 0.46,  n = 8 | 30.07 ± 1.86,  n = 8 | 36.63 ± 2.30,  n = 8 |
| *UAS-Sod2/+; BG57-Gal4/+*  + 50µM rotenone | 0.71 ± 0.05,  n = 7 | 5.29 ± 0.49,  n = 7 | 36.54 ± 1.55,  n = 8 | 54.15 ± 4.60,  n = 7 |

*BG57-Gal4* is a muscle driver. Values represent mean ± s.e.m.

**S21 Table, related to Figure 9:** Crawling behavior of various MCI loss or rescue combinations.

| **Genotype** | **Distance crawled in cm** |
| --- | --- |
| *BG57-Gal4/+* | 2.55 ± 0.09, n = 12 |
| *UAS-Sod2/+; BG57-Gal4/+* | 2.72 ± 0.18, n = 15 |
| *UAS-ND-30[RNAi]/BG57-Gal4* | 1.83 ± 0.12, n = 15 |
| *UAS-NDUFS7 [RNAi]/+; BG57-Gal4/+* | 0.79 ± 0.06, n = 10 |
| *UAS-NDUFS7 [RNAi]/UAS-Sod2; BG57-Gal4/BG57-Gal4* | 1.49 ± 0.15, n = 12 |
| *UAS-NDUFS7 [RNAi]/+; BG57Gal4/+*  *+*NACA | 1.60 ± 0.13, n = 8 |
| *elaV(C155)-Gal4/+* | 2.47 ± 0.14, n = 10 |
| *elaV(C155)-Gal4/+ ;NDUFS7 [RNAi]/+* | 2.51 ± 0.14, n = 10 |
| *ND-30^epgy^/Df* | 1.34 ± 0.15, n = 9 |
| *UAS-Sod2/+;ND-30^epgy^/Df,BG57-Gal4* | 1.85 ± 0.15, n = 9 |

*BG57-Gal4* is a muscle driver*. elaV(C155)-Gal4* is a pan neuronal driver. *Df* denotes the chromosomal deficiency *Df(3L)ED4288*, which uncovers the *ND-30* endogenous locus. Values represent mean ± s.e.m.

**S22 Table, related to STAR Methods section:** Reagents.

| ***Drosophila* gene name and Bloomington *Drosophila* stocks (stock number)** | **Antibodies (dilution), Drugs, and Fluorophores** |
| --- | --- |
| *UAS-ND-30[RNAi]/CG12079* (BL 44535) | Mouse α-BRP (DSHB-1:30),  Mouse α-DLG (DSHB-1:50) |
| *UAS-NDUFS7 [RNAi]/CG2014* (BL 62381) | Mouse α-Alpha Spectrin (DSHB-1:20) |
| *UAS-marf [RNAi]* (BL 67158),  *UAS-drp1[RNAi]* (BL 51483) | Mouse α-CSP (DSHB-1:50),  Mouse α-Synapsin (DSHB-1:30) |
| *UAS-mitoGFP* (BL 8442),  *UAS-mitoGFP, D42* (BL 42737) | Mouse α-GluRIIA (DSHB-1:50) |
| *UAS-Sod1* (BL 33605),  *UAS-Sod2* (BL 24494),  *UAS-Catalase* (BL 24621) | Mouse α-22C10 (DSHB-1:50) |
| *Df(3L)ED4288* (BL 8057),  *ND-30^epgy^* = *ND-30^EY03664^* (BL 16569) | Polyclonal rabbit α-GFP (Abcam: 1:250)  Polyclonal rabbit α-GluRIII  (Dr. Aaron DiAntonio lab-1:100)  Polyclonal rabbit α-DLG (1:700) |
| *elaV(C155-Gal4)* (BL 458),  *D42-Gal4* (BL 8816) | Dantrolene (Tocris: 10 µM),  Xestospongin C (Abcam: 20 µM)  Rotenone (Sigma aldrich: 50 µM),  Curcumin (Sigma aldrich: 0.5 mM) |
| *BG57-Gal4* (BL 32556) | NACA (Sigma Aldrich: 0.5mM), MitoSOX™ Red (Molecular Probes,  Thermo Fisher Scientific, 1:200),  BAPTA-AM (Sigma Aldrich)  Lonidamine (LDA: 150 µM Sigma Aldrich)  2-Deoxy-D-glucose (Sigma Aldrich) |
| *UAS-CalX [RNAi]* (BL 28306),  *UAS-mcu[RNAi]* (BL 42580, BL 67857),  *UAS-IP3-Sponge.m30* (Koganezawa lab)  *UAS-hexokinase A[RNAi]* (BL 35155)  *UAS-hexokinase C[RNAi]* (BL 57404)  *UAS-idh[RNAi]* (BL 41708)  *UAS-Cit.(Si)-Syn[RNAi]* (BL 36740)  *UAS-Scsα[RNAi]* (BL 51807)  *UAS-Sod2[RNAi]* (BL 24489) | Alexa α-HRP 488 (1:800), Rhodamine α-HRP (1:200),  Alexa α-HRP 647 (1:200),  Mouse and Rabbit Alexa Fluor 488 or 568 (1:400) |
